# Model-based tumor subclonal reconstruction

**DOI:** 10.1101/586560

**Authors:** Giulio Caravagna, Timon Heide, Marc Williams, Luis Zapata, Daniel Nichol, Ketevan Chkhaidze, William Cross, George D. Cresswell, Benjamin Werner, Ahmet Acar, Chris P. Barnes, Guido Sanguinetti, Trevor A. Graham, Andrea Sottoriva

## Abstract

The vast majority of cancer next-generation sequencing data consist of bulk samples composed of mixtures of cancer and normal cells. To study tumor evolution, subclonal reconstruction approaches based on machine learning are used to separate subpopulation of cancer cells and reconstruct their ancestral relationships. However, current approaches are entirely data-driven and agnostic to evolutionary theory. We demonstrate that systematic errors occur in subclonal reconstruction if tumor evolution is not accounted for, and that those errors increase when multiple samples are taken from the same tumor. To address this issue, we present a novel approach for model-based subclonal reconstruction that combines data-driven machine learning with evolutionary theory. Using public, synthetic and newly generated data, we show the method is more robust and accurate than current techniques in both single-sample and multi-region sequencing data. With careful data curation and interpretation, we show how the method allows minimizing the confounding factors that affect non-evolutionary methods, leading to a more accurate recovery of the evolutionary history of human tumors.

## Introduction

Cancers evolve through a process of clonal evolution^1^, inevitably resulting in intra-tumor heterogeneity^2^. Genome sequencing of one or more bulk samples from tumors has become the most common way to study intra-tumor heterogeneity and clonal evolution in human malignancies. Concerted efforts are dedicated to the identification of cancer (sub)clones^3^. Whereas a cancer “clone” remains loosely defined, its purest definition is that of a group of cells within the tumor that share a common ancestor. However, this implies that any ancestor in the phylogenetic tree of a tumor can be identified as the founder of a distinct “clone”. For a cancer composed of *N* cells, by this definition we would expect *N−1* clones, most of which have nothing ‘special’ in terms of tumor biology. This is why in the field we implicitly identify clones “of interest”, such as those that have growth/survival advantage (an ancestor under positive selection), or those that generate metastases (an ancestor that first arrives at a given site). It is important to bear in mind these limits in the definition of a clone when we attempt to recovery the tumor clonal architecture.

Unsupervised clustering of variant read counts from bulk cancer sequencing data is the established approach to resolve the clonal structure of a bulk tumor sample^4^, with each of the resulting clusters defined as a clone. This procedure, called “subclonal reconstruction” (or deconvolution), leverages on variant read counts and associated variant allele frequency (VAF) estimates of somatic mutations, which are normalized for copy number status (number of alleles at the mutation locus) and tumor purity (proportion of contaminating normal alleles). This normalization leads to the computation of the Cancer Cell Fraction (CCF), or the proportion of cancer cells bearing a given mutation. Subclonal reconstruction is central to cancer evolution analyses because it allows tracing the “life history” of a tumor by reconstructing the underlying clonal ancestral relationships in the form of a clone phylogenetic tree (called a “clone tree”)^3^.

Current methodologies approach subclonal reconstruction with sophisticated machine learning or combinatorial optimization algorithms^4^. The former class of algorithms is predominant, and a large set of methods use Dirichlet Processes clustering^3,5,6^ or Dirichlet finite mixture models^7^. These methods are entirely data-driven and are usually chosen because of their convenient statistical properties, rather than their adherence to the mechanisms of tumor evolution and sampling. Nevertheless, they can be efficient and accurate, as long as the underlying assumptions of the statistical method are correct. All current subclonal reconstruction methods assume that variant read counts from bulk tumor samples composed of different subclonal populations would present as a mixture of Binomial or Beta-Binomial mutational clusters, each corresponding to a clone. Biologically, each of these clusters must consist of the mutations present in the founder cell of the clone^8^. However, these clusters are not the only observable patterns in the data: the mutations that occur within each clone (intra-clone mutations) are also detectable in the data. Given the size of the human genome, even with extremely low mutations rates such as germline mutation rates (e.g. 10^−9^ nucleotide substitutions per base per division^9^), new mutations are expected at each cell division, and thus large numbers of “passenger” mutations inevitably accumulate within a clone. The evolutionary dynamics of this passenger mutation accumulation are neutral, giving rise to a power-law distributed “tail” of ever more mutations at ever lower frequency within the clone. This has been mathematically demonstrated in theoretical population genetics^10–13^ and is corroborated by genomic data at high resolution^8,14^. These within-clone neutral tails have not been directly addressed by previous methods, potentially confounding the measurement of clonal heterogeneity through the introduction of spurious subclones.

Here, we aimed to reconcile data-driven machine learning approaches to clustering VAFs with the insight given by theoretical models of tumor evolution. Specifically, we combined Dirichlet mixture models with the set of distributions predicted by theoretical population genetics models^10–13^, producing the first model-based unsupervised clustering method for subclonal reconstruction called `MOBSTER` (MOdel Based cluSTering in cancER). `MOBSTER` can process mutant allelic frequencies to identify and remove neutral tails from the data, so that any subclonal reconstruction algorithm can be applied to determine subclone parameters from read counts.

We also expand `MOBSTER` to analyze multivariate data from multiple samples of the same tumor. We show that unavoidable sampling bias and lineage admixture caused by the spatial structure of the tumor^15^ produces additional confounding factors that need to be considered when interpreting the output of the subclonal reconstruction. Rational curation based on understanding how a tumor expands, how its growth is affected by stochastic forces such as drift^16^, and a careful spatial sampling strategy, can be combined with `MOBSTER` to accurately reconstruct the tumor phylogenetic history.

## Results

### Mutation, drift and selection in clonally evolving cancer cell populations

Cancers grow from a single cell, and because of this growth process, neutral mutations that occur in the first few cell divisions are present at high frequency in the final cancer, irrespective of the action of selection. In addition, stochastic fluctuations in population size of cell lineages can also increase the frequency of mutations in the absence of selection, this is called genetic drift^16^. The same is true within (sub)clones: a clone originates as a single cell, and neutral mutations that occur early within the clone thereafter are carried to higher frequency by the clone’s own growth.

Tremendous insight into the accumulation of mutations in the absence of positive selection has come from the study of the Luria-Delbruck model in bacteria^17^. This has led to well-established population genetics theory describing the accumulation of mutations within neutrally growing populations^11,12^. The same theory applies to cancer clones^10,13^ and can be extended to model positively selected mutations in growing populations^8^. The theory states that we should expect a tail of neutral passenger mutations within a clone (Figure 1A). Neutral tails have only recently become evident in the data with the adoption of high-depth whole genome sequencing: lower depth sequencing (e.g. < 60x whole-genome sequencing – WGS^8^) is not sufficient to detect a tail, and exome or panel sequencing often assay too few mutations to show a clear VAF spectrum^18^.

**Figure 1.**
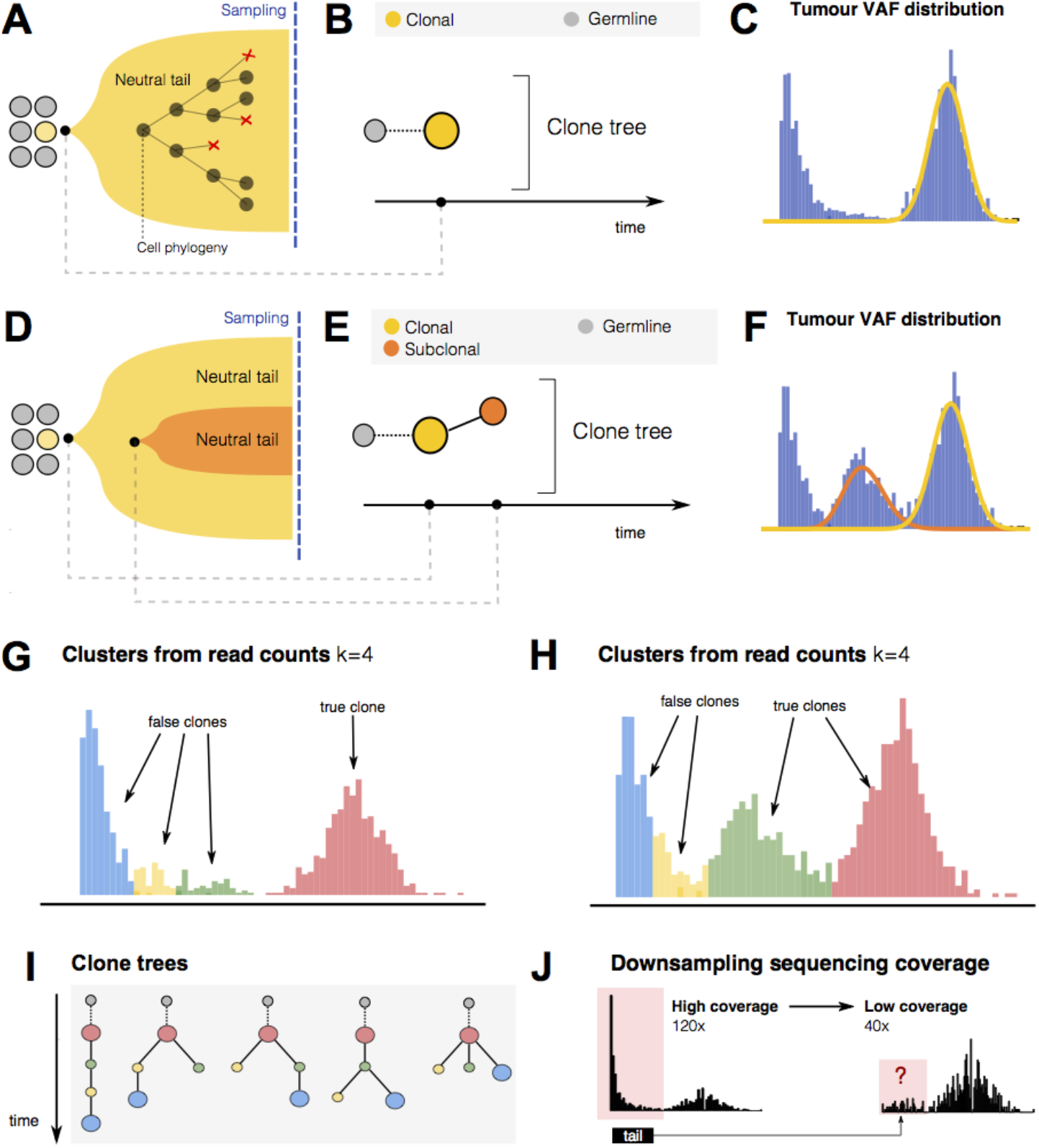
Cancer evolutionary dynamics with and without selection. (**A**) A tumor formed by a single clone expands following neutral evolutionary dynamics driven only by division, mutation, cell death and genetic drift. (**B**) The clone tree is represented as a single “truncal” clone. (**C**) The expected Variant Allele Frequency (VAF) distribution is characterized by a clonal cluster containing truncal mutations and a neutral 1/*f*^2^ tail. (**D**) A tumor with one subclone under positive selection. The evolutionary forces of mutation and neutral drift are still at play within each clone. (**E**) The clone tree is represented as a truncal clone that gives rise to a selected subclone within it. (**F**) Here the VAF histogram has one extra cluster due to subclonal mutations in the ancestor of the subclone that has risen in frequency due to positive selection. (**G,H**) Standard subclonal deconvolution from read counts of the tumors in panel A and B finds 4 clusters, multiple of which are not true subclones but are the result of clustering neutral tail mutations. (**I**) Inflated estimates of the number of tumor clones propagate errors and uncertainty in downstream evolutionary analysis. In this case, for instance, with 4 estimated clones we can fit several alternative phylogenetic trees (clone trees) to the clonal structure. The inferred clone trees will contain non-existing clones as nodes, portraying wrong evolutionary tumor histories. (**J**) Low coverage and low purity affect the ability to observe neutral tails in the data. In this synthetic example, we show how the VAF distribution of a neutral tumor with a clear neutral tail at 120x whole-genome sequencing can become difficult to interpret at 40x depth of sequencing. In this example, the degenerated tail at 40x may be interpreted as an actual subclone. With such data one is usually not powered to screen off true positive subclonal selection from actual neutral mutations.

We explored subclonal reconstruction in data containing neutral tails. Figure 1A shows the simplest example of a single uniform tumor expansion (i.e. no subclones). The corresponding clone tree has a single “truncal” node (Figure 1B). The VAF spectrum for this tumor consists of a clonal cluster at high frequency, corresponding to the mutations that are present in all cells in the sample (i.e., in the most recent common ancestor, MRCA, of the clone), and a neutral tail of mutations at low VAF originated as the clone expands (Figure 1C). In the case where a subclone with selective advantage is present (Figure 1D, E), the data will present as two clusters at high frequency (one clonal and one subclonal) as well as a mixture of two overlapping neutral tails (Figure 1F) – as previously reported^8^. Performing subclonal reconstruction on these data assuming a generative mixture of just Binomial or Beta-Binomial distributions will detect several clusters within the neutral tail that are erroneously identified as subclones (Figure 1G,H). When these clones are used downstream for phylogenetic reconstruction^4^, the resulting trees (Figure 1I) have a very different structure from the true trees (Figure 1B, E), with further high uncertainty for the fits because multiple possible trees can be consistent with the data.

Low depth sequencing data presents additional problems. In lower depth sequencing, neutral tails are undersampled and become even more likely to be mistaken for subclones as they lose their characteristic power-law shape. Comparison of a neutral tumor at 120x whole-genome to a 40x whole-genome, which shows how a “leftover” chunk of a neutral tail is indistinguishable from a subclonal cluster (Figure 1J). We note that these noisy subclonal VAF distribution that may represent under-sampled tails are commonly observed in low depth sequencing data previously reported^3,19^.

### Model-based clustering of variant allelic frequencies

The frequency *f* of passenger mutations in an expanding population follows a Landau distribution^11^, which at the frequency range detected by current sequencing standards can be approximated by a power law distribution *X~1/f*^2^ (Figure 2A), as we previously reported^10^. Subclonal alleles under positive selection, together with their hitchhiking passengers, will instead form clusters in the clone-size distribution as they rise in frequency due to Darwinian selection^20^.

**Figure 2.**
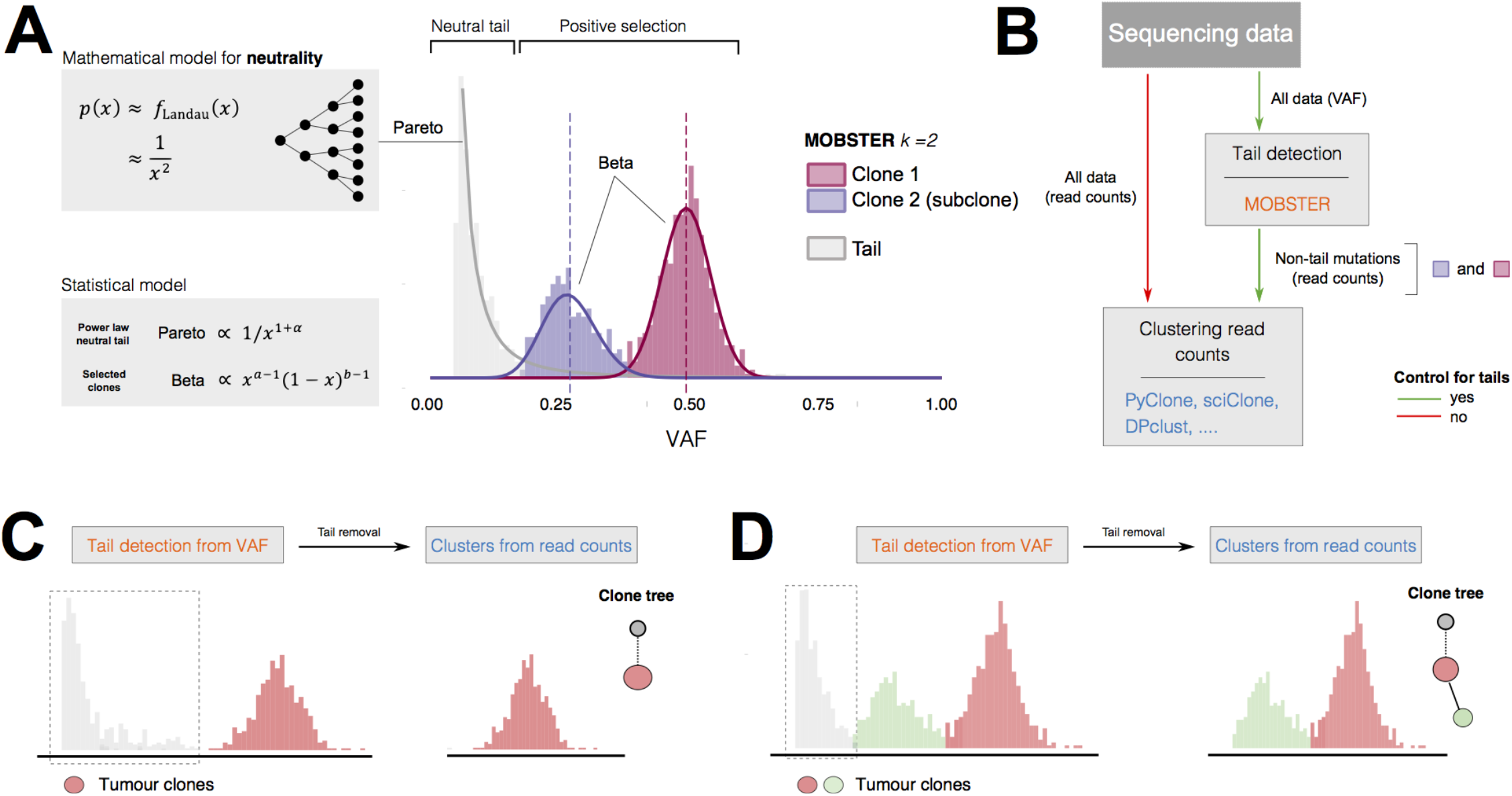
Model-based subclonal deconvolution. (**A**) `MOBSTER` combines a Pareto Type-I distribution (power law) with *k* Beta random variables, in a univariate finite mixture with *k* + 1 components. The Pareto captures the frequency spectrum of neutral mutant alleles (passengers), and Beta components detect alleles under positive selection. The Pareto distribution is predicted by standard theoretical population genetics, which shows that the probability of observing *x* mutant alleles in a neutral population follows a Landau distribution *f*_Landau_ that decays as 1/*x*^2^. `MOBSTER` uses a general formulation for a unique overall power law (with free parameter its exponent), and learns the model parameters from data via Expectation Maximization. In the histogram we show clustering assignments for a tumor with one subclone (*k* = 2, 1 subclone). (**B**) With `MOBSTER` we can control the confounding factor of neutral tails: we filter tail mutations from the VAF distribution, and then cluster remaining read counts with any other tool for subclonal reconstruction that uses read counts. Clusters detected in this way are more likely to accurately identify groups of alleles under positive selection (i.e., true clones). (**C, D**) In these synthetic examples, when we analyze tumors such as the ones in Figures 1A and 1B and control for tails, the number of clusters matches the number of true clones. The true clonal architectures and evolutionary histories of both tumors are therefore recovered.

Here, we consider VAF corrected for copy number status and tumor purity, with the expected clonal peak of a diploid tumor to be located at VAF ≈ 0.5 (which corresponds to a 100% Cancer Cell Fraction – CCF). We can model VAFs via Beta distributions^21^, and read counts with Binomial or Beta-Binomial distributions, depending on sequencing over-dispersion^3,5,6,21^. In `MOBSTER` we model the evolutionary dynamics of a growing tumor containing subclones by combining Beta distributions (expected from subclones under selection) with a power law (expected from the neutral tails of each subclone), and cluster VAF values (Figure 2A). After fitting the tail structure in the VAF distribution, the tail mutations – as they do not correspond to clones – can be removed, and clustering of read counts of remaining data (non-tail mutations) can be done via standard methods (Figure 2B). This analysis uses `MOBSTER` to control for tails while retaining the original variance of the data for the final clustering step of read counts. Critically, `MOBSTER` always compares in an unbiased way the fit of a mixture of subclones augmented with a neutral tail, versus the fit of a mixture of subclones alone. A regularized model selection strategy is then used to determine the best model fit to data.

`MOBSTER` combines one Pareto Type-I random variable (a type of power-law) with *k* Beta random variables; the overall model is a univariate finite mixture with *k + 1* components. The likelihood is

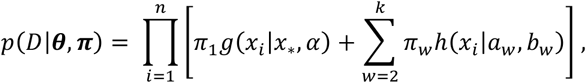

where *g* and *h* are density functions, ***θ*** = (*x*_*_, *α, a*_1_,.., *a_k_, b*_1_,…,*b_k_*) is a vector of parameters and ***π*** are mixing proportions in a standard setting with *N*×(*k* + 1) latent variables. The Pareto component follows *g*(*x|x*_*_, *α*) ∝ 1/*x*^1+*α*^ for *x* ≥ *x*_*_, and the Beta component follows *h*(*x|a_w_, b_w_*) ∝ *x*^*a_w_*−1^(1 − *x*)^*b_w_*−1^ in [0,1]. A detailed derivation of `MOBSTER`, its relation to other models in the field, and other technical comments are available (Online Methods, Supplementary Figure S1).

We can learn the parameters (*π,θ*) of a `MOBSTER` model via Expectation Maximization with numerical maximum likelihood estimates of the Beta parameters. Otherwise, we can adopt a much faster variation that exploits analytical moment-matching for Beta mixtures. To perform model selection and identify a parsimonious number of clusters (*k* ≥ 1), we minimize a score derived from the Integrative Classification Likelihood (`ICL`). `ICL` is an extension of the Bayesian Information Criterion (BIC) that includes the entropy of the latent variables to induce separation of the mixture components. In `MOBSTER` we either require a fit that separates all mixture components (subclones plus tail, standard `ICL`), or one that reduces the overlap among subclonal peaks, disregarding the tail (reduced `ICL`, re`ICL`)^22^.

`MOBSTER` can also fit models without tail (Beta mixtures), so it can choose the best model, with *vs* without tail, through a likelihood ratio test (Supplementary Figures S2 and S3). After tail detection and removal, any subclonal reconstruction tool can be used.

In the example of a tumor formed by a single expanding clone (Figure 1A), `MOBSTER` fits one Beta cluster of truncal mutations (clonal mutations present in all cancer cells) plus a neutral tail (Figure 2C). Similarly, for the tumor with one subclone (Figure 1D), `MOBSTER` fits two Beta clusters and a tail (Figure 2D). When `MOBSTER` removes tails, subsequent clustering of read counts for non-tail mutations identifies the true clones and their clone trees (Figure 2C-D).

### Synthetic validation of the method and confounding factors in univariate analysis

We used synthetic data to validate `MOBSTER` and to quantify the degree to which neutral tails confound subclonal deconvolution (see Online Methods, Supplementary Note 1 and Supplementary Data “Visualizing subclonal expansions”, for details). First, we generated *n* = 150 tumors evolved with plausible evolutionary parameters using a stochastic branching process simulator of tumor growth^8^. Out of these, 30 cases had no subclone (as in Figure 1A) and 120 one subclone (as in Figure 1D). For each tumor we simulated whole-genome bulk sequencing of a single biopsy, at high coverage (120x median, Poisson distributed) and perfect purity (*p* = 1; clonal cluster at adjusted VAF 0.5). We measured the predicted number of clones *k* (including the clonal cluster), as well as the distance between the true and the estimated peaks in the VAF distribution as a proxy for fit precision (Supplementary Figure S4). In all tests, we always compare `MOBSTER`’s fits with and without a tail, and retain the best fit (log ratio test) to compute performance. We carry out several other types of tests in order to assess the best model selection strategies, the ability of `MOBSTER` to detect low-frequency subclones, and its ability to distinguish tails from subclones. Adapting the computations by Williams *et al*.^8^, we also show that several evolutionary parameters of a tumor can be retrieved from `MOBSTER`’s output; these include the tumor mutation rate, the time of emergence of detected subclones and their selective advantage coefficient. Thus, `MOBSTER` is a clustering-based method for subclonal deconvolution that can also extract evolutionary dynamics information from the data (Supplementary Figures S4-S7).

By accounting for neutral tails, `MOBSTER` significantly outperformed standard approaches. This is observed with both Dirichlet variational mixture and a Dirichlet Process clustering run downstream `MOBSTER` (Figure 3A, B and Supplementary Figure S8); these statistical frameworks are at the core of popular tools like sciClone^7^, pyClone^23^ and DPclust^3^. In both methodologies, a parameter *α* > 0 tunes the propensity to introduce new clusters during the fit^3^; we tested both point estimates and a Gamma prior for *α*. Results confirm the importance of using stringent values of *α* (Figure 3A, B). Controlling for neutral tails with `MOBSTER` allowed retrieving the correct number of clones in the large majority of cases. Without `MOBSTER`, clone numbers were overestimated by up to a factor 4, with consistent errors. Importantly, stringent control of the clustering parameter *α* was insufficient to remove the errors, whereas the errors were avoided by explicitly correcting for neutral tails.

**Figure 3.**
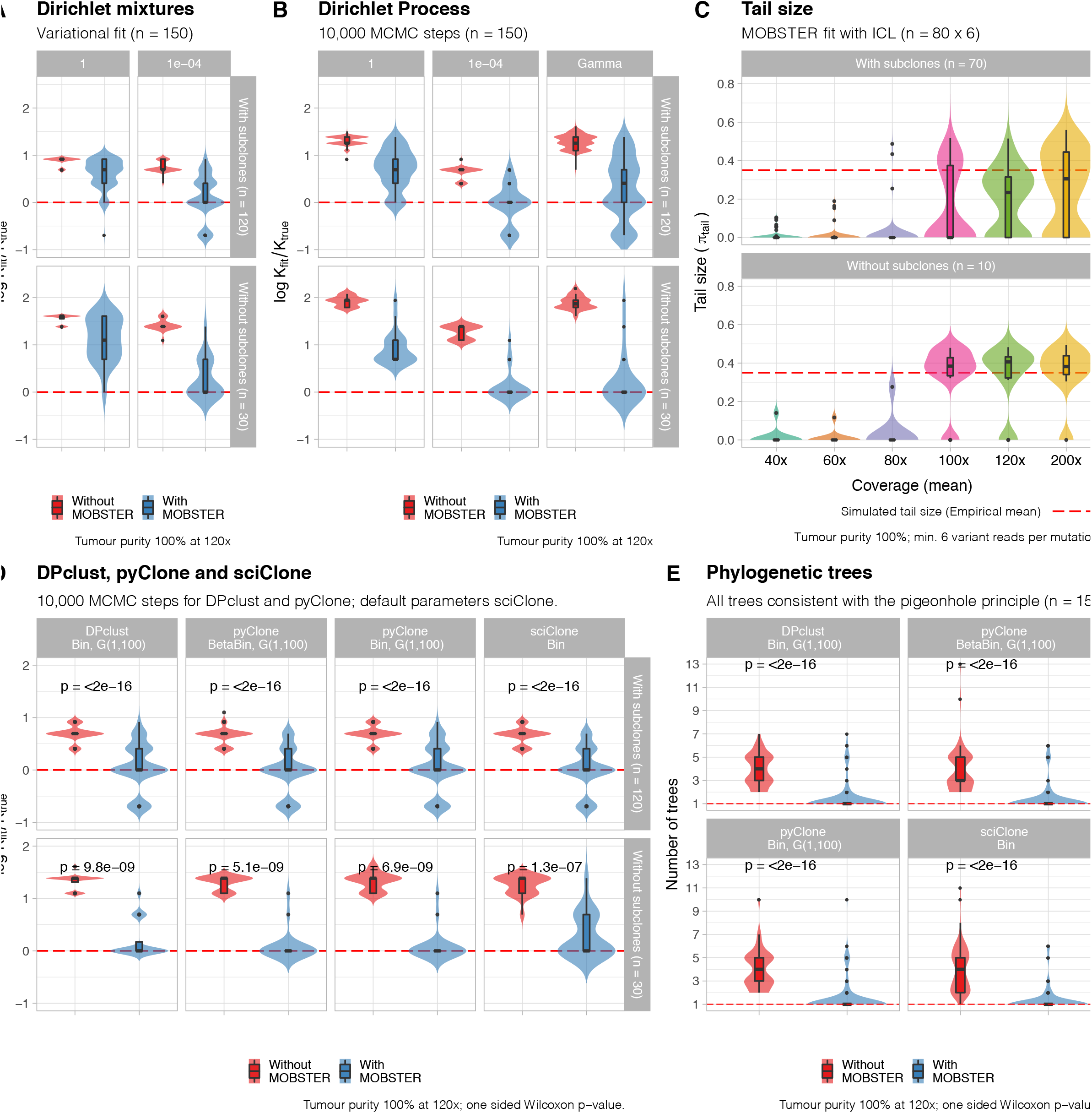
Robustness and accuracy of MOBSTER versus current approaches. (**A,B**) We compare subclonal reconstruction that controls for tail with `MOBSTER`, against standard methods. We use a variational fit to a Dirichlet mixture (**A**) and a Markov Chain Monte Carlo (MCMC) method for a Dirichlet Process (**B**). These methodologies are at the basis of standard approaches like sciClone, pyClone and DPClust. The test uses synthetic data of *n* = 150 simulated tumors (*n* = 120 with one subclone, and *n* = 30 without subclones), generated from a stochastic branching process of tumor growth. We report the logarithm of the ratio between the number of subclones found by the fit (*k*_fit_) and the true number of clones (*k*_true_); *k*_fit_=*k*_true_ when the ratio approaches 0. We assumed 120x mean coverage and 100% tumor purity (i.e., no normal contamination), and tested the performance of the clustering using different values of the mixture concentration parameter *α*, which tunes the propensity at which new cluster are called during the fit. Values *α* = 1,10^−4^ are point estimates of the concentration, and we test also a Dirichlet Process where *α* is learnt from the data using a Gamma prior. In these tests we use `ICL` for model selection; more extensive tests are available in Supplementary Figures S4-S8. (**C**) We measure how the proportion of mutations assigned to `MOBSTER`’s tails changes with coverage, assuming fixed 100% tumor purity. We span coverage values from 40x to 200x, using a subset of *n* = 80 tumors from the test in panel A/B. The red dashed line is the median tail size across the test set (ground truth obtained from simulated tumor). It is very difficult to fit neutral tails with below 100x coverage; a similar effect is observed reducing tumor sample purity (Supplementary Figures S5). (**D**) Using the same *n* = 150 synthetic tumors we tested a set of popular tools for subclonal deconvolution that exploit the methodologies shown in panel A/B. We compare Binomial mixtures from DPclust, pyClone and sciClone, and Beta-Binomial mixtures from pyClone. Compared to panels A/B, these tools can implement heuristics to merge clusters, or assign mutations to clusters. In this test, to avoid an excess of small subclones, we also remove output clusters with less than 2% of the overall mutational burden (*π* < 0.02). In this test we use a Gamma prior for *α* in DPclust and pyClone (default), with prior hyper-parameter (shape) set to 1 (value hardcoded in DPclust). sciClone uses a hardcoded *α* parameter. The error without pre-filtering with `MOBSTER` is dramatic in all cases. (**E**) Number of phylogenetic trees that can be fit by the pigeonhole principle using the output median VAF of each cluster. This number measures the reduction of the uncertainty in a downstream evolutionary analysis when we control for tails with `MOBSTER`.

We tested for the effect of two key confounders for tail detection: coverage (Figure 3C) and purity (Supplementary Figure S5). At 100% tumor purity (no contamination of non-tumor cells) tails could be reliably identified only if the median coverage exceeded 100x; progressively higher coverage was required with lower purity, as one might expect (e.g., at 120x median coverage purity >80% is required to detect tails consistently). Consequently, moderate depth whole-genome sequencing studies^19^ do not seem sufficiently powered to detect neutral tails, or to properly distinguish neutral tails from true subclones, as illustrated in Figure 1L.

In Figure 3D we explicitly compared DPclust, pyClone (with both Binomial and Beta-Binomial distributions) and sciClone, in combination with and without `MOBSTER`. Without `MOBSTER`, observed error rates for inferred clone number were comparable to results for baseline Dirichlet methods (Figure 3A-B). The impact on downstream phylogenetic analysis was severe. In particular, the inflated number of clusters detected when we do not control for tails generates a large number of trees that can be fit to data using the standard pigeonhole principle^3,24^ (Figure 3E).

### Confounding factors in subclonal analysis of multi-region sequencing data

We next investigated the influence of neutral tails on subclonal deconvolution from multi-region sequencing data. As for the single-sample data, our ultimate goal is to delineate the “important” clones from any other ancestor in the phylogenetic tree of a tumor, with specific interest in detecting clones that have experienced positive selection. In the single sample case, we have theoretical understanding of the clone size distribution under neutral evolution (clone size ~ 1/*f*^2^) and so selection is evident as a deviation from this expectation. Uncertainty over the spatial structure of a tumor (e.g. the spread and mixing of clones during tumor growth), coupled with the constraints and potential sampling-bias inherent to multi-region biopsies (e.g. variable number and physical sample size, non-uniform spatial sampling, heterogeneity in tumor content across tumor tissue, etc) preclude an analogous simple theoretical understanding of the clone size distribution in spatial multi-region data. Nonetheless, in `MOBSTER` we fit a general power law tail – rather than assuming its exponent to be equal to 2 – which gives us flexibility to take into account deviations from strict exponential growth due to spatial structure, as well as sampling bias.

In a sequencing dataset of multiple spatially-distinct samples from a tumor, clone identification is performed by identifying groups of mutations that are at the same VAF/ CCF in one sample (e.g. represent a cluster in single-region sequencing data) and which remain clustered together in other samples from the same patient^25^. Differences in the position of the cluster between samples represents differences in the abundance of the clone between samples^25,26^.

We generated synthetic data from spatial simulations of tumor growth under neutrality and positive selection (Online Methods) and observed that neutral tails in multi-region sequencing can lead standard methods to identify subclones from neutrally evolving ancestors, giving the illusion of numerous (false) selected clones. Strikingly, this error grows dramatically with the number of biopsies collected (number of regions). This means that, contrary to intuition, more samples generate proportionally more noise than signal for the subclonal reconstruction. This unpleasant effect is generated by different sources of spatial sampling bias, and the complex way in which passenger mutations from neutral tails spread in physical space within a growing neoplasm. We have isolated and identified these confounders, which we called the “hitchhiker mirage”, the “ancestor effect” and the “admixing deception” (Figure 4). The first confounder (hitchhiker mirage) is due to neutral tails, and can be formally solved using `MOBSTER`. The others are due to spatial sampling, and require careful interpretation of the results, which we discuss below.

**Figure 4.**
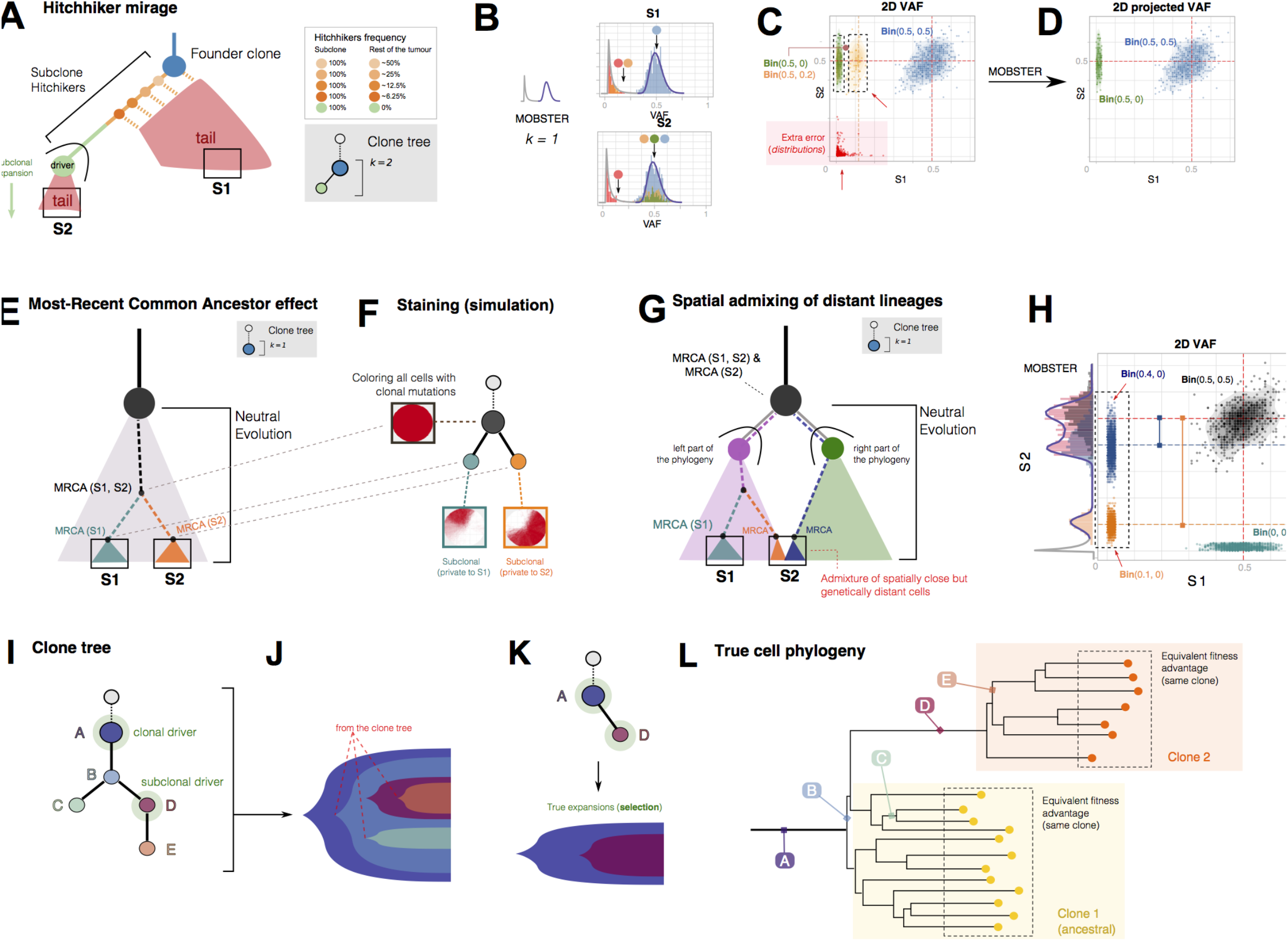
Confounding factors in multi-region bulk sequencing. (**A**) Evolutionary history of a tumor with one subclone. After accumulation of *n* drivers, the first cancer cell gives rise to the tumor (founder clone in blue), with the population evolving neutrally and accumulating passenger mutations (orange), until eventually a subclonal driver event occurs triggering a subclonal expansion (green). Both the background clone and the new subclone contain their own neutral tails. The subclonal driver event, together with its passenger hitchhiker mutations (orange) will rise in frequency with the subclonal expansion and be part of the subclone cluster in the VAF distribution. However, some early hitchhikers are also present elsewhere in the tumor as part of the tail of the founder clone. In the simple example of perfect cell doubling, we expect mutations in the first doubling to be in 50% of the cells of the tumor, mutations in the second doubling to be in 25% of the cells in the tumor etc. (**B**) If we take biopsies S1 and S2, we find the founder clone (S1) and a subclonal sweep (S2). The hitchhiker mirage is a confounder determined by passengers that hitchhike to the subclonal driver in S2, but diffuse neutrally in S1 (orange). This can be seen in the VAF distributions of the two samples where the orange mutations do not travel together in the two samples. (**C**) The VAF of S1 versus S2 shows that orange hitchhikers can generate an extra cluster with Binomial parameters (S1=0.5, S2=0.2), on top of the true green clone with different Binomial parameters (S1=0.5, S2 = 0). On top of this, extra clusters can be generated by fitting tail mutations with a Binomial mixture, further inflating the true number of clones (*k* = 2) and suggesting false clonal sweeps. (**D**) If we remove mutations assigned to a tail (projected VAF) by `MOBSTER`, we clean up the signal and we can retrieve the true clonal architecture. (**E**) The ancestor effect is due to the fact that, due to spatial structure in the tumor, cells from a spatially localized biopsy will always have a more recent common ancestor, compared to other biopsies. Hence, mutations that are observed at high frequency in one biopsy are not necessarily due to selection. (**F**) We simulate the growth of a neutral tumor in 2D, and sample two bulk biopsies (100% purity). Both samples will contain truncal mutations. Each biopsy also contains private mutations (green and orange) that are clonal within the sample but are not due to selection. When we generate a virtual staining of all cells that harbor the mutations in a cluster, we can see the separation between cells in S1 and S2, and the branched evolutionary structure in the clone tree that is not due to selection, but to spatial sampling. (**G**) The admixing deception happens when there is some level of spatial intermixing in the tumor and spatially close cells within a biopsy are genetically distant in the tumor phylogenetic tree. In this example, whereas S1 is a bulk of closely related cells, suffering only from the ancestor fallacy, S2 contains a mixture of cell lineages from distinct parts of the phylogenetic tree (here split between right and left side of the phylogeny). Even with low levels of intermixing, this is bound to happen as somewhere in the tumor two distant parts of a phylogeny must intermix. Again, in this example, no selection is at play. (**H**) If we sequence these two biopsies, we will find truncal mutations in black and mutations private to S1 in green (ancestor fallacy). In S2 we will find a mixture of distant lineages (orange and blue) that look like subclonal clusters (here we neglected neutral tails for simplicity). The orange and blue clusters deviate by an offset from the clonal peak that is determined by amount of admixing, which is unknown a priori (see the marginal VAF distribution of S2 on the left). (**I**) All these confounders lead to additional nodes in the estimated clone tree. (**J**) Translating these nodes into phenotypically distinct subpopulations is misleading, as the reasonable representation (**K**) would be instead much simpler. (**L**) Phylogenetic tree at the single cell level of the tumor in (**I**) showing that clusters B, C, E and F are arbitrary ancestors identified by the specific bias of the measurement.

### The *hitchhiker mirage*

Here we comment on an artifactual effect inherent to multi-region data that we term the hitchhiker mirage. In Figure 4A we show the schematic of a tumor with a founder clone (blue) that gives raise to a new positively selected subclone (green). Two spatially distinct samples were collected from the tumor, respectively containing only the original clone (S1) and the new subclone with selective advantage (S2); see also Supplementary Figure S9. Sequencing data from sample S2 would contain the subclonal driver event that gave rise to the new subclone (clonal in S2), and both samples contain neutral passengers. The marginal VAF distribution of each sample shows the expected neutral tail of passengers (Figure 4B). However, the joint tumor distribution (here 2-sample VAF distribution) shows multiple additional clusters of mutations (Figure 4C). The cluster(s) at low frequency (Figure 4B-C; red) are due to neutral tails, analogous to the single sample case. The source of the cluster at higher frequency (Figure 4C; yellow) is much subtler and relates to passenger mutations that accrued in the lineage that went onto form the selected subclone (orange mutations). When the subclone expanded, these passenger mutations that had accrued prior to clone growth were present in every cell in the clone, and therefore appeared clonal in sample S2 (hitchhiker mutations). However, some of these hitchhiker mutations (orange) were also contributed to other lineages that formed other parts of the tumor, at varying frequencies. For example, for a series of cell doublings, mutations at cell division 1, 2, 3 … will have frequency 50%, 25%, 12.5%, 6.25% etc. in the rest of the tumor. When these passengers were also then detected in sample S1, a clustering algorithm could “break” the group of subclonal mutations (orange and green) in two clusters with different Binomial parameters (Figure 4C). Statistically, this split of orange and green mutations is consistent with the fact that these alleles really move differently across biopsies, but the produced groups have little to do with the actual evolution of this tumor, and are rather determined by spatial sampling. In particular, the orange cluster does not correspond to a real subclone under positive selection; rather it is composed by neutrally evolving ancestors that they just happen to end up in S1, and a slightly different sampling strategy would have identified a different ancestor.

Prior removal of neutral tails solves this problem of ‘splitting’ hitchhiking mutations. In the cartoon example (Figure 4A), running `MOBSTER` separately on each sample can identify and remove neutral tail mutations from the orange group, so that clustering of the remaining read counts (projected VAF after `MOBSTER`, Figure 4D) correctly identifies only one single cluster/ subclone in the data.

### The *ancestor effect*

Mutations found at high frequency in a small sample of a larger population can be erroneously associated with selection. Just due to spatial structure, cells that are close in space are also likely to be close genetically and tend to have a recent common ancestor. We illustrate this in Figure 4E where we assumed a completely neutral tumor driven only by clonal drivers; see also Supplementary Figure S9. When we sampled two biopsies S1 and S2, in each sample we detected the clonal mutations corresponding to each sample’s most recent common ancestor (MRCA). In this case, the simulated tumor was evolving entirely neutrally, and no differentially selected clones existed.

Multi-region sampling inevitably resolves the clonal ancestry of the samples (e.g. finds subclones) but these again were just arbitrary, neutrally evolving ancestors picked up by the spatial sampling. We illustrate this in Figure 4F where the two “clones” private to each biopsy (S1: green; S2: orange) were used to reconstruct the clone tree and the cells in the simulation have been virtually stained for the mutations in each identified cluster (including the clonal cluster). Hence, in multi-region sequencing data, the identification of clusters (subclones) is an inevitable consequence of the repeated sampling process and, most importantly, does not necessarily imply underlying subclonal selective forces within the tumor.

### The *admixing deception*

Sampling bias is particularly problematic when there is some level of admixing of distant lineages within a sample. By definition, we can divide the tumor in two ancestral lineages (notionally “left” and “right”) that are the decedents of the first branch point of the phylogenetic tree (i.e. the first cancer cell division to produce two surviving lineages). This is illustrated for the simplest possible neutral case in Figure 4G, and we note the scenario is further complicated when positively selected subclones are present (Supplementary Figure S9). We analyze two spatially distinct tumor samples in this simple neutral case. Sample S1 contains cells entirely from the left lineage, whereas sample S2 contains, by chance, cells from both left and right lineages. This is bound to happen as somewhere in the tumor two distant parts of a phylogeny must meet. Sample S2 contains a large proportion of the lineages coming from the right tree (80%, blue), and a minor proportion of the lineages coming from the left part of the tree (20%, orange). These two groups of lineages have very distant MRCAs despite being spatially close, which causes the detection of subclones that are however entirely due to neutral evolution (Figure 4H). Both samples contain the clonal mutations (black), but only sample S1 contains the green mutations (ancestor effect). In S2, however, the observed adjusted VAF can split into two subclonal clusters at CCF ≈ 80% (adjusted VAF 0.4, blue) and ≈ 20% (adjusted VAF 0.1, orange). There are strong implications for our ability to identify the true generative model: the same subclone distribution can also be obtained from a tumor with subclones under selection, and distinguishing between admixing and positive selection is therefore extremely challenging.

This confounder needs to be considered because we expect clonal admixture within samples. The admixing skews the frequency of the dominant lineage in a sample by a small gap, enough to separate it from the clonal mutations (blue cluster is skewed to the left of the black cluster in Figure 4H). We note that a large number of state-of-the-art analyses report subclones at high CCF value in a majority of samples^19,25–27^, and this phenomenon can happen also when a single sample is taken. We argue that this “photo finish” effect of apparently multiple subclones about to sweep through the population could be also due to the *admixing deception*. Such clonal structures are unlikely to be real also because the dynamics of selection in expanding populations bias the CCF of subclones towards 1 or 0^28^, thus indicating that large subclones about to sweep should be rarely detected.

### From the tumor clone tree to its evolutionary history

`MOBSTER` pre-filtering can address the hitchhiker mirage, but there is no systematic way to correct the other two types of error. Instead, careful, skeptical, interpretation of the results is needed. In Figures 4I we show a hypothetical tree of putative clones recovered after `MOBSTER` filtering of tail mutations. The tree contains only two positively selected clones (A and D). The other clones (B, C and E) are just due to the confounding effects discussed above and represent arbitrary ancestors that evolved neutrally. There is nothing “special” about those ancestors: they are not phenotypically distinct subpopulations, and they have not experienced subclonal selection. Drawing a Muller plot of the evolutionary history of the tumor from this clone tree (Figure 4J) would give a misleading picture of the history of subclonal selection in this tumor (contrast with Figure 4K, where the true subclonal expansion is represented). This can be more clearly appreciated from the phylogenetic tree of the individual cells shown in Figure 4L, where the true subclone is annotated in orange. When we map the “clones” from the clone tree into the true cell phylogeny, we can see that clones C, B and E are just random ancestors in the phylogeny. If we resampled the tumor again, we would have picked different ancestors every time, obtaining a different clone tree. This is also true for the branching structure of the tree, which depends on cluster C. For this reason, drawing conclusions from the structure (e.g., linear, branching) or the size (e.g., number of clones) of the clone tree without accounting for these confounding factors, can likely lead to false conclusions.

### Guidance for subclonal reconstruction with MOBSTER using multiple biopsies

We used simulated data from a stochastic spatial branching process model of tumor growth^15^ to assess the confounding factors of spatial sampling discussed above, and to provide a rationale to handle them and interpret the data correctly.

In Figure 5A we show a simulated example of a 2D neutral tumor for which we perform 100x WGS of two bulk samples. `MOBSTER` can filter out the tails from both samples (Figure 5B). Analysis of the joint adjusted VAF distribution shows the relationship between the mutations in the samples (Figure 5C), which cluster into *k* = 3 groups: one clonal cluster (green), and two subclonal clusters due to the ancestor fallacy (one private to S1, and one private to S2). Due to the admixing deception, the mean VAFs of these two clusters are slightly shifted below the expected clonal peak (≈ 0.5 adjusted VAF). The contamination of mixing lineages is minimal, and the second lower-frequency VAF cluster cannot be detected at the resolution of 100x WGS. The effects of sampling a MRCA can be clearly appreciated in the virtual staining of the spatial distribution of the mutations (we color the cells based on mutational cluster assignments), where we see that a tumor split by sampling into two subpopulations misleads subclonal estimation. Clustering without `MOBSTER` suffers from a much worse overfit effect, and detects twice as many clones (*k* = 6) with two false positive clusters of private tail mutations (clusters C1 and C2 in Figure 5D).

**Figure 5.**
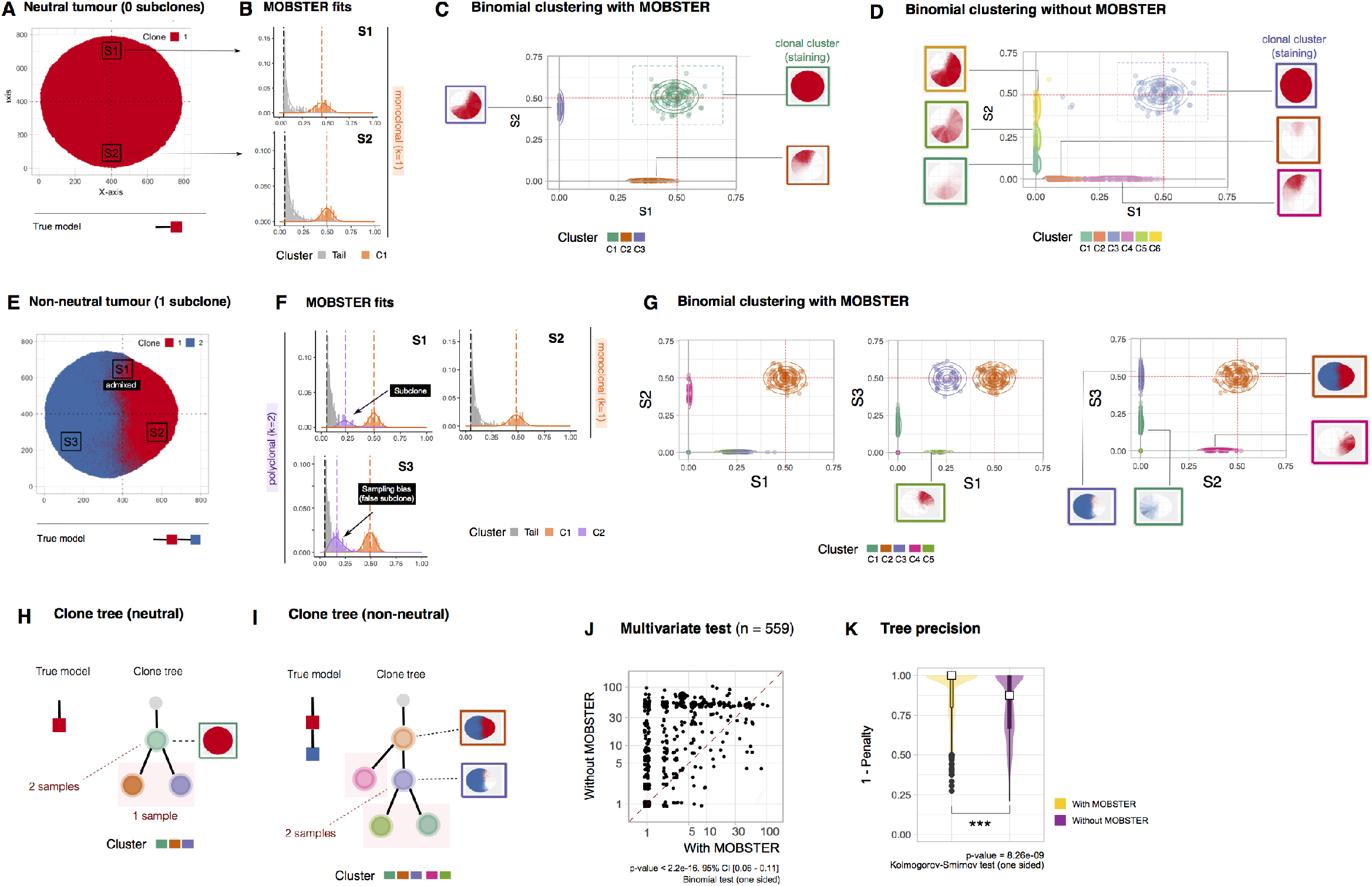
MOBSTER performance on multi-region sequencing. (**A**) Spatial simulation of a tumor without subclones (i.e., neutral) where two samples were collected and sequenced at 100x WGS. (**B**) `MOBSTER` fit of two monoclonal biopsies shows that tails can be easily removed. (**C**) A multivariate analysis of non-tail mutations with a variational Binomial mixture (Supplementary Note 2) after `MOBSTER` identifies *k* = 3 clusters (one truncal and two private to each biopsy), showing the admixing deception (shift of blue and yellow clusters from the 0.5 clonal expectation). The staining of these clusters shows the ancestor effect as well. (**D**) When we do not control for tails we detect *k* = 6 clusters and a much more complex evolutionary history; staining of the mutations identified with this clustering shows the error. (**E**) Spatial simulation of a tumor with one subclone under positive selection (blue), and collection of 3 biopsies (one boundary, and two monoclonal). (**F**) The fit of `MOBSTER` to S1 and S2 is perfect, but in S3 `MOBSTER` calls an extra subclone due to genetic drift (the true tail deviate from a power law, resulting in a subclonal cluster). (**G**) When we analyze read counts after `MOBSTER` we detect a clear subclone in S1 against S3. This is correct because the subclone has swept through only S3, while it is admixed to its ancestor in S1. In all samples the ancestor fallacy and admixing deception are clearly observable. (**H**) When we try to detect the actual clones under positive selection and their clonal architecture, a sensible and conservative heuristic is to consider only clusters detected in a minimum number of biopsies. We note that removing private subclones can underfit, as in the case of a genuine subclone detected in a single biopsy. In these tumors, removing leaf subclones for both the neutral case and the subclonal selection case leads to the determination of the true model. (**J**) Performance of `MOBSTER` versus standard methods for multi-region sequencing analysis, with *n* = 559 simulated 2D tumors with a number of biopsies that ranges from 2 to 9. Each tumor contains either 0, 1 or 2 subclones, and we measure statistics from the reconstructed clone trees after variational Binomial clustering of read counts after `MOBSTER` (see also Supplementary Figure S10). The trees are constructed via the pigeonhole principle, whose violations are measured with a score. Here we show the number of possible trees with and without `MOBSTER`. The points in the upper/lower diagonal of the panel show which approach (with/without `MOBSTER`) give less clone tree fits to the simulated tumor. From then number of points in each part of the panel we derive a p-values via standard one-sided Binomial test, which shows that the reduction in the uncertainty of the tree estimate with `MOBSTER` is a statistically significant. (**K**) The distribution of the pigeonhole violations as one minus the penalty score assigned to each tree; values closer to 1 reflect fewer violations (0 penalty). We test for the difference with a one-sided Kolmogorov-Smirnov test. These plots show that with `MOBSTER` we systematically retrieve fewer clonal trees with consistent higher quality, at a rate that is significantly higher (*p* < 0.001 in both tests). In Supplementary Figure S10 we plot results by number of biopsies, showing that the error rate without `MOBSTER` is dramatic, and increases with the number of samples taken.

In Figure 5E we use the same principles to show that similar errors are present when subclonal selection is simulated, and a new subclone emerges within the background (blue staining). Three biopsies of this tumor are sampled, and `MOBSTER` is used to identify and filter out tail mutations (Figure 5F). The VAF distribution for the boundary sample S1, which has a clear bump as predicted by theory, is a clear mixture of background and selected subclone. Sample S2 is entirely composed of the background clone, and shows intra-sample neutral dynamics. Sample S3, instead, shows a small cluster due to sampling bias and genetic drift, which is misleading because the biopsy is monoclonal and contains only the selected subclone. Multivariate subclonal reconstruction run after `MOBSTER` detects *k* = 5 clusters (Figure 5G). Virtual staining of each cluster clearly shows that multiple clusters are the result of neutral drift (C1), ancestor fallacy (C4, C5) and admixing deception (C1, C4 and C5, which are shifted). This is where informed interpretation of the data needs to take place after subclonal reconstruction.

In Figure 5H we show a conservative approach to reason on the output of subclonal reconstruction. In the neutral tumor from Figure 5A the true model is a single monoclonal population. The subclonal reconstruction identifies *k* = 3 clusters, but two of them are just due to sampling two bulks. Being private to each sample they cannot be discriminated from the ancestor fallacy. A conservative analysis could then eliminate them, recovering the true model. Importantly, the exact same phylogeny could be obtained with two monoclonal samples from two distinct clones (e.g. equivalent to S2 and S3 in Figure 5E; see also Supplementary Figure S9). In that case, however, it is reasonable to expect the length of the two branches outgoing C1 to be significantly different. A longer branch can indicate possible selection, under the assumption that the size of the biopsy samples is similar; if this were not the case, biopsy size would act as an extra confounder for branches length (larger biopsies have more cells, and therefore less private mutations as their MRCA is further back in time). In the non-neutral tumor, a more complex clone tree is recovered because three samples were taken. The true model is a founder clone which gives rise to one positively selected subclone. Cluster C2 corresponds to the founder clone and contains only truncal mutations. Cluster C3 contains mutations that are in the common ancestors of both C5 and C1, but not C4. Importantly, all the leaves are clusters that are private to each sample. The most conservative clonal tree is then “linear evolution” of C2 (clonal) to subclone C3, the true tumor evolution. We note that the additional power in calling C3 a true subclone comes from the fact that the subclone has been observed in multiple biopsies.

We tested the performance of `MOBSTER` against standard subclonal reconstruction with *n* = 559 simulated tumors, with different subclonal architectures and multi-region sampling. Every simulated tumor contains either 0,1 or 2 subclones which grow in 2D from a single initial cell; we simulate collection and sequencing of 2 to 9 equal-size biopsies at 100x whole genome, assuming ideal purity (100%) and diploid genomes. In this setting we could isolate and measure the effect of the spatial confounders introduced in this section. In each run we measured statistics from the clone trees that we can reconstruct processing the output clusters; for read counts clustering we used a variational Binomial mixture model akin to sciClone^7^ (Supplementary Note 2), combined with and without `MOBSTER`. We are interested in solving confounders that augment the number of trees, and measure their effect on the goodness of fit. We find many less possible alternative trees (*p* < 0.001, one-sided Binomial test) constructed via the standard pigeonhole principle^24^ when we run `MOBSTER`, compared to a standard method (Figure 5J). This trend is observed in more than 400 cases, in a large number of cases we find one tree when we run first `MOBSTER` on the data, compared to a number of trees that spans from 5 to about 100 when we use a standard method. Precisely, in 38% (*n* = 212) of simulated cases we identify only one tree with `MOBSTER` while several trees without, while in 16% (*n* = 92) of cases both analyses find only one tree. Moreover, when we measure the violations of the pigeonhole principle as a proxy for the goodness of fit, we find that trees derived after `MOBSTER` have median 0 violations (perfect fit), while trees without `MOBSTER` have systematic violations of the principle (Figure 5K). The number of violations with `MOBSTER` is significantly lower (*p* < 0.001, one-sided Kolmogorov-Smirnov test). Results expanded by number of biopsies and subclones in the simulated tumor confirm this trend, and show that the error rate without `MOBSTER` is large, and increases with the number of samples taken (Supplementary Figure S10).

This shows that the spatial confounders that we have identified are driving the output of subclonal deconvolution, especially when we collect multiple tumor biopsies. By leveraging on the higher precision of `MOBSTER`, combined with *a posteriori* inspection and curation, it is possible to minimize the effect of these confounders, and to focus on clusters likely representing truly selected subclones.

### Analysis of genomic data from human cancers

We applied `MOBSTER` to the highest resolution (>100x) whole-genome sequencing data that is available in the public domain. As we have shown, purity at this resolution can become a confounder for tail detection, and it is crucial to achieve suitable coverage for this type of analysis. We first re-analyzed the breast cancer sample PD1420a sequenced at 200x depth by Nik-Zainal *et al*.^3^, which has ≈70% tumor purity and *n* = 4,643 SNVs in a highly confident diploid chromosome 3. Compared to the original analysis which found 3 subclones and in agreement with our previous work^8^, the analysis with `MOBSTER` unravels that the lowest frequency cluster is actually a neutral tail (Figure 6A). This tail is as large as the largest true subclone – i.e., ≈ 1,000 SNVs which account for ≈ 20% of the overall mutational burden, compared to the larger subclone that has ≈ 1,100 SNVs. sciClone’s analysis of read counts for non-tail mutations confirms *k* = 3 Binomial clusters (2 subclones), and not 3 subclones as originally proposed^3^.

**Figure 6.**
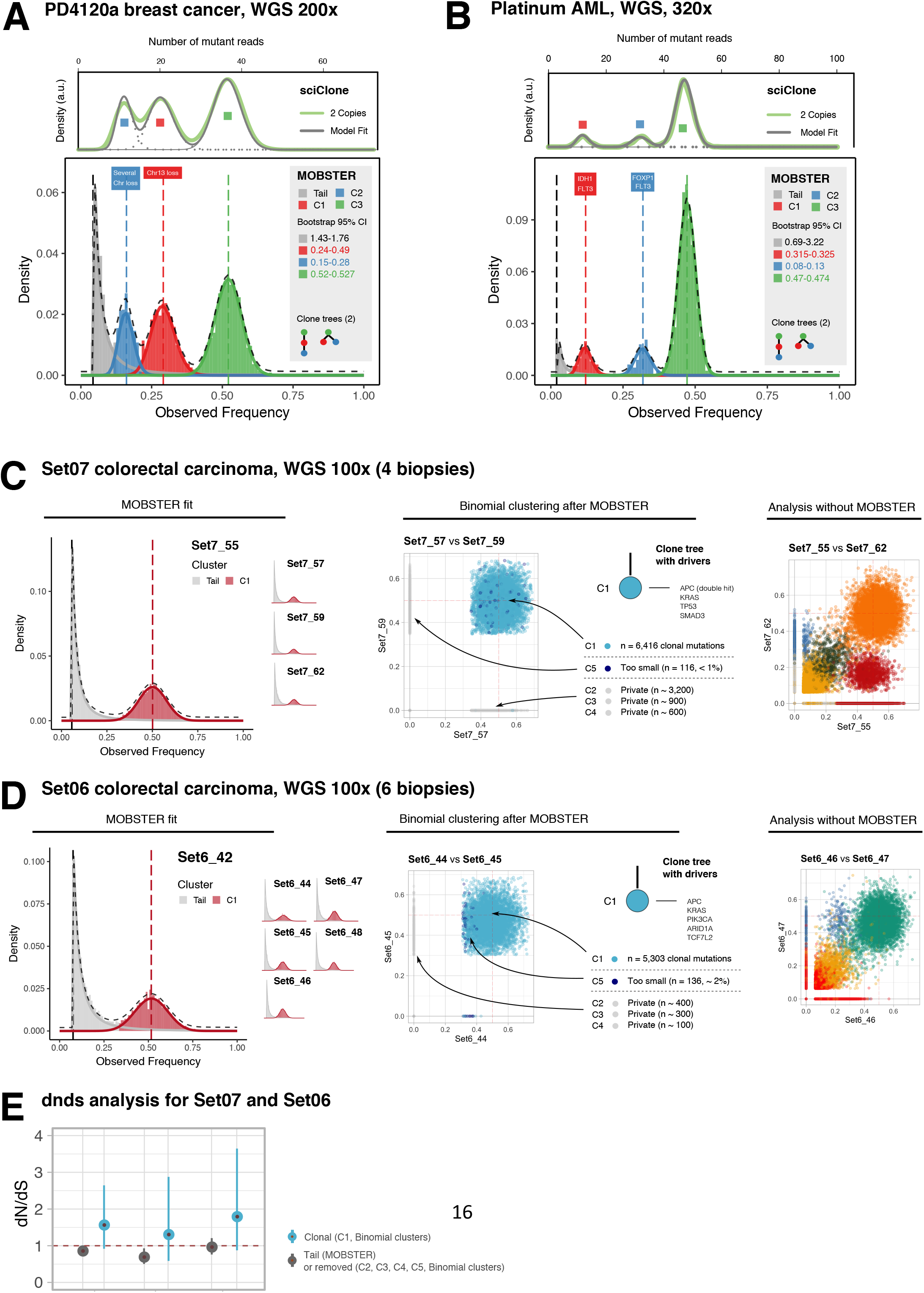
Analysis of real genomic data. (**A**) The PD4120a is a breast carcinoma sample analyzed ~200x WGS. Here we use >4000 SNVs in diploid regions from chromosome 3 (which is all diploid) to run `MOBSTER` and identify a clear tail plus *k* = 3 Beta clusters (2 subclones) that we use to select read counts for sciClone thus confirming 3 clones in the data (2 subclones). Notably, sciClone without `MOBSTER` calls an extra clone instead of the tail. A non-parametric bootstrap procedure is used to estimate the 95% confidence intervals for the parameters (tail’s shape and Beta means). Two possible clone trees can be fit to these clusters via the pigeonhole principle (linear and branched tree). Compared to the original analysis^3^, we are able to retrieve a more precise evolutionary history of this tumor. (**B**) The AML Platinum is a leukemia sample sequenced at 320x WGS. When we analyze >1000 diploid SNVs from this patient, we find a large clonal cluster and two low-frequency subclones (*k* = 3). The tail of this tumor is quite small compared to the one in the breast sample, indicating lower mutation rates. Subsequent analysis with sciClone confirms two subclones with the same clustering assignments, and 2 possible clone trees that fit the clusters. Even in this case sciClone without `MOBSTER` calls an extra clone instead of the tail (not shown). (**C**) We generated new WGS sequencing data at 100x coverage for 4 biopsies of colorectal cancer patient Set07. We identified reliable diploid regions of this tumour, from which we obtained >50,000 SNVs. Median sample purity was >80% and overall ploidy was two (Online Methods). We then analyzed each individual biopsy with `MOBSTER`, identified neutral tails and found a simple monoclonal architecture in each of the four samples. After removal of tail mutations we run the variational multivariate Binomial method that is available within `MOBSTER` on the remaining SNVs (Binomial fit with stringent concentration value *α* = 10^−9^), which finds *k* = 5 clusters. Cluster C1 is the clonal cluster. Clusters C2, C3 and C4 are private to a biopsy, suggesting ancestor effect. Cluster C5 is then removed as it accounts only less than 1% of non-tail mutations. The clone tree for this patient depicts thus a perfect neutrally evolving tumor; a comparative analysis without `MOBSTER` would have dramatically inflated the estimate of subclonal selection, depicting a wrong clonal history for this patient (Supplementary Figures S11-S14). In the clonal tree we annotate driver SNVs validated by Cross *et al*. from well-known colorectal driver genes. (**D**) We generated new WGS sequencing data at 100x coverage for 6 biopsies of another colorectal cancer patient Set06. Also in this case the analysis shows a neutral tumor with no evidence of positive subclonal selection in any of the sequenced regions. With a similar analysis we conclude that also this tumor is monoclonal, a finding that the analysis without `MOBSTER` would have largely missed due to the strong effect of spatial and evolutionary confounders in these data. (**E**) dN/dS analysis for Set06 and Set07 comparing mutations detected as clonal (C1), versus those removed because found in a tail by `MOBSTER`, or because we filter output clusters as explained in panels C and D. For each estimate we report its 95% confidence interval. These values confirm lack of evidence for positive selection at the subclonal level, corroborating our analysis.

Two possible clone trees can be fit to the output clusters (linear and branched evolution), and the branching tree matches the tree published in the original analysis, which used phasing to resolve this ambiguity and select the branching model^3^. We note that the extra cluster detected in the original paper could not be attached to the tumor clone tree in a unique position, and it rather spreads over multiple leaves of the clone tree (cluster A in Figure 3D of Nik-Zainal et al.^3^). Our analysis with `MOBSTER` shows that the low frequency “cluster” is not a clone, but just the result of neutral within-clone dynamics possibly generating from both of the tumor subclones. We measured the evolutionary parameters of this tumor from `MOBSTER` fits, and found concordant estimates with our previous work^8^. This tumor has a mutation rate *μ* = 3.5 * 10^−7^ mutations per base pairs per tumor doubling. The subclones emerged at times *t* = 5.5 (smaller subclone) and *t* = 10.4 (larger subclone) when estimated in tumor doubling times, and have different selective advantage coefficients *s* = 0.3 and *s* = 0.66 respectively – these values quantify how fast the subclone grows compared to the ancestor.

We obtain a similar result when we re-analyze *n* = 1,332 SNVs in diploid regions of the primary AML Platinum sample, for which Griffith *et al*. had identified 3 subclones^29^. This sample is sequenced at 320x whole-genome with high tumor purity (>90%). Running `MOBSTER` identifies again *k* = 3 clusters, but only 2 subclones with 103 and 116 SNVs each, and a neutral tail (Figure 6B). The same two subclones were detected also by Griffith *et al*.^29^, and were confirmed running sciClone, the same tool used in the original paper, after `MOBSTER` (Figure 6D). In this tumor the tail that we fit is much smaller (66 SNVs) compared to the breast sample, suggesting possibly lower mutation rate for this tumor. As the breast sample, output clusters can be fit to both a linear and a branching clone tree without violating the pigeonhole principle^24^; to further resolve the clonal evolution of this patient other samples have to be used, as in the original analysis which used a whole-exome relapse sample^29^. With `MOBSTER` we simplify the clonal structure of this primary tumour removing one spurious low frequency “subclone”, compared to what has been estimated in the original analysis. This improves the interpretation of these data: in the original analysis, in fact, it was observed that the only cluster without a subclonal driver mutation was the one that we actually identify as tail. As for the breast sample, measurement of the evolutionary parameters that we can obtain from the fits, are again concordant with our previous work^8^. This tumor has a much lower mutation rate *μ* = 9.9 * 10^−10^, as suggested by the lower mutational load in the tumor tail; the subclones emerged at similar times *t* = 22 and *t* = 27, and have quite different selective advantage coefficients *s* = 1.3 and *s* = 3 respectively, which is why one of the two reaches much higher proportions compared to the other.

We then generated new multi-region whole-genome sequencing data at median coverage 100x from multiple regions of two colorectal cancers, Set06 (6 regions) and Set07 (4 regions), previously analyzed at lower sequencing depth by Cross *et al*.^30^ with standard phylogenetic methods. We analyzed, for each patient, high-quality SNVs in diploid regions identified across the full genomes of all the patient’s samples, and run the analysis with and without `MOBSTER` (Online Methods). For Set07 we analyzed ≈ 50,000 SNVs (Supplementary Figures S11 and S12) in high-purity samples with ≥80% tumor cellularity. The analysis with `MOBSTER` shows no evidence of positive subclonal selection (Figure 6C), corroborated by the lack of any subclonal driver alteration. This was consistent with the original findings by Cross et al., showing balanced phylogenetic trees and lack of subclonal drivers, supporting neutral subclonal dynamics. The downstream analysis with tail mutations filtered out confirmed a simple monoclonal structure for this tumor, which harbors driver clonal mutations in *APC* (double hit), *KRAS, SMAD3* and *TP53*. Not surprisingly, the analysis without `MOBSTER` depicts a totally different clonal architecture with a much more complex clone tree (5 clusters, which accounts for 4 subclones; Supplementary Figures S13 and S14).

The analysis for Set06 gives very similar results. Using the same strategy adopted for Set07, we could analyze ≈30,000 SNVs (Supplementary Figures S15 and S16) in high-purity samples with ≈ 80% tumor cellularity. Consistent with Cross et al., the clone tree harbored only clonal driver mutations in *APC, KRAS, PIK3CA, ARID1A* and *TCF7L2*. Also in this case the standard analysis would have identified an inflated clonal architecture with 4 subclones, and a much more complex clone tree (Supplementary Figures S17 and S18). By sequencing at higher resolution two of the cases previously analyzed by Cross *et al*.^30^ and by employing a `MOBSTER`-based analysis, we reach concordant conclusions with the original study even if the adopted methodologies are different. The striking difference between the results obtained with and without `MOBSTER` confirms the importance of controlling for tails in a multivariate analysis. For both patients, a standard method would have portrayed totally different evolutionary histories, giving the illusion of positive subclonal selection in these patients. Concerning the evolutionary history of Set06 and Set07, we could measure mutation rates per sample very precisely from every tail fit; the median estimates per patient was found to be *μ* = 5.6 * 10^−7^ (mutations per base pairs per tumor doubling) for Set07, and *μ* = 4.3 * 10^−7^ for Set06. These concordant estimates are consistent with the fact that these tumors have comparable mutational load, similar type of clonal architecture and related evolutionary trajectories. Notably, orthogonal dN/dS analysis that uses the ratio of non-synonymous to synonymous mutations to detect selection^31,32^ confirms the lack of evidence of positive selection at the subclonal level in data from both tumours (Figure 6E, Online Methods).

## Discussion

Subclonal reconstruction from cancer bulk sequencing data has paved the way for the study of intra-tumor heterogeneity (ITH) and evolutionary cancer dynamics^3,33^. Measurement of subclonal architectures have also clinical relevance: subclone multiplicity and other measures of ITH are prognostic biomarkers^34–36^. Naturally therefore, there is a need to ensure that subclonal reconstructions are accurate.

Here we have presented a model-based subclonal reconstruction method that is rooted in theoretical population genetics. Subclonal reconstruction is performed exploiting a generative model of the data. This is in contrast to purely data-driven approaches that lack an underlying evolutionary model, and here we have showed that this evolutionary model-based method outperforms standard data-driven approaches. Moreover, we have identified fundamental confounding factors inherent to the subclonal analysis of bulk sequencing data, and we have demonstrated that correcting for neutral tumor evolution is necessary for reliable evolutionary analyses.

In general, our analysis shows that extreme care must be taken to infer subclonal architectures from the available “bulk” sequencing data, and that in some cases it may not be possible to infer an accurate subclonal architecture at all. Sequencing depth is a key determinant of data quality, and we suggest that only high depth sequencing data (>100x) is appropriate to infer subclonal architectures. Subclonal reconstruction from low-depth data risks a systematic over-calling of spurious subclones, jeopardizing our attempts to retrieve the correct life history of tumors. Various issues arise in multi-region sequencing data that ultimately result from biases that are intrinsic to spatial sampling. These issues lead to inflated estimates of positive subclonal selection from VAF distributions. Single-cell sequencing is becoming more common, and we note that while it removes the problem of admixing of populations^37^, issues of spatial sampling bias remains, especially if individuals cells are isolated from spatially localised bulks^38–41^.

In general, we have presented the case for the rigorous examination of cancer sequencing data in light of quantitative generative models of tumor evolution that describe 1) how cancers grow, 2) how subclones form and 3) how spatial sampling strategies affect our estimates. Critically, we should also recognize the intrinsic limitations of current data, foremost the systematic confounding factors caused by sampling of complex threedimensional tumors. Our analysis represents a step towards a more refined approach to subclonal reconstruction in bulk cancer data.

## Acknowledgements

A.S. is supported by the Wellcome Trust (202778/B/16/Z) and Cancer Research UK (A22909). T.G. is supported by the Wellcome Trust (202778/Z/16/Z) and Cancer Research UK (A19771). We acknowledge funding from the National Institute of Health (NCI U54 CA217376) to A.S and T.A.G. This work was also supported a Wellcome Trust award to the Centre for Evolution and Cancer (105104/Z/14/Z). C.P.B. acknowledges funding from the Wellcome Trust (209409/Z/17/Z).

## Authors contribution

GC conceived, designed and implemented the method. TH and KC developed the spatial tumor growth simulations; TH and MW generated the data for synthetic tests, which were carried out and analyzed by GC, TH, MW and DN. GC, MW and LZ analyzed the data, with input and support from WC, GDC and AA. GS, CB, TAG and AS supervised method design. AS and TAG conceived and supervised the study. All authors contributed to and approved the manuscript.

## Online Methods

**Availability.** `MOBSTER` will be hosted at https://github.com/caravagn/MOBSTER

The repository will be made accessible over the next months as soon as the preliminary implementation is turned into a stable software package, accompanied by a manual. The tool will include documentation in the form of RMarkdown vignettes, as well as data from all case studies (simulated and patient derived).

Please write to Giulio Caravagna to receive further information on the tool and its availability (preliminary versions etc.).

### Model-Based Clustering of Subclonal Populations in Cancer (MOBSTER)

The subclonal deconvolution problem is popular in the cancer literature^42^. Given read counts for a list of Single Nucleotide Variants (SNVs) detected from bulk sequencing of multiple tumor samples, we want to detect clusters of SNVs that represent cancer subpopulations admixed in our samples. We shall call them clones under positive selective pressures, and detect them from the frequency of their mutant alleles.

`MOBSTER` is a mixed method that combines two types of random variables to approach this problem.

#### The frequency spectrum and the observational process

Kessler and Levin^11^ have shown that, in the large population solution of the stochastic Luria-Delbrück model, the probability of having *m* mutants follows a fat-tail Landau distribution

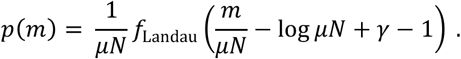

Here *N* is population size, *μ* the average fraction of birth events and *γ* a constant. The asymptotic behaviour of *f*_Landau_ can be approximated as

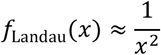

which leads to the power-law approximation that has also been derived by others^13,43,44^

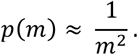

A generative model for this power laws can be constructed with a standard Markovian stochastic birth-death process of cell division – sometimes called *branching process*^8^. The existence of patterns of neutral evolution is thus a consolidated result from Population Genetics arguments that describe the spread of alleles in growing populations without recombination, such as cancer^16^. Very simply, the progeny of each clone accumulates neutral passenger mutations until any of their daughter cells acquires a new mutation (SNV just for simplicity) that undergoes selection: the power-law spectrum emerges by the frequencies of passengers. When a daughter cell undergoes selection, however, the frequency of its variant alleles will grow, eventually becoming detectable if selection forces are strong compared to background. In turn, the progeny of this new subclone will start dividing, giving rise to another power-law distributed tail.

Importantly, the power-law part of the spectrum – i.e., the *tail* – results from the accumulation of passenger mutations in the progeny of any clone. We note that this result – in particular the exponent 2 – refers to the overall population structure of the tumor, and that any specific finite sample that we collect and sequence, which might also be contaminated by normal cells, might exhibit deviations from this theoretical distribution^8^. Deviations from strict exponential growth due to for example constrained spatial can also cause deviations from exponent two^43,45^. However, we can use this result to create a model-based approach to analyze cancer data.

We assume we work with copy number and purity adjusted frequency values for the variant alleles of *n* SNVs. With *k* ≥ 1 detectable clones, a reasonable model for the *frequency spectrum ρ* of the observed SNVs is a random variable that follows

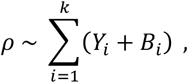

where

- *Y_i_* ∝ *x*^−*α*^ is a power-law random variable for frequencies of neutral SNVs in the progeny of clone *i*. The generic exponent *α* > 0 gives flexibility to accommodate all the confounders described above;
- *B_i_* ∈ [0,1] is a Beta random variable modelling the signal of clone *i*. In layman terms, *B_i_* models the “bump” in the frequency distribution due to the clones. These distributions range in [0,1], rendering them suitable to describe allelic frequencies.

This model looks simple, but further observations are required to turn it into a standard mixture-model. First of all, the random variables for the tail and the bump of every clone are coupled to capture a joint signal. While the overall mixing proportions can be assumed to be independent, this compound random variable requires an extra level of mixing within each clone – i.e., to properly capture the proportions of the clone tail, and bump. This would require extra parameters to fit the model, but it does not seem particularly necessary because in the end we are interested in the information that we derive from the cluster of each clone, which we use to identify subpopulations in the frequency spectrum. Precisely, we use the clone’s peak, obtained from the cluster’s mean, to assess the phylogenetic history of the tumor.

We can simplify this model and retrieve a simpler formulation by noting that all tails have the same exponent *α*, as they are described by the same theoretical distribution across clones. For this reason, they are multiple copies of the same random variable. Thus, we can group them together in a single power-law tail

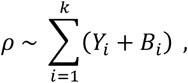

Here the random variables have the same meaning as above, but the clone is no longer indexed by *k*. This model has a key advantage over the one where each clone “emits” his own tail. Here the random variables are decoupled and allow a simple mixture-model formulation which we will present below.

Before concluding, we observe that given *ρ*, the *observational model* for read counts collected from NGS sequencing, is a standard binomial process

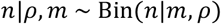

where *m* is the coverage (total number of reads), and *w* the number of reads harbouring the variant allele; *ρ* is then the success probability for *m* iid Bernoulli trials. It is important to observe that the frequency spectrum and the observational process look at the data from different perspectives: the former is a distribution on allelic frequencies, while the latter on read counts. In this observational model we can in principle use Beta-Binomial distributions to account for coverage overdispersion.

#### Relation to other models in the literature

The literature is rich with models that describe the above observational process and variation thereof, either with Binomial or Beta-Binomial distributions. We briefly discuss those that are more related to our framework.

Bayesian methods that employ Dirichlet Processes for infinite mixture models are a popular generalization of the observational process. These semi-parametric/ non-parametric methods can fit an unspecified number of clusters *k* to the data, simplifying model selection procedures. `pyClone`^23^, `DPclust`^3^ and `PhyloWGS`^6^ are three popular tools for clonal deconvolution that use this framework. `pyClone` and `DPclust` implement Binomial mixtures, and `pyClone` supports also Beta-Binomial ones; in both cases a stick-breaking construction for Dirichlet Process priors is adopted^46^. `PhyloWGS`, instead, combines Binomial distributions with a tree stick-breaking construction for the Dirichlet Process priors^47^. Because of that, `PhyloWGS` can jointly cluster the input SNVs and construct a phylogenetic tree for the detected clones.

An alternative popular approach based on finite mixture models is `SciClone`^7^, which supports Binomial, Beta and Gaussian mixtures. SciClone fits the models to data via Variational Inference, an information-theoretic approach to approximate the posterior distribution over the model’s parameters. `SciClone` is a hybrid tool, as it can cluster allelic frequencies via Beta/ Gaussian mixtures, and read counts via Binomial mixtures. We want to note that, with Beta distributions, canonical Bayesian modeling leads to intractable priors, even if the conjugate prior distribution of the Beta distribution can be found by following the principles of conjugate priors for the exponential family. For this reason, Variational Inference of Beta mixtures exploits a Gamma approximation to the prior and posterior distributions, originally derived by Mao and Li^48^. In this approximation we cannot derive the so-called expected lower bound, a standard measure to monitor convergence of a fit.

These models are related to `MOBSTER`’s framework: they assume that *ρ* can be approximated by a point-process (e.g. a Dirac distribution) centered at the Beta means. The potential pitfall is clear: by applying the observational process to SNVs that include tail(s), the number of clones is overestimated. Clusters will be called from tail’s SNVs, which is contradictory because we look for clones under selection. To be precise, SciClone with Beta distributions models the frequency spectrum as well; however, that method does not include power-law frequencies, leading to a similar overfit.

#### Implementation and fit

`MOBSTER`’s implements a statistical model that fits the frequency of the variant allele corrected for copy-number status and purity, for *n* SNVs. Thus, we expect the clonal peak to be at roughly 0.5 VAF, with outliers spreading around 0.5 but below 1. We prefer these values to the so-called Cancer Cell Fraction values, which would require truncation above 1 to use Beta distributions^3^. We stress that, it is important to use SNV s in high-confident diploid regions for this type of analysis because miscalled copy number states might confound the inference (via artifact clusters of mutations).

`MOBSTER` can map SNVs to *Y*, the tail, and to any of the *B_i_*, the clones. Once one has fit a model to data, tail SNV s can be removed, and other methods can be used to fit the observational process on the read counts of the remaining SNVs. For this reason, `MOBSTER` is complementary to the tools mentioned above, as it works upstream the observational process. Nonetheless, our method provides also a preliminary indication on the possible number of subclones in the tumor.

#### Distributions and likelihood

The fit uses a pre-specified number of *k* + 1 components, where:

- *Y* is a Pareto Type-I distribution to capture the power-law tail. For a scale *x*_*_ and shape *α* > 0, its density is

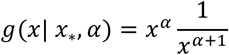

for *x* > *x*_*_, and 0 otherwise. Notice that the density is 0 for values below the scale parameter, and that its support is [0, +∞).
- *k* Beta distributions *B*_1_,…, *B_k_* to model clonal/subclonal clusters. For a shape *a* > 0 and *b* > 0 the density of a Beta random variable is

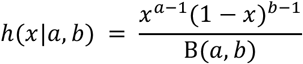

where 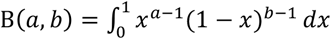 is the beta-function. The support of this distribution is [0,1], the full frequency spectrum.

The overall model uses a Dirichlet prior on the abundance of each clone; thus `MOBSTER` is a Finite Dirichlet Mixture Model with Beta and Pareto distributions. The model likelihood for a dataset *X* = {*x_i_|i* = 1,…,*n*} where we assume each *x_i_* to be iid, is a combination of two types of densities

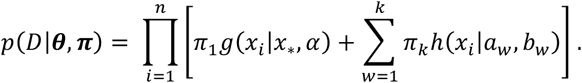

We use ***θ*** as a shorthand to the model parameters, and ***π*** = [*π*_1_…*π*_*k*+1_] for the mixing proportions – a standard Dirichlet variable on the (*k* + 1)-dimensional probability simplex. Notice that, just for notational convenience, we are assuming that the first model component is the Pareto random variable (the tail).

#### Fitting MOBSTER

The formulation uses *n* × (*k* + 1) latent variables ***z***; the plate notation is in Supplementary Figure S1. A variational approach to fitting this mixture is possible: we could use conjugate Gamma priors for the Pareto, and we would approximate the posteriors for the Beta components as in `sciClone`. However, we would could only approximate a criterion for convergence of the fit, as explained above.

We prefer to fit the model parameters via Maximum Likelihood Estimation (MLE) through an adaptation of a standard Expectation-Maximization approach (EM). This provides a solid alternative and a faster solution compared to a Monte Carlo sampling strategy for a Bayesian approach. We perform these standard steps:

- <u>E-step:</u> posterior estimates of the latent variables are defined as usual, once we account for the two different distributions involved

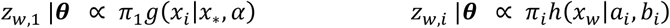 In both cases the normalisation constant *C_w_* is the overall density mass for point *x_w_*

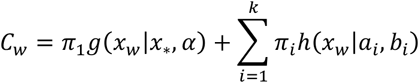
- <u>M-step:</u>

∘ <u>Pareto tail</u>. We begin by noting that the scale *x*_*_, of the Pareto distribution can be set to its MLE estimator, the smallest observed frequency *x*_*_, = min *X*^49^. This is a constant of the data, so we have one less parameter to fit. We fit the Pareto shape *α*, given *x*_*_; switching to the log-likelihood and including latent variables its MLE estimator is

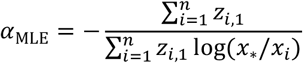
∘ <u>Beta clones</u>. The MLE estimator for Beta distributions has no closed form, but we can approximate it numerically. This can slow down the fit; however, we can rely on a recent result by Schröder and Rahmann on mixture models of Beta distributions^50^ concerning the alternative Moment-Matching (MM) estimator, which is analytical. MM consists in matching *t* empirical moments of the data *X* to the theoretical moments of the distribution, and solving for them. Here *t* = 2 (so mean and variance); a Beta distribution has mean *μ* and variance *σ* given by

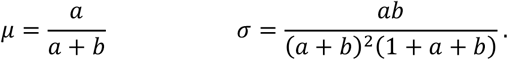 For a Beta, conditioned on the latent variables, the MM estimator is

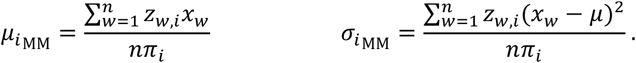 Given estimates for *μ_i_* and *σ_i_*, we can re-parametrise the Beta as

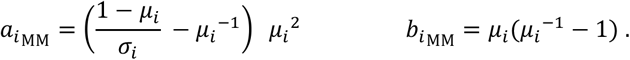

We remark that MM is not the same as computing the MLE, which computes the zeroes of the derivative of the likelihood with respect to the parameters, *∂h/∂****θ***. Thus, the properties of standard EM do not hold when we compute updates via MM: we cannot guarantee that the likelihood increases monotonically, because we cannot employ Jensen’s inequality. It is however shown in ref^50^ that the differences between the estimators are negligible in most cases. For the sake of precision, Schröder and Rahmann propose to call a fit through the MM for Beta distributions the “iterative method of moments”, rather than EM.

In `MOBSTER`’s implementation we provide both a standard EM fit with numerical solution for the MLE of Beta distributions, and the faster iterative method of moments. In the former case we monitor convergence of the likelihood, as standard. In the latter we use the posterior estimates of ***π*** since the likelihood is not monotonically increasing. A theoretical property of this MM approach is that, in each step, before updating the component weights, the expectation of the estimated density equates the sample mean. In particular, this is true at a stationary point; a proof of this is in Lemma 1 of Schröder and Rahmann^50^.

#### Initial conditions

As standard in EM approaches and variations thereof, we compute the fit with several random initial conditions. We provide two heuristics to compute the initial condition of the fit (Supplementary Figure S1).

- <u>Peak detection</u>. A simple peak detection heuristics in the frequency range [0.1,1] is applied to VAF values binned with size 0.01. To detect *k* initial peaks we perform kmeans clustering of each peak’s *x*-coordinate, and store their centres. If there are *w* < *k* peaks to cluster, we sample *k* − *w* random values in (0,1) for the remaining peaks. We use the centres of these clusters as mean of *k* Beta distributions with random variance in [10^−3^,0.25]; we sample variance values until the corresponding Beta parameters *a* and *b* are positive. For the tail, *α* is randomly sampled in the interval [0.01,5]. These values provide wide ranges of different initial distributions.
- <u>Randomized</u>. This procedure is as above, but we sample at random the Beta means and variances.

Experimental results show that peak detection is a more robust initialization method; the random counterpart sometimes leads to Beta distributions with mean approaching one, a region of parameter values where the likelihood becomes less stable, leading to numerical difficulties.

#### Clustering assignments and model selection

We do not want the fit to be biased towards tails, as we would miss low-frequency subclones that hide in the tail. Besides, simulations suggest limits to the detectability of tails. Thus, in general, given the quality of currently available NGS data, we cannot be a priori sure that the data supports the existence of a tail.

For this reason, in `MOBSTER` we can “turn off’ the Pareto component of the mixture and fit just *k* Beta. Hence we can perform model selection for *k* considering both models with and without a tail. This induces a statistical competition and allows us to select the model that best explains the data.

In `MOBSTER` we first compute the negative log-likelihood NLL = −log *f*(*X*|***θ,π***), which we use to derive AIC and BIC scores

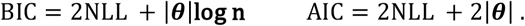

These criteria favor simpler fits by penalizing a model for the number of its parameters |***θ***|. A model with *k* Beta distributions and one tail has

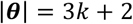

that break down as: *k* + 1 for the Dirichlet mixture ***π***, 2*k* for the Beta(s) and 1 for the Pareto tail (*α*). When we run the fit without tail, the model has |***θ***| = 3*k* − 1 parameters. Fewer parameters reduce the penalty of these scores, thus favoring fits without a tail.

In `MOBSTER` we want to drive the fit to select separate clusters, i.e., fits with few overlapping components. We achieve that by using two types of entropy terms. In one case we compute, from the output latent variables, the entropy H(***z***)

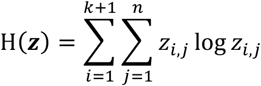

which leads to the standard Integrative Classification Likelihood

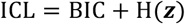

approximated through the BIC^22^. We also introduce a variation to the `ICL`, which we call `relCL`, a reduced-entropy criterion where we use the entropy of mutations that are not assigned to a tail (Supplementary Figure S1). This is defined as

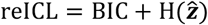

where 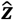 are the latent variables for the set of mutations {x|1 ≠ argmax ***z***_*x*,._}, re-normalized. Notice that 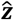 is defined from the hard clustering assignments that we use to assign mutations to clusters.

Entropy terms in `ICL` and `relCL` help to fit separate clusters because overlapping mixture components have higher entropy/ penalty. This happens since the uniform distribution has maximum entropy, which is when we cannot confidently assign mutations to clusters. By definition, `ICL` will prefer fits with a clear separation among tail *and* Beta components, while `relCL` will only require separation of the Beta components. This modification to the `ICL` seems reasonable: the tail overlaps to all subclonal clusters by definition, which in turn leads to excessive entropy penalization in standard `ICL`. For this reason, `ICL` will be more stringent in calling tails than `relCL`. See also Supplementary Figure S1 for a graphical explanation.

Notice that, because we are using NLL, we seek to *minimize* these scores. In the next sections we investigate also the different model-selection strategies. The default score for model selection in `MOBSTER` is `relCL`, which seems to provide a clean signal to identify the Beta components while retaining the tail structure.

### Simulation of cell tumor populations (1D and 2D)

We used two different simulators that use a stochastic branching process to simulate the growing population of a tumor. We have previously described the principle of both simulators and any additional modifications are described below. The code and scripts used to generate the synthetic sequencing data are released with the software material of this paper. We also provide an animation of tumor subclonal expansions as Supplementary Data (“Visualising subclonal expansions”). That simple animation shows how subclones emerge from low frequency up to their subclonal sweep (i.e., when they are detectable etc), how the VAF distribution changes over time, and how the fit of `MOBSTER` changes accordingly.

A non-spatial simulator^51^ was used to generate the data we used to measure the performance of `MOBSTER` in a univariate setting (Figure 3). The method allows the simulation of variant allelic frequencies (over time) of mutations accumulating during the growth of a tumor with a known clonal structure. The following smaller modifications were made. Instead of a Poisson distributed coverage *C_i_* of a mutant allele *x_i_*, an over-dispersed beta-binomial distribution was used. Given the averaging sequencing depth 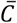 and a constant dispersion parameter *ρ* = 0.08, per-allele coverage *C_i_* values were determined as

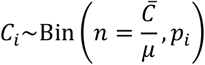

with *μ* = 0.6 and

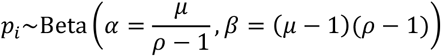

Variant allele frequencies values were assumed to be Binomial distributed, with the known fraction of mutated tumor cells in the population (*x*), given normal contamination (1 − *a*) and constant ploidy (*π* = 2)

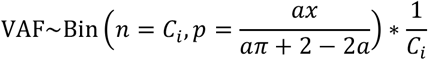

For the non-spatial simulations we simulated, similar to results from ref^51^, one ancestral population, with a single mutant subclone. The evolutionary parameters were set as follows, and kept constant through simulations: the tumor mutation rate *μ* = 16 (in mutations per cell doubling), the death rate *ω* = 0.2, the total number of reactions *t*_end_ ≈ 1.8 * 10^8^ (*N_cells_* ≈ 1 * 10^8^), average sequencing depth (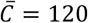 for a 120x simulated coverage) and number of clonal mutations *N*_clonal_ = 500. Nine random simulations for various subclone birth rates

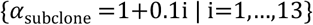

and number reactions prior to initiation of a subclonal expansion

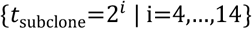

were produced. All simulations in which the subclone accumulated less than 50 mutations prior to its transformation (i.e. less than 4-5 divisions) were removed and three datasets with specific fraction of mutated cells in the population (*x*_subclone_) were generated by randomly selecting from the remaining simulations as follows:

- 20 effectively neutral cases where: *x*_subclone_ < 5%;
- 20 effectively neutral cases where: *x*_subclone_ > 90%;
- 110 cases with a detectable subclone: 20% < *x*_subclone_ < 80%.

These cases represent tumors with small subclones (almost undetectable), tumors where the subclone has swept through the overall population and cases where the subclone is detectable within the VAF spectrum.

To analyze the influence of sequencing depth as well as sample purity two additional datasets with variable purity (*a* = {0.3,0.6,0.7,0.9}) or variable depth 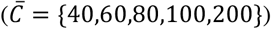 and otherwise identical parameters were created.

A second simulator model described in ref^15^ was used to generate several synthetic multi-region sequencing datasets to test the behavior of `MOBSTER` in a multivariate setting and to assess confounders (Figures 4 and 5). In brief, birth and death (reaction rates: *α_j_* and *ω_j_*) of cells on 2D square lattice were simulated via the Gillespie algorithm^52^. A cell selected to die was removed from the lattice. Daughter cell created during birth were either placed on a random empty neighboring grid point (Moore neighborhood) or if no empty adjacent grid point existed, on a neighboring grid point that was freed by pushing adjacent cells into a random direction (**v**) up to a given distance (*d_j_*). The number of additional mutations introduced into the genome of a daughter cell was drawn from a Poisson distribution with mean given by the mutation rate (*μ_j_*). New subpopulations (*j*) were inserted at given time point (*t_j_*) by modification of a random member of a selected subpopulation. Simulation of next-generation sequencing for each bulk sample (squares on the lattice) was done as described above for the non-spatial simulator.

We crated synthetic datasets in which 2-9 samples were taken from ten synthetic tumors with one (*n* = 50), two (*k* = 10) or three (*n* = 10) subpopulations with increasing fitness (birth rates). Simulations were ended when any of its cells reached the edge of the 800×800 2D lattice (≈ 5×10^5^ cells). Time points at which a new subpopulation was introduced (*t*_clone_ = {0,4,6.7}) and corresponding birth rates (*α* = {1,1.6,2.4}) were chosen to allow coexistence of each subpopulation at a approximately equal abundance at the end of the simulation. The remaining parameters were kept constant: *μ* = 10, *N*_clonal_ = 100, 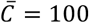,*ω* = 0 and *d* = 100. Bulk samples of about 10,000 cells (100×100) were taken along the outer perimeter with an equal angular distance relative to the centre between them.

### MOBSTER’s analysis of synthetic data

We used synthetic data for *n* = 150 non-spatial (i.e., 1D) tumors to measure how tails confound subclonal deconvolution (30 cases of neutral tumors with 0 subclones, 120 with one subclone). All tests have been carried out for various configurations of simulated mean coverage, and purity; we consider the ideal performance at 120x mean coverage with perfect tumor cellularity (purity 100%). Input sequencing datasets have been created as described in the previous section. To run `MOBSTER` we adjust the observed allelic frequency. Simulated mutations have no coy number associated, and we consider all simulated mutations to occur in diploid genome regions. We remark that copy number events are another confounder that acts in these analyses, as the shape of the adjusted VAF distribution depends on the correction by copy number status of each mutation. However, since here we are interested in the identification of subclonal populations from the VAF per se, we omit to include other confounders due to noisy copy number estimates in our analyses. The formula for the standard adjustment of allelic frequencies reads as

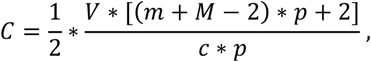

where *V* is the raw VAF in the data (ratio between depth of reads with the mutant allele, over overall depth at the mutant locus), *m* and *M* the minor and major copy number of the mutation, *p* is sample purity, and *c* are the copies of the mutant allele. This is half the value of the Cancer Cell Fractions, and thus the clonal peak is expected at 0.5 instead of 1, which allows using Beta distributions that range in [0,1] for clonal peaks. This value might be properly called the adjusted VAF for a diploid mutation in the tumor; in our simulated tumors *k* = *m* = *M* = *c* = 1, and thus *C* = *V/p* as expected (adjustment for purity). Example fits with simulated data are shown and commented in Supplementary Figures S2 and S3.

In Supplementary Note 1 we explain all the parameters that we have adopted in these simulations, which we used to analyze several output metrics for clustering precision and sensitivity. In particular, we measured:

1. The number *k* of clusters that fit the data, finding `MOBSTER` highly accurate. Genuine errors of the tool are often due to particular idiosyncrasies of the data. This is expected, as this clustering problem is very difficult: by construction tails and clone’s peaks overlap (mixtures overlap), and hence a weak signal of selection might be difficult to detect from the VAF spectrum (Supplementary Figure S4).
2. The confidence in the prediction of a tail, via log odds between competing statistical models, and found that when `MOBSTER` fits a tail to the data, the odds ratio is much higher (i.e., there is more evidence towards that model, Supplementary Figure S4). This shows that a tail improves the fit of the data, from a statistical point of view.
3. The fit precision from the rates of true positives and false negatives, and from the Euclidean distance between each predicted and its closest true peak. We found very good rates and lower distance for `MOBSTER`, which both decreased when we fit a tail (Supplementary Figure S4).
4. The effect of coverage (40x, 60x, 80x, 100x, 120x and 200x) and purity (0.3, 0.5, 0.7 and 0.9) on the inference of the clonal structure, and found limits to the ability to detect tails with coverage below 100x/ 120x at very high purity (>0.9); Supplementary Figure S5.
5. `MOBSTER`’s ability to call subclones with different size – number of SNVs – and peak position, and the power to distinguish small subclones from tails. We found that `MOBSTER` can fit both the subclone and the tail for a wide range of parameter values, but that the overlap of tails and subclones complicates the inference, as we might expect (Supplementary Figure S6).
6. The lack of bias in the method. This is assessed showing that, in tumors without tail, `MOBSTER` correctly identifies a small subclone via a Beta component, if its peak is above the minimum value for detectability (Supplementary Figure S6).
7. The effect of model selection strategies via BIC, `ICL` and re`ICL`. We found `ICL` and re`ICL` to be more suitable to properly balance the number of clones in the data. This is expected because BIC does not account for the mixtures overlap, which is an issue in this particular clustering problem, while `ICL` and re`ICL` do so by using the entropy of the latent variables. Because `ICL` is more stringent in calling tails, we used that to draw these conclusions about `MOBSTER`’s performance (Supplementary Figures S7).

See Supplementary Note 1 for a detailed description of all the simulated tumors and data to support these conclusions. We also compared the effect of using `MOBSTER` before analyzing read counts for non-tail mutations, as opposed to a direct analysis as commonly done in the field. We tested

1. Two core statistical methodologies for clonal deconvolution (Dirichlet Processes with Monte Carlo sampling, and Binomial Finite Mixtures fit via variational methods). The multivariate variational method with Binomial distributions has been implemented in `MOBSTER` (Supplementary Note 2).
2. Popular tools for the problem based on the above methodologies, as described in the Main Text.

We have observed that the overfit happens at the core of the statistical methodology, leading to systematic errors without `MOBSTER`. We have also assessed the role of a parameter which largely affects the number of output clones predicted by this method: the concentration *α* > 0, for which we scanned point estimates (1,10^−2^,10^−4^,10^−8^), and a Bayesian Gamma distributed *α* ~ Γ(0.01, 0.01). The error in all these tests is very large (overfit) without `MOBSTER`. For tumors without subclones and high concentration we call *k* =4 Binomial clusters (3 subclones), where there should be 0. The error persists but is diminished for lower *α*, and the trend is the same regardless the number of subclones, which suggests that `MOBSTER` improves the analysis. The full set of results, which extend the ones showed in the Main Text, is in Supplementary Figure S8.

For multivariate analyses we used the spatial 2D simulator described above (simulated tumors with 0, 1 and 2 subclones). We randomly sampled from 2 to 9 biopsies of the same simulated tumor, located evenly spread across the simulated tumor so that some biopsies might fall on the overlap between two geographically distinct subclones, while others might be monoclonal. For this test we wanted to focus on spatial confounders, and then fixed the mean sequencing coverage to 100x, with purity 1 for all samples. We compared the fits from a multivariate variational Binomial clustering method on read counts of all data, and on the subset that passed `MOBSTER`’s analysis. The latter is computed running `MOBSTER` first on each simulated biopsy, and calling “tail mutation” any SNVs detected at least once in the tail of a sample. In this way we avoid calling clones in tails, and the observed performance is genuinely reflective of the effects of sampling a spatially heterogeneous tumor.

To aid the explanation of the confounders discussed in the Main Text, we used a “virtual staining” method: to stain for a set of SNVs, we color all cells that have one or more such SNVs. Each cell is colored by its true clone’s color, thus clonal SNVs provide a representation of the simulated diffusion of clones in space, while private subclonal mutations represent wedges of such plot. With these plots, it is also straightforward to assemble the clone tree for complex scenarios by nesting staining plots; see for instance Supplementary Figure S9 for an example tumor with 2 subclones. We focused the analysis of these simulations to estimate the clone tree that we can infer with and without `MOBSTER`. This seems particularly important as the overfit of clones due to spatial confounders and neutral tails can largely increase the number of possible trees. The results discussed in the Main Text are confirmed even if we divide tests by number of biopsies and subclones simulated (Supplementary Figure S10, all the points lay in the upper diagonal of the plot). The larger number of trees follows from the larger number of clones fit by these two methods; without `MOBSTER`, extra low-frequency clusters of subclonal mutations can be positioned in different branches of the clone trees, thus increasing the uncertainty over the generative model.

The strategy that we propose in order to minimize the effect of these confounders is heuristic. When for each Binomial cluster we have computed its parameter (peak) for each sample, we retain only those that have value above a threshold *x* in at least *y* biopsies (for instance *x* = 0.05 in *y* ≥ 2 biopsies). This is an intuitive heuristic which imposes a certain amount of empirical evidence to call a subclone; the parameters *x* and *y* should be set based on the study and the data under scrutiny. With this method, `MOBSTER` results are more confident and less noisy than the ones that we can obtain without. The median error reaches 0% (top score) as measured in violations of the pigeonhole principle according to the clone trees generated by REVOLVER, a previously published method for clone tree inference from cancer cell fractions and subclonal deconvolution outputs^24^. Briefly, from the output clusters the method computes all possible clone trees that fit each sample independently, and assemble the detected edges in a graph. Then, it scans the graph to detect all possible spanning trees rooted in the clonal cluster, and ranks trees using a scoring method for the number of violations of the pigeonhole principle (perfect score 1 has 0 violations). A violation for a sample is when, in a branch *x* towards *y* and *z*, the observed adjusted VAF (here ½ * CCF) of *x* is lower than that of *y* plus *z*. Interestingly, we observe that the strategy that we propose to identify clones is less effective without `MOBSTER` because tail mutations that spread across samples tend to form clusters that co-occur in all samples, and thus no reasonable value of *y* can identify them. These analyses suggest that evolutionary interpretations derived from the structure of clone trees should take into account the effects of tumor spatial sampling bias. In general, these analyses also highlight that mutations that derive from neutral processes should be removed with `MOBSTER`; otherwise one calls clusters of alleles that do not reflect true forces of positive selection.

We also used `MOBSTER` fits to measure quantitative evolutionary parameters of tumor growth. We could do adapting the computations originally carried out in Williams *et al*. via similar principles^8^. When we fit a tail with `MOBSTER` we can measure the tumor mutation rate *μ* scaled by the probability of lineage survival *β*

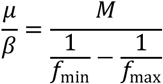

where *f*_min_ is the minimum VAF and *f*_max_ is the maximum, and *M* is the number of mutations between *f*_min_ and *f*_max_. The ranges of the VAF distribution can be taken from the posterior assignments of the latent variables that map mutations to the tail (if any), to avoid outliers we use the mutations within certain empirical quantiles (for instance, 2% and 98%). The time unit of this rate are tumor-doubling times, and the conversion to rate per base pairs can be computed dividing this value by number of sequenced nucleotides; for a whole diploid genome this conversion factor would be 3 * 10^9^. With the mutation rate and the fit parameters of each subclone we can calculate the time the subclone emerges, and its selection intensity. Selection *s* > 0 is defined as the relative growth rates of host tumor cell populations versus the subclone. See our earlier work for a detailed explanation of the formulas to derive the selection coefficient and emergence time for a subclones^8^.

### Analysis of patient derived data

We obtained WGS mutation calls for the breast cancer sample PD4120a and the AML Platinum sample from our earlier manuscript^8^; originally these two samples have been discussed in two separate publications (cfr^3^ and cfr^29^). For PD4120a we used SNVs from a highly confident diploid chromosome (chr3), for Platinum we used whole genome data, consistently with previous works^8^. Both datasets are the highest-resolution single-sample WGS studies available so far in the public domain (200x and 320x); comparatively, samples from large-scale studies unlikely hit the minimum limit (>100x) suggested in our study to detect reliable subclonal structures. For both datasets we used only annotated SNVs with adjusted VAF above 0.05%, and run `MOBSTER` and sciClone with default parameters. From the clusters we computed manually the possible clonal trees via the pigeonhole principle. `MOBSTER` analyses these tumors in about 5 minutes on a standard multicore high-performance computing environment (parallel implementation).

We measured the evolutionary parameters for both the breast and AML patients, and found estimates that are in agreement with the analysis by Williams et al., which used Approximate Bayesian Computation to fit a stochastic branching process of tumor growth^8^. The parameters identified are reported in the Main Text.

For Set07 and Set06, a colorectal carcinoma whole-genome sequenced at median coverage 100x, we adopted the same pipeline described in Cross *et al*.^30^. Notice that the original analysis referred to these two patients as “carcinoma 6” and “carcinoma 7”. Compared to the original analysis of these patients, these new data have higher resolution (100x versus ≈40x). Briefly, we used Mutect and CloneHD to call somatic mutations and copy number segments across all samples (copy number calls were also double checked with Sequenza, data not shown); sample purity was estimated from clonal diploid mutations. The purity of most samples is around 80%; for Set07 values per sample are 0.88, 0.88, 0.88 and 0.80, for Set06 they are 0.66, 0.72, 0.80, 0.80, 0.80, and 0.80. To use `MOBSTER` we computed copy number and purity adjusted allelic frequencies. Reliable diploid regions are called from segments with minor and major copy number equal 1, excluding 10 megabases around the centromere of each chromosome (Supplementary Figure S11 and S15); this cleans the signal in centromere areas that are difficult to align. We retained SNVs and adjusted the observed VAF of each sample, and checked quality of the calls; to reduce false positive calls and contamination from germline mutations, we retained only putative somatic mutations called privately to each one of the patients that we analyzed (to implement this filter, we also used data from a third adenoma sample available in Cross *et al*.^30^). Then, we imposed other filters on the data, removing all SNVs with adjusted VAF below a 5% cutoff adjusted for sample purity, and removed those with VAF above 0.7 (70%) as they correspond from miscalled diploid segments. These filters altogether identify good quality reliable somatic mutation calls; besides, by using a whole-genome sequencing protocol, after the filter we still retain a substantial number of SNVs to perform subclonal deconvolution (≈ 80,000 in total, ≈ 50,000 for Set07 and ≈ 30,000 for Set06). In Supplementary Figure S12 (Set07) and S16 (Set06) we show several views of the data distributions for VAF values and coverage in all the sequenced samples.

From the remaining SNVs, we run `MOBSTER` with default parameter and `ICL` model selection, which is more stringent in calling tails and subclones. Each run of `MOBSTER` converged on a solution with 0 subclones (*k* = 1) and subclonal mutations assigned to a very evident power law tail. The tails fit for these data are very large and suggest the lack of positive selection at the subclonal level, for these colorectal cancers. This observation is also in line with the preliminary analysis of these patients (Figure 2 in Cross *et al*.^30^) which used lower resolution data (mixed whole genome and whole exome) and sample trees – a classical type of phylogenetic tree with tumor samples as leafs of the tree – to show that there was no evidence of positive subclonal selection in these tumors. We used another method, based on different premises, to assess how likely that there is no subclone under positive selection under the tail of these tumors. We used the dN/dS method from the dndscv tool by Martincorena *et al*. ^31^ to estimate the ratio of non-synonymous to synonymous substitutions in these tumors; we computed the dnds value for both patients independently, and pooling the data of both patients altogether (as usually done in these analyses). We have split mutations to compute the dnds statistics into two groups: the first group are those that are called clonal by the Binomial clustering after `MOBSTER`, which in both cases are those assigned to cluster C1. The remaining mutations are assigned to the other group; these are then the union of the mutations that are assigned at least once to a tail by `MOBSTER` (tail mutations), and the ones that we remove with our heuristic to prioritize subclones from Binomial clustering after `MOBSTER`. The presence of possible positive selection is reported by estimated values strictly greater than 1 (dnds > 1); neutral mutations should have dnds ~1, and mutations under negative (or purifying) selection show dnds values below 1. For each estimate, the method returns both a point estimate of the dnds value, as well as a 95% confidence interval. In order to achieve statistical power for this analysis we used all the genes available in each patient, because there are no enough mutations in cancer genes with just 2 patients. For pooled data and clonal mutations we find a value above one (dnds = 1.55; CI [0.91, 2.65]) as expected, which breaks down as (dnds = 1.78; CI [0.87,3.6]) for Set_07 and (dnds = 1.29; CI [0.58, 2.8]) for Set_06. This analysis confirms the lack of evidence of positive subclonal selection; in fact for the remaining mutations we find a pooled dnds value below or almost equal to 1, and with narrow confidence intervals (pooled dnds = 0.85; CI [0.70, 1.02]; Set_07 dnds = 0.95; CI [0.76, 1.20] and Set_06 dnds = 0.68; CI [0.50, 0.92]). Interestingly, the observation of negative selection forces, hereby suggested by dnds values below 1, has been recently linked to power-law neutral dynamics for the clone size distribution^53^. Summarizing, this analysis provides further evidence that joint analysis with `MOBSTER`, as well as our heuristics should not have discarded possible subclones from the available data. However, a remark for this analysis and its results is important. Usually to achieve statistical significance with these analysis one needs to pool together data from several patients to reduce the size of confidence intervals. In this case with just 2 patients, even when we pool together all data we do not have many coding mutations and the upper value of the confidence interval contains 1 – precisely, the value is 1.02 for pooled dnds of non-clonal mutations. However, the trend of the results suggests that the values are quite far off from being above 1 and, on top of this, the confidence interval of the dnds value for Set_06 alone is included below 1. We also not that similar values of point estimates and confidence intervals are obtained also with the dnds method by Zapata *et al*.^32^, which uses slightly different computations to estimate the dnds statistics (data not shown). We also checked for the presence of somatic driver mutations in the VAF spectrum, and found only the same set of driver events already identified by Cross *et al*.^30^. These are somatic mutations (SNVs and indels) in well-known colorectal driver genes (via Cosmic): for patient Set_07 we find mutations in *APC* (p.R787X and p.R1432X), *KRAS* (p.G12D), *SMAD3* (p.Y42X) and *TP53* (p.E159X), while for patient Set_06 we find mutations in *APC* (p.R216X), *KRAS* (p.G12V), *PIK3CA* (p.C420R), *ARID1A* (p.W1453X) and *TCF7L2* (indel). Some of these mutations are not SNVs, or happen to be found in non-diploid regions (see also^30^); for such mutations we use the model fits to map, a posteriori, the mutations to the clusters detected by our analysis. A remark is due for the presence of the *PIK3CA* mutation, which is annotated as driver in the trunk of the tree of Set_06. Cross *et al* have annotated that mutation originally as clonal^30^; in our data we find that to be part of the clonal cluster in 5 out of 6 samples, and tail in one other (in Set6_42). For this reason, in our analysis the mutation is actually removed from the successive Binomial clustering step. Nonetheless, that mutation in Set6_42 has adjusted VAF ~30%, which places the mutation slightly below the point of crossing of the tail and the clonal cluster. This is also reflected by the posterior latent variables that we use to assign the mutation to the tail (~80% probability) and the clonal cluster (~20% probability). The mutation is clearly clonal in all the other biopsies (adjusted VAF > 40%). However, the purity of Set6_42 is well below the purity of the other five samples (~66% versus ~80%), which makes it possible that this reduced VAF is also due to excessive contamination of normal tissue in this sample, and consequent higher noise in the sequencing data. For this reason, we preferred to annotate this mutation as clonal in this tumour. It should be evident that this discrepancy does not affect our analysis, and that our conclusions are not driven by this difference. In particular, in the dnds analysis – which is the only one that might be affected by this clonal/ subclonal classification – we have assigned the *PIK3CA* mutation to the set of tail mutations as reflected by our analysis. This ensures that we do not to bias the reported dnds statistic in favor of neutrality, which is a key point of our analysis.

To compute the final clusters we used `MOBSTER`’s variational Binomial method on the raw read counts of non-tail mutations. The output clusters have been filtered as suggested in the Main Text, imposing a minimum number of observations (2) for each subclonal cluster above a minimum VAF value (subclonal peak 0.02). In both patients we find – as expected – private subclonal mutations in each biopsy sample, and a very small subclonal cluster (which is called C5 in both analyses). This cluster contains mutations that are clonal in some of the available biopsies, and thus cannot be excluded by our heuristic. However, we still decide to remove this cluster because its size is largely below what a standard analysis would have considered suitable (usually, 5% of tumor size); precisely, the amount of mutations assigned to this cluster account for ≈1% of the tumor load in Set_07 and ≈ *2%* in Set_06 (Figure 6), after `MOBSTER`’s tail removal – which means that with respect to the original tumor size this cluster would have had much smaller (≪ 1%) because in both cases tail mutations account for ≈ 80% of the tumor mass.

From the clusters we computed the possible clone trees, measuring their violations of the pigeonhole principle with the same method implemented in REVOLVER^24^ as done for the multivariate simulations described above. The same procedure has been repeated omitting to run `MOBSTER`, which would be the standard method’s analysis for these tumors. Notably, `MOBSTER` is very fast: the analysis of each patient takes about 20 minutes (parallel implementation of `MOBSTER` fits, the variational Binomial mixture and clone tree generation). The output trees are shown in Supplementary Figures S14 for Set07, and in Supplementary Figure S18 for Set06. In both cases, we find a much more complex clonal history when we do not use `MOBSTER`, because the confounders due to neutral evolution are not accounted for. In particular, fake small subclonal clusters confound the construction of the clone tree, as they can be attached to different internal nodes leading to potential violations of the pigeonhole principle. This is consistent with results from simulations in the multivariate setting, as we have described above. In many cases, we have to allow for violations of the principle otherwise, without `MOBSTER`, we never find a tree to fit the data; in other words, the best tree has at least one violation. With `MOBSTER`, instead, the output trees can be trivially constructed because there are only 2 clusters that we prioritize – if we neglect C5, the clonal architecture would contain no edge at all, because the predicted tumor is monoclonal.

Beyond the structure of the clone trees, we are interested in how the clusters are assembled and the SNVs organized. This is fundamental, as clustering assignments are a proxy for the putative genotype of each clone in the tumor. For this reason, we have mapped the clusters that we obtain with `MOBSTER` to the ones that we obtain without, and viceversa. So doing, we could see how the mutations that one analysis assigns to a cluster (say to cluster *x*), are assigned by the other analysis. Ideally, if the two analyses were perfectly concordant, every cluster should map to only one other cluster. Otherwise, the analyses are not clustering the data in the same way, and we can try to investigate why. Very interestingly, from these mappings we can observe that tail mutations, when we do not use `MOBSTER`, spread to all clusters (i.e., they are assigned ubiquitously across clusters). This has profound implications and means that the confounder of neutral mutations has strong effects that drive the outputs of subclonal deconvolution. Some clusters, in fact, are just driven by tail mutation as we might expect but the most striking effect is that tail mutations largely contribute to each one of the output clusters. From a data point of view, this can be explained observing that genuine tail mutations largely overlap (i.e., they are found at the same values of VAF) with non-tail clusters. For this reason, the statistical signal that determines each cluster cannot be decoupled in tail *vs* non-tail mutations unless one uses specific tools like `MOBSTER`. From these plots we also see explicitly that the clonal cluster determined without `MOBSTER` contains tail SNVs that are clonal in some biopsy, but subclonal in others. This suggests a complex scenario of intra-tumor spatial heterogeneity for these samples, and the effect of spatial sampling bias when we attempt a reconstruction without `MOBSTER`.

