## Supplementary Notes for "Model-based tumor subclonal reconstruction"

MOBSTER

#### This PDF file includes:

Supplementary text
Supplementary Figures S1 to S18
Caption for Database S1
References for SI reference citations

#### Other supplementary materials for this manuscript include the following:

Database S1

### Supporting Information Text

**3 Note 3: Software and availability 14**

### 1. Note 1: Analysis of single sample simulated data

Unless otherwise specified, we have run MOBSTER with these parameters: number of clones $k = 1, \dots, 5$ (fit with and without a tail, for every $k$), initialisation via peak detection and 10 independent runs for each parameters configuration, fit via iterative method of moments, with a cutoff at 6000 iterations, and stopping when the variation in the estimate of $\boldsymbol{\pi}$ is smaller than $\epsilon = 10^{-8}$. Thus, for each tumour, $5 \times 2 \times 10 = 100$ independent fits are compute; the best model is the one that minimises the score used for model selection (usually ICL or reICL) across all runs; we report explicitly the score adopted when that is not ICL.

**A. Example executions.** Example fits from a subset of the large scale test (120x WGS with 100% purity; ICL model selection) are shown in Supplementary Figures S2 and S3. In Supplementary Figure S2, MOBSTER fits the true model to tumours with and without subclones; peaks of Beta distributions are SNVs undergoing selection, while the others evolve neutrally as a power law tail in the VAF distribution. The subclonal deconvolution problem is difficult, and the true clonal structure can be mistaken for a variety of reasons that we summarise in Supplementary Figure S3:

1. when there is an almost total, but not complete, subclonal sweep:
   - here the fit correctly identifies the clones, but because the variance of the Beta distribution is very flexible, MOBSTER does not separate clonal from subclonal SNVs. This is not a problem because the subsequent analysis of non-tail read counts uses Binomial distributions. The variance of a Binomial component depends on sequencing coverage*, thus model selection for the number of mixture components can split the Beta mixture in two (or even more) close Binomials. This is not necessarily true if we analyse read counts with Beta-Binomial distributions that have flexible variance via overdispersion;

*A random variable Bin$(n; m, p)$ has variance $np(1-p)$ where $p$ is the peak and $n$ is the number of trials (coverage).

2. when a subclone is “hidden” by the tail:

   - This is a genuine error of MOBSTER. It is difficult to detect subclones that originate at a time close to biopsy collection, or that grow not much faster compared to the ancestor (slow subclonal sweep). In the former case, for instance, these subclones have a few mutations more than the ancestor (small subclone);

3. when there is a tradeoff between detecting subclones and tails.

   - If a large low-frequency subclone overlaps largely with a tail, the ICL score favours a Beta fit to the leftmost part of the frequency spectrum (no tail). In this case, the method will assign real tail mutations to the detected subclone, which is a minor inconvenient because we identify the subclone. The reICL score uses a reduced entropy to penalise for the overlap of subclones, achieving better intermix of tails and subclones.

#### B. Tail detection and fit precision.

**Large scale test with $150$ tumours (Supplementary Figure S4).** Here we'll comment on the analysis of $150$ tumours described in the Main Text, where in each run we measure several outputs to quantify the quality of fit with MOBSTER. These tests with ICL show that MOBSTER is very precise in estimating the peaks of the clones, as well as the tail.

- **Identification of the true $k$.** We measured $k$, the number of clusters fit (neutral tumours $k = 1$, non-neutral tumours $k = 2$, one subclone). Summary statistics show that MOBSTER is highly accurate. We paid particular attention to analyse $59$ cases in which we underestimate the true $k$, and investigated if these are associated to particular idiosyncrasies of the data, or the method. We found that in $\sim 17\%$ of cases the peaks of the subclone hides in the tail, and is undetectable (low frequency). In $\sim 70\%$ of cases, instead, there was almost a complete subclonal sweep, and MOBSTER fits one Beta component to jointly fit the clonal cluster and the subclone, as in Supplementary Figure S2. $\sim 13\%$ of the cases are genuine errors of MOBSTER.

- **Confidence in the prediction of a tail.** The ICL odds between competing statistical models is used to determine the proper fit with and without a tail, when it is computed from the odds of the highest-scoring models with and without tail. Values above one support a model with a tail, otherwise without. Results show that when MOBSTER fits a tail to the data, the odds ratio is much higher (i.e., there is more evidence towards that model). This clear trend of higher odds persists regardless the number of subclones in a tumours; this suggests that a tail improves the fit of the data, from a statistical point of view.

- **Precision of the fit.** We measured the precision of the fit in two different ways:

  1. as the rate of true positives and false negatives. These represents respectively the rate at which MOBSTER calls a true peak, and that at which it misses a true peak. In order to match peaks we use a tolerance $\delta_{\text{peaks}} = 0.05$; this means that we center an interval at the true peak value, and accept a fit within $\pm(\delta_{\text{peaks}}/2) = 0.025$ of this value.
  2. as the euclidean distance between each predicted peak, and its closest true peak. This measure gives us an idea of “how off” some peaks are when we create false positives. For instance, if we call a false peak where we should have fit a tail, a large value for this distance is observed compared, for instance, to when we miscall a cluster within VAF range where there are mutations under positive selection.

  Measurements show that both error rates and distances decrease when we fit a tail.

**C. Confounding factors for tail detection: coverage and tumour purity.** A fundamental hurdle to detect tails is the lack of proper coverage and purity. To assess this effect on MOBSTER we analysed $80$ tumours of the $150$ generated tumour, in $5$ configurations of parameters (total $480$ tumours, Supplementary Figure S5). For each tumour we simulated:

1. mean coverage at $40\times$, $60\times$, $80\times$, $100\times$, $120\times$ and $200\times$, perfect purity, and accepting simulated SNVs with at least 6 variant reads. This accounts for a $5\%$ cutoff at $120\times$ shown in Supplementary Figure S4;

2. tumour purity spanning $0.3$, $0.5$, $0.7$ and $0.9$, at median coverage $120\times$.

Results varying the coverage highlight that it is difficult to fit a tail with less coverage than $80\times$. Tails, even with perfect purity, emit a clear signal for coverage above $100/120\times$. These considerations are reported by analysing both the size of the tail (i.e., number of SNVs assigned), as well as the overall number of tumours fit with a tail. Analysis of different configurations of purity shows a similar trend, as both confounders impact on the detectability of low frequency SNVs for the tail. In particular, for purity below $0.7$ tail's detection is impractical even at $120\times$. At purity above $0.9$ the tail signal is instead clear, and MOBSTER can retrieve the true model.

These results are important as they explain why many studies fail to observe tails in the data,: many large scale whole genome sequencing cohorts have coverage below $100\times$, with samples with different purity levels. Thus the data resolution is still too low, and forthcoming studies will have to overcame these limitations.

**D. Identification of subclones.** One crucial feature of a subclonal deconvolution algorithm is its abiilty to identify subclones. Besides the tests discussed above, we carried out two other types of tests to measure the performance of the method.

**General results on the identifiability of subclones (Supplementary Figure S6).** We have used MOBSTER's statistical model as a generative process of the frequency spectrum. So doing, we could create complex VAF scenarios and assesses the method's ability to call subclones with different size – number of SNVs – and peak position. In this test we simulated a clonal cluster with 400 SNVs, and a tail with 500; we randomly sampled the shape of the Pareto tail. Then, we simulated a subclone spanning from very low frequencies to large – i.e. from 0.05 to 0.23 – and with a number of SNVs that spans from $50$ to $1250$. We then measured the probability of fitting a tail with ICL, as well as the probability of fitting the subclone with 10 independent runs per configuration of parameters.

Results show that MOBSTER can fit both the subclone and the tail for a wide range of parameter values, but that the overlap of tails and subclones complicates the inference, as we might expect. In particular, in some configurations the subclone might be difficult to detect, as explained above. When we inspect the average number of Beta components fit by MOBSTER we confirm this intuition. It is extremely difficult to call a subclone with $< 250$ mutations if the tail has $500$, or if it has a peak below $0.10$, where the minimum observed VAF is $0.05$. In these cases, it is almost impossible to detect a subclone that hides perfectly under the tail. In these pathological cases the data harbours a very weak signal of subclonal selection, in all remaining cases the subclonal peak is clear and the method can retrieve it.

**Fitting tumours that do not have a tail (Supplementary Figure S6).** Through a direct simulation of the VAF distribution we assessed that MOBSTER is not biased towards "calling" tails with ICL. We simulated a tumour with 2 clones (Beta components), and no tail. The VAF distribution has only two bumps for to the clonal cluster and the subclone. A tail might be missing with low purity, coverage etc. We analysed only SNVs with allelic frequency above $5\%$, and simulated a subclonal peak spanning $0.05$ to $0.2$, with increases of $0.02$. This mimics the positioning of a subclone that is growing out of the tail: we want MOBSTER to fit it with a Beta, not with a tail. Results suggest that the subclone is always detectable when its peak is above $7\%$ (Beta fit). When the subclonal peak hits the detectability limit, instead, the observed distribution of the subclonal mutations splits into two symmetrical shapes that decay like a power law. In this case they are undistinguishable from a tail, and MOBSTER mistakes the subclone. These general results show that the method is largely reliable, and that it can correctly disentangle tails from subclones with suitable data.

**Model selection strategies (Supplementary Figures S7).** From the overall set of simulations analysed in Supplementary Figure S4, we performed model selection by using the BIC, ICL and reICL scores. Results confirm the principles underlying these methods:

- BIC does not minimise overlapping mixture components, and consistently calls two clones;.
- ICL is the most stringent method to call tails, which are often dropped for a Beta subclone when the signal of selection is strong enough. This is achieved by minimising the entropy among the tail and the clones;
- in our simulated tumours, reICL always calls a tail and subclones. This is because its reduced entropy term penalises only the overlap among Beta components, disregarding the tail.
- ICL and reICL are similar and more precise than BIC. reICL calls tails that overlap with subclones, which is reasonable when we want to "clean" the signal from neutral mutations, and call subclonal peaks in the VAF distribution.

**E. Comparison with popular Bayesian methods.** Our analysis shows that if we directly cluster data of tumours with a tail, then we overfit the number of clones regardless what method we use to approach subclonal deconvolution.

***Preamble: Bayesian methods for Binomial mixtures.*** At the core of widely used tools like pyClone, DPclust, PhyloWGS, and SciClone (1–4) there are two Bayesian statistical methodologies for mixture modeling (see (5, 7) for a thorough introduction):

- non-parametric Dirichlet Processes that describe distributions over distributions (here over mixtures of any size); posterior estimates for those models are computed via Markov Chain Monte Carlo (MCMC) sampling;
- semi-parametric Dirichlet Finite Mixture Models for a mixture of a finite number of Binomial distributions; posterior estimates for these models are approximated via Variational Inference (VI).

We focus on how these methods determine the number of output clusters $k$. Both models use a parameter $\alpha$ to model the propensity at which a new cluster is created during posterior computations; this parameter can be set either to a predefined value (point estimate), or learnt from the data via Bayesian computations. The interpretation of $\alpha$ is according to adopted model, and both methodologies estimate $k$ from data.

- with Dirichlet Processes (DP), parameter $\alpha$ – sometimes called concentration or scaling – appears in well-known generative models (e.g., the stick-breaking process, the Polya urn or the Chinese Restaurant process). The DP is a distribution over Binomial mixtures, and a draw from the DP is a full mixture distribution. Because drawing a concrete distribution is unfeasible, the constructive approach is often adopted. The foundation of these constructions exploits exchangeability theorems for random variables (5). If we assume the input observations to be analysed in a specific order, the behaviour for a new observation $x_n$ given the previous $n-1$ is to draw a new sample from a baseline measure $G_0$ with probability

$$x_n \mid x_{n-1}, \ldots, x_1 \sim \frac{\alpha}{n-1+\alpha}\,.$$

A new draw generates a new cluster and fixes its parameters according to some prior distribution $p(\boldsymbol{\theta})$; with Binomial components the sample is a new success probability $(p)$, from a conjugate Beta prior. Thus, $\alpha$ drives the fit of $k$: in the limit behaviour $\alpha \to 0$, the realisations are all concentrated at a single value and $k = 1$. Instead, if $\alpha \to \infty$ the realisations become continuous, and infinite clusters are created. This is why these models are non-parametric. We observe that, in practice, these models are semi-parametric because the number $n$ of observations in the dataset is an upper bound to $k$. The theoretical construction, however, is truly non-parametric as $k$ can grow to infinite for $n \to \infty$.

We want to highlight that in many implementations of Dirichlet Processes, posterior estimates of $\alpha$ are computed from the data via a conjugate Gamma prior $\alpha \sim \Gamma(a, b)$ with hyper-parameters $a$ (shape) and $b$ (rate). Its density is

$$\mathsf{Gamma}(x \mid a, b) = \frac{b^a}{\Gamma(a)} x^{a-1} \exp\{-bx\} .$$

where $\Gamma(z) = \int_0^\infty x^{z-1} e^{-x}\, dx$ is the gamma function. Notice that sometimes (e.g., in pyClone), the Gamma density is equivalently expressed through a scale parameter which is the inverse of the rate.

- with Finite Mixture Models fit via variational inference, the statistical model uses a Dirichlet prior on the mixing proportions. Assuming $k$ components

$$\boldsymbol{\pi} \mid \boldsymbol{\alpha} \sim \mathsf{Dir}(\alpha_i, \ldots, \alpha_k)$$

where $\alpha_i \in \mathbb{R}$ are real values that determine prior beliefs on the mixing proportions. We usually set a constant across all dimension to have flat prior (i.e., $\alpha = \alpha_i$).

A variational method is semi-parametric, as it estimates $k$ by considering all mixtures with less then an upper bound of components $k < K$. Again we can take $K = n$ for $n$ input points, or a lower value to speed up convergence. Thus, again $\alpha$ controls for the final number of clusters returned by the method. In a variational formulation, posterior estimates are approximated as

$$p(\mathbf{Z}, \boldsymbol{\pi}, \boldsymbol{\theta}) = q(\mathbf{Z})\, q(\boldsymbol{\pi}) \prod_{w=1}^{k} q(\theta_w) .$$

Here $\mathbf{Z}$ are latent variables, $\boldsymbol{\pi}$ mixing proportions and $\boldsymbol{\theta}$ parameters; the posterior distributions $q$ are assumed to factorise. This formulation is an optimisation problem in which we minimise a Kullback-Leibler divergence, and we measure convergence by monitoring the *Evidence Lower Bound* (ELBO).

***Overfitting the data.*** We expect that a direct clustering of all input read counts, i.e., without MOBSTER, inflates $k$. Here, we confirm that this happens with statistical models widely adopted in the literature, in two different ways:

1. with two implementations of core statistical methodologies for clonal deconvolution described above.
   - For Dirichlet Processes, we use the R package DPpackage which provides a high-performance Fortran MCMC sampler (function DPbetabinom) that supports both scalar values for $\alpha$, and the Gamma prior (6);
   - For Dirichlet Mixture Models fit by VI, we have implemented in MOBSTER a multivariate variational fit with Binomial distributions. In our implementation we have full control over the parameters of the model, and we monitor the ELBO to assess convergence of the fit. The mathematical model is detailed in Supplementary Note 2.
2. with popular tools for the problem, based on the above methodologies.

The first test is analyses the core of the methods, the second assesses also post fit heuristics that each tool might implement, like filtering for cluster size, strength of clustering assignments, clone merging and density smoothing. What we observe here is that the overfit happens at the core of the statistical methodology, leading to systematic errors. To assess the effect of $\alpha$ on the fit, we scanned the following values:

- pointwise estimates of $\alpha = 1, 10^{-2}, 10^{-4}, 10^{-8}$ for both methodologies;
- a prior $\alpha \sim \Gamma(0.01, 0.01)$ for the Dirichlet Processes. In this case we refer to the scale parametrisation of the Gamma prior which is adopted by the DPpackage: this corresponds to a parametrisation with rate $1/0.01 = 100$.

We analysed all the $150$ tumours, and for each tumour we have fit:

1. a direct Binomial clustering of the full set of read counts in the data;
2. a first run with MOBSTER to detect tail mutations. Then, from the posterior estimates of the latent variables – responsibilities $z_{i,k}$, in clustering jargon – we used *hard clustering assignments* as

$$\mathsf{cluster}(x_i) = \arg \max_{c=1,\ldots,k} z_{i,k} .$$

We remove from data $X$ all tail SNVs: they have $\mathsf{cluster}(x_i) = 1$, because we assumed a fixed id for the tail component. Then, we run Binomial clustering on the read counts of the reduced set of input

$$X_{\mathsf{nt}} = \{x_i \in X \mid \mathsf{cluster}(x_i) \neq 1\} .$$

Notice that if MOBSTER fits the data without a tail in step 1, then $X_{\mathsf{nt}} = X$ in step 2 and hence the performance is just that of a direct clustering of the data. This means that in our tests we are considering also the ability of MOBSTER to fit tails to the data. The fit parameters of the methods are:

- with the DPpackage, we used 10,000 MCMC steps with a burn-in of 5,000 and a thinning of 3, which gives over $3,000$ samples to estimate the posterior distributions of the parameters;
- with MOBSTER's variational method we set the upper bound on the number of clusters $K = 5$, for the Beta priors of the Binomial mixture components a standard flat prior with $a_0 = b_0 = 1$, and we monitor the ELBO and stop the fit when it does not change by $\epsilon = 10^{-10}$. To avoid local optima of this method we sample $10$ independent initial conditions for the fit, but we observed no substantial variations across the results.

Key results are in the Main Text, and confirm our intuition about the expected behaviours; the full set of results is available as Supplementary Figure S8. We plot the ratio $r = k_{\mathsf{fit}}/k$ between the estimated number of clusters $k_{\mathsf{fit}}$ and the true $k$. The error is very large ($r > 1$, overfit) without MOBSTER. For tumours without subclones the tail is large and the median error for high $\alpha$ can reach $r = 4$: we call $k = 4$ Binomial clusters (1 clonal cluster, plus three subclones), where there are no subclones. The error diminishes with lower values of $\alpha$, but persists. This demonstrates how input parameters influence the outputs. For tumours with subclones, tail and subclones overlap and the error is reduced but remains inflated. The trend observed for variations of $\alpha$ is the same across all tumours, regardless of the number of subclones.

These errors reduce almost to zero if one uses MOBSTER to "clean" the signal from tail SNVs. We still observe a trend due to the variation of $\alpha$, but it is evident that for $\alpha \leq 10^{-4}$ the true model is retrieved. With $\alpha = 10^{-4}$ we obtain the same performance of a Gamma prior on $\alpha$, in the Dirichlet Process fit. Both a prior and a pointwise estimate of $\alpha$ cannot retrieve the true model, without MOBSTER. In general, the combination with lowest variance and highest precision of the fit, is obtained by combining MOBSTER with the Dirichlet Process fit, and $\alpha = 10^{-4}$.

**Comparison against well-known tools.** In the Main Text we also compare DPclust, pyClone – with both Binomial and Beta-Binomial distributions – and sciClone on the same test set described above ($n = 150$ tumours, 120x median coverage and purity 1). DPclust and pyClone are based on Dirichlet Processes, sciClone on Dirichlet Finite Mixtures. The tests above has been used to identify optimal values for $\alpha$, a parameter which is hardcoded in the implementation of DPclust and sciClone. Parameters and comments on these simulations are reported in the Main Text.

### 2. Note 2: Variational inference for Dirichlet mixtures of multivariate Binomials

There are several tools that one can use in order to cluster read counts from tumour mutations, after MOBSTER analysis of tumour VAF. Our tool is already interfaced to some of them (sciClone and pyClone), but we also reasoned that one should be able to carry out a full analysis with just one tool; for this reason, we implemented in MOBSTER a variational inference method for Dirichlet mixtures of multivariate Binomial distributions (which should be faster than a Montecarlo sampler for a Dirichlet Process). This statistical model represents and alternative to popular tools for this task, and mimics the statistical model available in SciClone (4). Here however there are two main differences: $(i)$ we do not implement post-fit heuristic to filter output clusters or assign mutations to clusters (as SciClone does through some form of hypothesis testing), and $(ii)$ we expose all parameters for the fit and convergence (e.g., the concentration, the priors, etc.), which in tools like SciClone are hardcoded in the implementation. We first introduce the variational method for single-sample (i.e., univariate) data, and then we extend it to multi-region (i.e., multivariate) data.

**Data.** The input data $\mathbf{X}$ consists of $N$ pairs of values describing independent experiments $\mathbf{X} = [\boldsymbol{s}\ \boldsymbol{t}]$ where $\boldsymbol{s}$ and $\boldsymbol{t}$ are the $N$-dimensional vectors with the counts of successful and total trials in each experiment. In this context $\boldsymbol{s}$ represents reads with the mutated allele, while $\boldsymbol{t}$ the total number of reads at the locus (the per-locus coverage). For the $n$-th datapoint, we denote its values as scalars $s_n$ and $t_n$; this notation is used to explicit the Binomial likelihood – e.g., $\mathsf{Bin}(s_n \mid t_n, \theta_k)$.

**Bayesian model.** We consider a *Dirichlet Finite-Mixture Model with $K$ Binomial mixtures*, where each mixture's parameter – the Binomial probability $\theta$ – has a conjugate Beta prior

$$
\begin{aligned}
\boldsymbol{\pi} \mid \boldsymbol{\alpha} &\sim \mathsf{Dir}(\alpha_0, \ldots, \alpha_0) && \text{[mixture proportions]}\\
\mathbf{Z} \mid \boldsymbol{\pi} &\sim \mathsf{Cat}(\boldsymbol{\pi}) && \text{[latent variables]}\\
\theta_k \mid a, b &\sim \mathsf{Beta}(a_0, b_0) && \text{[parameter of a mixture]}\\
f_k \mid \theta_k &\sim \mathsf{Bin}(\boldsymbol{s} \mid \boldsymbol{t}, \theta_k) && \text{[likelihood of a mixture]}
\end{aligned}
$$

so the fixed-value *hyperparameters* are the scalars $\alpha_0$, $a_0$ and $b_0$.

**Joint and variational distributions.** We want to infer the following factorized distribution

$$
\begin{aligned}
p(\mathbf{X}, \mathbf{Z}, \boldsymbol{\pi}, \boldsymbol{\theta}) &= p(\mathbf{X} \mid \mathbf{Z}, \boldsymbol{\pi}, \boldsymbol{\theta})\, p(\mathbf{Z}, \boldsymbol{\pi}, \boldsymbol{\theta})\\
&= p(\mathbf{X} \mid \mathbf{Z}, \boldsymbol{\pi}, \boldsymbol{\theta})\, p(\mathbf{Z} \mid \boldsymbol{\pi}, \boldsymbol{\theta})\, p(\boldsymbol{\pi}, \boldsymbol{\theta})\\
&= p(\mathbf{X} \mid \mathbf{Z}, \boldsymbol{\pi}, \boldsymbol{\theta})\, p(\mathbf{Z} \mid \boldsymbol{\pi}, \boldsymbol{\theta})\, p(\boldsymbol{\pi})\, p(\boldsymbol{\pi}) = \underbrace{p(\mathbf{X} \mid \mathbf{Z}, \boldsymbol{\theta})}_{\text{mix. likelihood}}\, p(\mathbf{Z} \mid \boldsymbol{\pi})\, p(\boldsymbol{\pi}) \prod_k p(\theta_k)
\end{aligned}
\qquad [1]
$$

where the term that depends on the Binomial mixtures are $p(\mathbf{X} \mid \mathbf{Z}, \boldsymbol{\theta})$ and the prior $p(\theta_k)$. Notice that we write product terms over $k$ without the upper index $K$; we use the same notation for the samples, and omit $N$. As usual, the variational distribution is written as

$$q(\mathbf{Z}, \boldsymbol{\pi}, \boldsymbol{\theta}) = q(\mathbf{Z})\, q(\boldsymbol{\pi}, \boldsymbol{\theta}) \,. \tag{2}$$

**Key equations.** We will use these equations to compute expectations.

- the multinomial distribution of the latent variables $\mathbf{Z} \mid \boldsymbol{\pi} \sim \mathsf{Cat}(\mathbf{Z} \mid \boldsymbol{\pi})$ is

$$\ln p(\mathbf{Z} \mid \boldsymbol{\pi}) = \ln \mathsf{Cat}(\mathbf{Z} \mid \boldsymbol{\pi}) = \ln \prod_{n,k} \pi_k^{z_{nk}} = \sum_{n,k} z_{nk} \, \ln \pi_k \,. \tag{3}$$

- the Binomial likelihood of the data given the latent variables with iid samples is

$$\begin{aligned}
\ln p(\mathbf{X} \mid \mathbf{Z}, \boldsymbol{\theta}) &= \ln \prod_{n,k} \mathsf{Bin}(s_n \mid t_n, \theta_k)^{z_{nk}} = \sum_{n,k} z_{nk} \ln \mathsf{Bin}(s_n \mid t_n, \theta_k) \\
&= \sum_{n,k} z_{nk} \Big\{ s_n \ln \theta_k + (t_n - s_n) \ln(1-\theta_k) \Big\} + const \\
\ln \mathsf{Bin}(s_n \mid t_n, \theta_k) &= \ln \binom{t_n}{s_n} \theta_k^{s_n} (1-\theta_k)^{t_n - s_n} = s_n \ln \theta_k + (t_n - s_n) \ln(1-\theta_k) + const \,.
\end{aligned} \tag{4}$$

  Here and in what follows *const* is the normalization constant.

- the prior for the mixture components is the Dirichlet likelihood with hyperparameter $\boldsymbol{\alpha}_0 = [\alpha_0 \cdots \alpha_0]$ (notice that the components are independent)

$$\ln p(\boldsymbol{\pi}) = \ln \mathsf{Dir}(\boldsymbol{\pi} \mid \boldsymbol{\alpha}_0) = \ln \prod_k \pi_k^{\alpha_0 - 1} + const = (\alpha_0 - 1) \sum_k \ln \pi_k + const \,. \tag{5}$$

- The conjugate prior for the parameters of a mixture component $\theta_k \sim \mathsf{Beta}(a_0, b_0)$ has hyperparameters $a_0$ and $b_0$ and log-likelihood

$$\ln p(\theta_k) = \ln \mathsf{Beta}(\theta_k \mid a_0, b_0) = (a_0 - 1) \ln \theta_k + (b_0 - 1) \ln(1-\theta_k) + const \,. \tag{6}$$

#### A. Update factors $q(\mathbf{Z})$, $q(\boldsymbol{\pi})$ and $q(\theta_k)$.

**Derivation of $q(\mathbf{Z})$** This is the expectation with respect to $\boldsymbol{\pi}$ and $\boldsymbol{\theta}$ of the logarithm of the joint distribution that we want to infer

$$\begin{aligned}
\ln q(\mathbf{Z}) &= \mathbb{E}_{\boldsymbol{\pi},\boldsymbol{\theta}}[\ln p(\mathbf{X}, \mathbf{Z}, \boldsymbol{\pi}, \boldsymbol{\theta})] \\
&= \mathbb{E}_{\boldsymbol{\theta}}[\ln p(\mathbf{X} \mid \mathbf{Z}, \boldsymbol{\theta})] + \mathbb{E}_{\boldsymbol{\pi}}[\ln p(\mathbf{Z} \mid \boldsymbol{\pi})] + \mathbb{E}_{\boldsymbol{\pi}}[\ln p(\boldsymbol{\pi})] + \mathbb{E}_{\boldsymbol{\theta}}[\ln p(\boldsymbol{\theta})] && \text{[ln of (1)]} \\
&= \mathbb{E}_{\boldsymbol{\theta}}[\ln p(\mathbf{X} \mid \mathbf{Z}, \boldsymbol{\theta})] + \mathbb{E}_{\boldsymbol{\pi}}[\ln p(\mathbf{Z} \mid \boldsymbol{\pi})] + const \,,
\end{aligned}$$

where with *const* we group all terms that do not depend on $\mathbf{Z}$ (the parameter of $q$).

These are expectations of factors computed above: for the first term the expectation is over $\boldsymbol{\theta}$, and distributes over the Binomial terms, for the latter over $\pi_k$. We obtain

$$\begin{aligned}
\mathbb{E}_{\boldsymbol{\pi}}[\ln p(\mathbf{Z} \mid \boldsymbol{\pi})] &= \sum_{n,k} z_{nk} \, \mathbb{E}_{\pi_k} \left[\ln \pi_k\right] \,, && \text{[by (3)]} \\
\mathbb{E}_{\boldsymbol{\theta}}[\ln p(\mathbf{X} \mid \mathbf{Z}, \theta)] &= \sum_{n,k} z_{nk} \, \mathbb{E}_{\theta_k} \left[s_n \ln \theta_k + (t_n - s_n) \ln(1-\theta_k)\right] + const && \text{[by (4)]} \\
&= \sum_{n,k} z_{nk} \left\{ s_n \mathbb{E}_{\theta_k} \left[\ln \theta_k\right] + (t_n - s_n) \mathbb{E}_{\theta_k} \left[\ln(1-\theta_k)\right] \right\} + const \,.
\end{aligned}$$

Notice that the normalization constants are absorbed in *const* (they do not depend on $\theta_k$). We then combine everything to derive $q(\mathbf{Z})$, and group the terms by $z_{nk}$

$$\begin{aligned}
\ln q(\mathbf{Z}) &= \sum_{n,k} z_{nk} \left\{ \mathbb{E}_{\pi_k} \left[\ln \pi_k\right] + s_n \mathbb{E}_{\theta_k} \left[\ln \theta_k\right] + (t_n - s_n) \mathbb{E}_{\theta_k} \left[\ln(1-\theta_k)\right] \right\} + const \\
&= \sum_{n,k} z_{nk} \, \ln \lambda_{nk} + const
\end{aligned} \tag{7}$$

where we defined $N \times K$ terms

$$\ln \lambda_{nk} = \mathbb{E}_{\pi_k}[\ln \pi_k] + s_n \mathbb{E}_{\theta_k}[\ln \theta_k] + (t_n - s_n)\mathbb{E}_{\theta_k}[\ln(1-\theta_k)] \,. \tag{8}$$

The *responsibilities* $r_{nk}$ are computed taking logarithms out, and exponentiating

$$q(\mathbf{Z}) \propto \prod_{n,k} \lambda_{nk}^{z_{nk}}$$

so we can finally normalize this distribution

$$q(\mathbf{Z}) = \prod_{n,k} r_{nk}^{z_{nk}} \,, \qquad r_{nk} = \frac{\lambda_{nk}}{\sum_j \lambda_{jk}} \,. \tag{9}$$

**Derivation of** $q(\boldsymbol{\pi}, \boldsymbol{\theta})$ First we show that $q(\boldsymbol{\pi}, \boldsymbol{\theta})$ can be factorized, and then proceed further on its factors

$$\begin{aligned} \ln q(\boldsymbol{\pi}, \boldsymbol{\theta}) &= \mathbb{E}_{\mathbf{Z}}[\ln p(\mathbf{X}, \mathbf{Z}, \boldsymbol{\pi}, \boldsymbol{\theta})] = \\ &= \mathbb{E}_{\mathbf{Z}}[\ln p(\mathbf{X} \mid \mathbf{Z}, \boldsymbol{\theta})] + \mathbb{E}_{\mathbf{Z}}[\ln p(\mathbf{Z} \mid \boldsymbol{\pi})] + \mathbb{E}_{\mathbf{Z}}[\ln p(\boldsymbol{\pi})] + \mathbb{E}_{\mathbf{Z}}[\ln p(\boldsymbol{\theta})] \,. \end{aligned} \qquad \text{[ln of (1)]}$$

We consider all terms because they involve the parameters of $q(\boldsymbol{\pi}, \boldsymbol{\theta})$. Expectations over $\mathbf{Z}$ are trivial for terms that do not depend on latent variables, while the mixture likelihood has now an expectation over $\mathbf{Z}$

$$\begin{aligned} \mathbb{E}_{\mathbf{Z}}[\ln p(\boldsymbol{\pi})] &= \ln p(\boldsymbol{\pi}) \\ \mathbb{E}_{\mathbf{Z}}[\ln p(\boldsymbol{\theta})] &= \sum_k \ln p(\theta_k) \\ \mathbb{E}_{\mathbf{Z}}[p(\mathbf{X} \mid \mathbf{Z}, \boldsymbol{\theta})] &= \sum_{n,k} \mathbb{E}_{\mathbf{Z}}[z_{nk}] \ln \mathsf{Bin}(s_n \mid t_n, \theta_k) \,. \end{aligned} \qquad \text{[by (4)]}$$

When we put all together we obtain

$$\ln q(\boldsymbol{\pi}, \boldsymbol{\theta}) = \sum_k \underbrace{\sum_n \Big\{ \mathbb{E}_{\mathbf{Z}}[z_{nk}] \, \ln \mathsf{Bin}(s_n \mid t_n, \theta_k) + \ln p(\theta_k) \Big\}}_{\ln q(\theta_k)} + \underbrace{\mathbb{E}_{\mathbf{Z}}[\ln p(\mathbf{Z} \mid \boldsymbol{\pi})] + \ln p(\boldsymbol{\pi})}_{\ln q(\boldsymbol{\pi})}$$

and by taking logarithms out and exponentiating we obtain the factorisation

$$q(\boldsymbol{\pi}, \boldsymbol{\theta}) = q(\boldsymbol{\pi}) \prod_k q(\theta_k) \,. \tag{10}$$

**Derivation of** $q(\boldsymbol{\pi})$ This involves only the Dirichlet distribution, which is the conjugate prior of the categorical (7). The expectation for the latent variables are the responsibilities ($\mathbb{E}_{\mathbf{Z}}[z_{nk}] = r_{nk}$)

$$\begin{aligned} \ln q(\boldsymbol{\pi}) &= \mathbb{E}_{\mathbf{Z}}[\ln p(\mathbf{Z} \mid \boldsymbol{\pi})] + \ln p(\boldsymbol{\pi}) + const \\ &= \sum_{n,k} \underbrace{\mathbb{E}_{\mathbf{Z}}[z_{nk}]}_{r_{nk}} \ln \pi_k + \sum_k \ln \pi_k^{\alpha_0 - 1} + const && \text{[by (3) and (5)]} \\ &= \sum_{n,k} \ln \pi_k^{r_{nk}} + \ln \pi_k^{\alpha_0 - 1} + const && [\mathbb{E}_{\mathbf{Z}}[z_{nk}] = r_{nk}] \end{aligned}$$

Then by taking out the logarithms and exponentiating one gets

$$\begin{aligned} q(\boldsymbol{\pi}) &\propto \prod_k \pi_k^{\alpha_0 - 1} \cdot \pi_k^{\sum_n r_{nk}} = \prod_k \pi_k^{\alpha_0 + N_k - 1} && [N_k = \textstyle\sum_n r_{nk}] \\ &= \mathsf{Dir}(\boldsymbol{\pi} \mid \boldsymbol{\alpha}_0 + N_k) && (11) \end{aligned}$$

so the posterior is Dirichlet with components $\alpha_k = \alpha_0 + N_k$

***Derivation of*** $q(\theta_k)$**.** The posterior for a Binomial mixture component with Beta prior is Beta by conjugacy, as we show here.

$$\ln q(\theta_k) = \sum_n \mathbb{E}_{\mathbf{Z}}[z_{nk}] \, \ln \mathsf{Bin}(s_n \mid t_n, \theta_k) + \ln p(\theta_k) + const$$
$$= \sum_n \mathbb{E}_{\mathbf{Z}}[z_{nk}] \Big\{ s_n \ln \theta_k + (t_n - s_n) \ln(1-\theta_k) \Big\} + (a_0 - 1) \ln \theta_k + (b_0 - 1) \ln(1-\theta_k) + const$$

Considering $\mathbb{E}_{\mathbf{Z}}[z_{nk}] = r_{nk}$ as in the derivation of $q(\boldsymbol{\pi})$ we have

$$\ln q(\theta_k) = \sum_n \ln \theta_k^{s_n r_{nk}} + \ln(1-\theta_k)^{(t_n - s_n) r_{nk}} + \ln \theta_k^{a_0 - 1} + \ln(1-\theta_k)^{b_0 - 1} + const$$
$$= \ln \theta_k^{a_0 - 1} \prod_n \theta_k^{s_n r_{nk}} \cdot (1-\theta_k)^{b_0 - 1} \prod_n (1-\theta_k)^{(t_n - s_n) r_{nk}} + const$$

which has the general form $\ln \theta_k^x \cdot (1-\theta_k)^y + const$ with

$$x = \sum_n s_n r_{nk} + a_0 - 1 \qquad\qquad y = \sum_n (t_n - s_n) r_{nk} + b_0 - 1 \, .$$

Now if we define $s_k^\star = \sum_n s_n r_{nk}$ and $t_k^\star = \sum_n t_n r_{nk}$, we find

$$q(\theta_k) \propto \theta_k^{a_0 + s_k^\star - 1} \cdot (1-\theta_k)^{b_0 + t_k^\star - s_k^\star - 1} = \mathsf{Beta}(\theta_k \mid a_0 + s_k^\star, b_0 + t_k^\star - s_k^\star) \, , \qquad [12]$$

which suggests the posterior update rules $a_k = a_0 + s_k^\star$ and $b_k = a_0 + t_k^\star - s_k^\star$.

**B. Update equations (variational E and M-steps).** By the previous equations we obtain the E-variational and M-variational steps.

**Variational E-step** This step consists in computing the responsibilities $r_{nk}$, which depend on our approximation to the posterior over the parameters. To compute $r_{nk}$ we have to compute the $\lambda_{nk}$ terms, then the normalization is just empirical to get each of the $r_{nk}$ values. The formula for the $\lambda_{nk}$ terms is

$$\ln \lambda_{nk} = \mathbb{E}_{\pi_k}[\ln \pi_k] + s_n \mathbb{E}_{\theta_k}[\ln \theta_k] + (t_n - s_n) \mathbb{E}_{\theta_k}[\ln(1-\theta_k)] \, , \qquad [13]$$

which can be split in the computation of three expectations (under $q$).

Term $\mathbb{E}_{\pi_k}[\ln \pi_k]$ is known (7) to be $\mathbb{E}_{\pi_k}[\ln \pi_k] = \Psi(\alpha_k) - \Psi(\boldsymbol{\alpha}_*)$ with $\boldsymbol{\alpha}_* = \sum_k \alpha_k$, where $\Psi$ is the *digamma* function. For term $s_n \mathbb{E}_{\theta_k}[\ln \theta_k]$it is possible to see that[†]

$$\mathbb{E}_{\theta_k}[\ln \theta_k] = \int_0^1 \ln \theta_k \cdot q(\theta_k) \, d\theta_k = \frac{1}{B(a_k, b_k)} \int_0^1 \ln \theta_k \cdot \theta_k^{a_k - 1} (1-\theta_k)^{b_k - 1} \, d\theta_k \qquad \text{[by def.]}$$
$$= \frac{1}{B(a_k, b_k)} \frac{\Gamma(a_k)\Gamma(b_k)}{\Gamma(a_k + b_k)} \Big[ \Psi(a_k) - \Psi(a_k + b_k) \Big]$$
$$= \Psi(a_k) - \Psi(a_k + b_k) \qquad \text{[since } B(a_k, b_k) = \Gamma(a_k)\Gamma(b_k)\Gamma(a_k+b_k)^{-1}\text{]}$$

which means that $s_n \mathbb{E}_{\theta_k}[\ln \theta_k] = s_n \Big[ \Psi(a_k) - \Psi(a_k + b_k) \Big]$.

**Term** $(t_n - s_n)\mathbb{E}_{\theta_k}[\ln 1 - \theta_k]$. This term is similar to above; the expectation is again over the same Beta distribution. To solve it, we can apply a direct substitution for $1 - \theta_k$

$$\mathbb{E}_{\theta_k}[\ln 1 - \theta_k] = \int_0^1 \ln(1-\theta_k) \cdot q(\theta_k) \, d\theta_k \qquad \text{[by def.]}$$
$$= \frac{1}{B(a_k, b_k)} \int_0^1 \ln(1-\theta_k) \cdot \theta_k^{a_k - 1} (1-\theta_k)^{b_k - 1} \, d\theta_k$$
$$= \frac{-1}{B(b_k, a_k)} \int_{0=y}^{1=y} \ln y \cdot y^{b_k - 1} (1-y)^{a_k - 1} \, dy \qquad [y = 1 - \theta_k, \, d\theta_k = -dy]$$
$$= \Psi(b_k) - \Psi(a_k + b_k) \qquad \text{[as above, with reversed sign]}$$

where we used $B(b_k, a_k) = B(a_k, b_k)$ and obtained a Beta with swapped parameters, overall

$$(t_n - s_n)\mathbb{E}_{\theta_k}[\ln 1 - \theta_k] = (t_n - s_n) \Big[ \Psi(b_k) - \Psi(a_k + b_k) \Big] \, .$$

[†]There is an explicit derivation at https://stats.stackexchange.com/questions/241993/pdf-of-y-logx-when-x-is-beta-distributed-the-expected-value-of-y/242020

**Computing the $\lambda_{nk}$ terms.** The formula for $\lambda_{nk}$ is just

$$\begin{aligned}\lambda_{nk} &\propto \exp\left\{\Psi(\alpha_k) - \Psi(\boldsymbol{\alpha}_*) + s_n\Big[\Psi(a_k) - \Psi(a_k+b_k)\Big] + (t_n - s_n)\Big[\Psi(b_k) - \Psi(a_k+b_k)\Big]\right\}\\ &\propto \exp\left\{\Psi(\alpha_k) - \Psi(\boldsymbol{\alpha}_*) + s_n\Big[\Psi(a_k) - \Psi(b_k)\Big] + t_n\Big[\Psi(b_k) - \Psi(a_k+b_k)\Big]\right\},\end{aligned} \quad [14]$$

and values for $r_{nk}$ can be computed by normalizing these terms.

**Variational M-step** The other quantities of interest are the rules to compute the posterior approximations to the parameters of the Dirichlet and Beta distributions, and they have been derived earlier.

$$\alpha_k = \alpha_0 + N_k \qquad a_k = a_0 + s_k^\star \qquad b_k = b_0 + t_k^\star - s_k^\star \quad [15]$$

where

$$N_k = \sum_n r_{nk} \qquad s_k^\star = \sum_n s_n r_{nk} \qquad t_k^\star = \sum_n t_n r_{nk}\,.$$

**C. Variational evidence lower bound $\mathcal{L}(q)$.** The *evidence lower bound* (ELBO) $\mathcal{L}(q)$ is computed at each iteration of our variational inference to assess convergency. For our mixture of Binomials the ELBO is

$$\begin{aligned}\mathcal{L}(q) &= \sum_{\mathbf{Z}} \int\int q(\mathbf{Z},\boldsymbol{\pi},\boldsymbol{\theta}) \ln\left\{\frac{p(\mathbf{X},\mathbf{Z},\boldsymbol{\pi},\boldsymbol{\theta})}{q(\mathbf{Z},\boldsymbol{\pi},\boldsymbol{\theta})}\right\} d\boldsymbol{\pi} d\boldsymbol{\theta} =\\ &= \mathbb{E}_{\mathbf{Z},\boldsymbol{\pi},\boldsymbol{\theta}}\left[\ln p(\mathbf{X},\mathbf{Z},\boldsymbol{\pi},\boldsymbol{\theta})\right] - \mathbb{E}_{\mathbf{Z},\boldsymbol{\pi},\boldsymbol{\theta}}\left[\ln q(\mathbf{Z},\boldsymbol{\pi},\boldsymbol{\theta})\right]\end{aligned}$$

Since $p$ and $q$ factorize we can rewrite the bound as

$$\begin{aligned}\mathcal{L}(q) &= \mathbb{E}_{\mathbf{Z},\boldsymbol{\pi},\boldsymbol{\theta}}\left[\ln p(\mathbf{X}\mid\mathbf{Z},\boldsymbol{\theta})\,p(\mathbf{Z}\mid\boldsymbol{\pi})\,p(\boldsymbol{\pi})\prod_k p(\theta_k)\right] - \mathbb{E}_{\mathbf{Z},\boldsymbol{\pi},\boldsymbol{\theta}}\left[\ln q(\mathbf{Z})q(\boldsymbol{\pi})\prod_k q(\theta_k)\right]\\ &= \mathbb{E}_{\mathbf{Z},\boldsymbol{\theta}}\left[\ln p(\mathbf{X}\mid\mathbf{Z},\boldsymbol{\theta})\right] + \mathbb{E}_{\mathbf{Z},\boldsymbol{\pi}}\left[\ln p(\mathbf{Z}\mid\boldsymbol{\pi})\right] + \mathbb{E}_{\boldsymbol{\pi}}\left[\ln p(\boldsymbol{\pi})\right] + \sum_k \mathbb{E}_{\theta_k}\left[\ln p(\theta_k)\right]\\ &\quad - \mathbb{E}_{\mathbf{Z}}\left[\ln q(\mathbf{Z})\right] - \mathbb{E}_{\boldsymbol{\pi}}\left[\ln q(\boldsymbol{\pi})\right] - \sum_k \mathbb{E}_{\theta_k}\left[\ln q(\theta_k)\right]\,.\end{aligned}$$

We rewrite this form of the ELBO by using the KL-divergence among the involved distributions (Section E)

$$\mathcal{L}(q) = \mathbb{E}_{\mathbf{Z},\boldsymbol{\theta}}\left[\ln p(\mathbf{X}\mid\mathbf{Z},\boldsymbol{\theta})\right] + w[q(\mathbf{Z}), p(\mathbf{Z}\mid\boldsymbol{\pi})] + w[q(\boldsymbol{\pi}) \parallel p(\boldsymbol{\pi})] + \sum_{k=1}^{K} w[q(\theta_k) \parallel p(\theta_k)]$$

where $w[q,p] = -\mathsf{KL}[q,p]$. The only terms that we need to compute are the first, and the last, as the other are independent on the mixture and derived in Section E.

**Term $\mathbb{E}_{\mathbf{Z},\boldsymbol{\theta}}\left[\ln p(\mathbf{X}\mid\mathbf{Z},\boldsymbol{\theta})\right]$.** Once expanded, this is an expectation of independent random variables ($z_{nk}$, $\theta_k$ and $1-\theta_k$), thus is the product of their expectations; here we include also the normalization constant $\ln C_n = \ln t_n! - \ln s_n! - \ln(t_n - s_n)!$ for a Binomial random variable

$$\begin{aligned}\mathbb{E}_{\mathbf{Z},\boldsymbol{\theta}}\left[\ln p(\mathbf{X}\mid\mathbf{Z},\boldsymbol{\theta})\right] &= \mathbb{E}_{\mathbf{Z},\boldsymbol{\theta}}\left[\sum_{n,k}\Big\{s_n \ln\theta_k + (t_n - s_n)\ln(1-\theta_k) - \ln C_n\Big\}\right]\\ &= \sum_{n,k} \mathbb{E}_{\mathbf{Z}}\left[z_{nk}\right] \mathbb{E}_{\theta_k}\left[s_n \ln\theta_k + (t_n - s_n)\ln(1-\theta_k) - \ln C_n\right]\,.\end{aligned}$$

These terms have been computed for the variational E-step, we have

$$\mathbb{E}_{\mathbf{Z},\boldsymbol{\theta}}\left[\ln p(\mathbf{X}\mid\mathbf{Z},\boldsymbol{\theta})\right] = \sum_{n,k} r_{nk}\Big\{s_n\Big[\Psi(a_k) - \Psi(b_k)\Big] + t_n\Big[\Psi(b_k) - \Psi(a_k+b_k)\Big] - \ln C_n\Big\}\,.$$

**Term $\sum_k w[q(\theta_k) \parallel p(\theta_k)]$.** A term of this summation is the negative KL-divergence among two Beta random variables, one given by the prior $p$, the other by the posterior $q$ (7)

$$\begin{aligned}w[q(\theta_k), p(\theta_k)] &= -\mathsf{KL}[q(\theta_k) \parallel p(\theta_k)]\\ &= -\left\{\ln\frac{B(a_0,b_0)}{B(a_k,b_k)} + (a_k - a_0)\Psi(a_k) + (b_k - b_0)\Psi(b_k) + (a_0 - a_k + b_0 - b_k)\Psi(a_k+b_k)\right\}\\ &= \ln\frac{B(a_k,b_k)}{B(a_0,b_0)} + (a_0 - a_k)\Psi(a_k) + (b_0 - b_k)\Psi(b_k) + (a_k - a_0 + b_k - b_0)\Psi(a_k+b_k)\,.\end{aligned}$$

**The ELBO $\mathcal{L}(q)$.** Define the following quantities

$$\hat{\Psi}_{a_k,b_k} = \Psi(a_k) - \Psi(b_k) \qquad \hat{\Psi}_{b_k,a_k} = \Psi(b_k) - \Psi(a_k + b_k)$$
$$\ln \rho_k = \Psi(\alpha_k) - \Psi(\boldsymbol{\alpha_*}) \qquad w[q(\theta_k) \parallel p(\theta_k)] = \omega_k$$

to see that the ELBO has analytical form

$$\mathcal{L}(q) = \sum_{n,k} r_{nk} \left\{ s_n \hat{\Psi}_{a_k,b_k} + t_n \hat{\Psi}_{b_k,a_k} - \ln C_n + \ln \rho_k - \ln r_{nk} \right\}$$
$$+ \ln \frac{C(\boldsymbol{\alpha}_0)}{C(\boldsymbol{\alpha})} + \sum_k \left[ (\alpha_0 - \alpha_k) \ln \rho_k + \omega_k \right]. \qquad [16]$$

**D. Multivariate extension.** Each data point is now a vector with dimension $W$, we denote with an extra superscript the input data when we refer to specific component of the data. As in other papers, we assume that the $W$ dimensions are independent; the correlation structure is given by the latent variables which map each point (a vector) to one of $K$ clusters.

**Statistical model.** With respect to the univariate case we now have a vector of parameters $\boldsymbol{\theta}_k$ for each mixture component, and we use multivariate distributions (written in bold)

$$\boldsymbol{\theta}_k \mid \boldsymbol{a}, \boldsymbol{b} \sim \textbf{Beta}(\boldsymbol{a}_0, \boldsymbol{b}_0) \qquad \text{[parameter of a mixture]}$$
$$f_k \mid \boldsymbol{\theta}_k \sim \textbf{Bin}(\boldsymbol{s} \mid \boldsymbol{t}, \boldsymbol{\theta}_k) \qquad \text{[likelihood of a mixture]}$$

and our hyperparameters for the Beta are now vectors as well. The joint and the variational distributions are unchanged, provided that we assume a further factorization of the likelihood function and the prior.

***Key equations.*** The multinomial distribution (latent variables), and the prior for the components are as in the univariate case.

**Binomial likelihood of the data.** The log-likelihood of a multivariate Binomial random variable with independent dimensions is

$$\ln \textbf{Bin}(\boldsymbol{s}_n \mid \boldsymbol{t}_n, \boldsymbol{\theta}_k) = \ln \prod_w \text{Bin}(s_{n,w} \mid t_{n,w}, \theta_{k,w}) = \sum_w \left[ s_{n,w} \ln \theta_{k,w} + (t_{n,w} - s_{n,w}) \ln(1 - \theta_{k,w}) \right] + const\,.$$

We omit to write that $w$ ranges from one to $W$ to reduce the notation. The Binomial complete likelihood reads as

$$\ln p(\mathbf{X} \mid \mathbf{Z}, \boldsymbol{\theta}) = \ln \prod_{n,k} \textbf{Bin}(\boldsymbol{s}_n \mid \boldsymbol{t}_n, \boldsymbol{\theta}_k)^{z_{nk}} = \sum_{n,k} z_{nk} \ln \textbf{Bin}(\boldsymbol{s}_n \mid \boldsymbol{t}_n, \boldsymbol{\theta}_k)\,. \qquad [17]$$

**Prior for the mixture parameters.** The prior is a $W$-dimensional multivariate Beta random variable $\boldsymbol{\theta}_k \sim \textbf{Beta}(\boldsymbol{a}_0, \boldsymbol{b}_0)$ with vector hyperparameters and independent dimensions

$$\ln p(\boldsymbol{\theta}_k) = \ln \textbf{Beta}(\boldsymbol{\theta}_k \mid \boldsymbol{a}_0, \boldsymbol{b}_0) = \ln \prod_w \text{Beta}(\theta_{k,w} \mid a_{0,w}, b_{0,w})$$
$$= \sum_w \left[ (a_{0,w} - 1) \ln \theta_{k,w} + (b_{0,w} - 1) \ln(1 - \theta_{k,w}) \right] + const\,. \qquad [18]$$

***Update factors for variational distributions, and update steps.*** We need to derive $q(\mathbf{Z})$ and $q(\boldsymbol{\pi}, \boldsymbol{\theta}) = q(\boldsymbol{\pi}) \prod_k q(\boldsymbol{\theta}_k)$. Note that term $q(\boldsymbol{\pi}) \propto \text{Dir}(\boldsymbol{\pi} \mid \boldsymbol{\alpha}_0 + N_k)$ is still a Dirichlet posterior as in the univariate case.

**Term $q(\mathbf{Z})$.** For term $\ln q(\mathbf{Z})$ the only change is in the expectation of the complete likelihood

$$\mathbb{E}_{\boldsymbol{\theta}}[\ln p(\mathbf{X} \mid \mathbf{Z}, \theta)] = \mathbb{E}_{\boldsymbol{\theta}} \left[ \sum_{n,k} z_{nk} \ln \textbf{Bin}(\boldsymbol{s}_n \mid \boldsymbol{t}_n, \boldsymbol{\theta}_k) \right] =$$
$$= \sum_{n,k} z_{nk} \left\{ \sum_w \left[ s_{n,w} \ln \mathbb{E}_{\theta_{k,w}}[\theta_{k,w}] + (t_{n,w} - s_{n,w}) \mathbb{E}_{\theta_{k,w}}[\ln(1 - \theta_{k,w})] \right] \right\} + const$$

We derive as usual $\ln q(\mathbf{Z}) = \sum_{n,k} z_{nk} \ln \lambda_{nk} + const$, but now we define $N \times K$ terms

$$\ln \lambda_{nk} = \mathbb{E}_{\pi_k}[\ln \pi_k] + \sum_w \left[ s_{n,w} \ln \mathbb{E}_{\theta_{k,w}}[\theta_{k,w}] + (t_{n,w} - s_{n,w}) \mathbb{E}_{\theta_{k,w}}[\ln(1 - \theta_{k,w})] \right]. \qquad [19]$$

The responsibilities are defined as usual as $r_{nk} \propto \lambda_{nk}$.

**Term $q(\boldsymbol{\theta}_k)$.** We derive the posterior parameter for a multivariate Binomial mixture component similarly to the univariate case, but with a vector of parameters per component

$$\begin{aligned}\ln q(\boldsymbol{\theta}_k) &= \sum_n \mathbb{E}_{\mathbf{Z}}[z_{nk}] \ln \textbf{Bin}(\boldsymbol{s}_n \mid \boldsymbol{t}_n, \boldsymbol{\theta}_k) + \ln p(\boldsymbol{\theta}_k) + const \\ &= \sum_n \mathbb{E}_{\mathbf{Z}}[z_{nk}] \left\{ \sum_w \ln \text{Bin}(s_{n,w} \mid t_{n,w}, \theta_{k,w}) \right\} + \sum_w \ln \text{Beta}(\theta_{k,w} \mid a_{0,w}, b_{0,w}) + const\end{aligned}$$

Considering $\mathbb{E}_{\mathbf{Z}}[z_{nk}] = r_{nk}$ we have

$$\ln q(\boldsymbol{\theta}_k) = \ln \prod_w \theta_{k,w}^{x} \cdot \prod_w (1-\theta_{k,w})^{y} + const\,.$$

with the following substitutions

$$x = \sum_n s_{n,w} r_{nk} + (a_{0,w} - 1) \qquad\qquad y = \sum_n (t_{n,w} - s_{n,w}) r_{nk} + (b_{0,w} - 1)$$

Now if one defines

$$s^{\star}_{k,w} = \sum_n s_{n,w} r_{nk} \qquad\qquad t^{\star}_{k,w} = \sum_n t_{n,w} r_{nk}$$

then

$$\begin{aligned}q(\boldsymbol{\theta}_k) &\propto \prod_w \theta_{k,w}^{a_{0,w}+s^{\star}_{k,w}-1} \cdot (1-\theta_{k,w})^{b_{0,w}+t^{\star}_{k,w}-s^{\star}_{k,w}-1} \\ &= \prod_w \text{Beta}(\theta_{k,w} \mid a_{0,w} + s^{\star}_{k,w}, b_{0,w} + t^{\star}_{k,w} - s^{\star}_{k,w}) = \textbf{Beta}(\boldsymbol{\theta}_k \mid \boldsymbol{a}_0 + S, \boldsymbol{b}_0 + T - S)\,, \end{aligned} \qquad [20]$$

which suggests the posterior update rules (per component)

$$a_{k,w} = a_{0,w} + s^{\star}_{k,w} \qquad [21]$$

$$b_{k,w} = a_{0,w} + t^{\star}_{k,w} - s^{\star}_{k,w}\,. \qquad [22]$$

***Update equations.*** By the previous equations we obtain the E-variational and M-variational steps. The formula for the $\lambda_{nk}$ terms is

$$\ln \lambda_{nk} = \mathbb{E}_{\pi_k}[\ln \pi_k] + \sum_w \left[ s_{n,w} \ln \mathbb{E}_{\theta_{k,w}}[\theta_{k,w}] + (t_{n,w} - s_{n,w}) \mathbb{E}_{\theta_{k,w}}[\ln(1-\theta_{k,w})] \right]$$

which is split again as three expectations under $q$. The first term is analogous to the univariate case, the other terms are just adjustment to the multivariate case of the original terms (since they are expectations per-component)

$$\begin{aligned} s_{n,w}\mathbb{E}_{\theta_{k,w}}[\ln \theta_{k,w}] &= s_{n,w}\left[\Psi(a_{k,w}) - \Psi(a_{k,w} + b_{k,w})\right] \\ (t_{n,w} - s_{n,w})\mathbb{E}_{\theta_{k,w}}[\ln 1 - \theta_{k,w}] &= (t_{n,w} - s_{n,w})\left[\Psi(b_{k,w}) - \Psi(a_{k,w} + b_{k,w})\right]. \end{aligned}$$

The formula for $\lambda_{nk}$ (and normalized $r_{nk}$ values), are obtained as

$$\lambda_{nk} \propto \exp\left\{ \Psi(\alpha_k) - \Psi(\boldsymbol{\alpha}_*) + \sum_w s_{n,w}\left[\Psi(a_{k,w}) - \Psi(b_{k,w})\right] + t_{n,w}\left[\Psi(b_{k,w}) - \Psi(a_{k,w} + b_{k,w})\right] \right\}.$$

The variational M-step uses the posterior approximations to the parameters derived earlier.

***Variational evidence lower bound $\mathcal{L}(q)$.*** Consider the ELBO

$$\mathcal{L}(q) = \mathbb{E}_{\mathbf{Z},\boldsymbol{\theta}}[\ln p(\mathbf{X} \mid \mathbf{Z}, \boldsymbol{\theta})] + \omega[q(\mathbf{Z}), p(\mathbf{Z} \mid \boldsymbol{\pi})] + \omega[q(\boldsymbol{\pi}) \parallel p(\boldsymbol{\pi})] + \sum_k \omega[q(\boldsymbol{\theta}_k) \parallel p(\boldsymbol{\theta}_k)]$$

where $\omega[q,p] = -\text{KL}[q,p]$. The only terms that we need to compute are the first, and the last, as the other are independent on the mixture and derived in Section E.

**Term** $\mathbb{E}_{\mathbf{Z},\boldsymbol{\theta}}\left[\ln p(\mathbf{X} \mid \mathbf{Z},\boldsymbol{\theta})\right]$. For a multivariate Binomial random variable

$$
\begin{aligned}
\mathbb{E}_{\mathbf{Z},\boldsymbol{\theta}}\left[\ln p(\mathbf{X} \mid \mathbf{Z},\boldsymbol{\theta})\right] &= \mathbb{E}_{\mathbf{Z},\boldsymbol{\theta}}\left[\sum_{n,k} z_{nk}\Big\{\sum_{w} \ln \mathsf{Bin}(s_{n,w} \mid t_{n,w},\theta_{k,w})\Big\}\right] \\
&= \sum_{n,k} \mathbb{E}_{\mathbf{Z}}\left[z_{nk}\right] \mathbb{E}_{\boldsymbol{\theta}_k}\left[\sum_{w} \ln \mathsf{Bin}(s_{n,w} \mid t_{n,w},\theta_{k,w})\right] \\
&= \sum_{n,k} \mathbb{E}_{\mathbf{Z}}\left[z_{nk}\right] \sum_{w} \mathbb{E}_{\theta_{k,w}}\left[\ln \mathsf{Bin}(s_{n,w} \mid t_{n,w},\theta_{k,w})\right] \\
&= \sum_{n,k} r_{nk} \sum_{w} \mathbb{E}_{\theta_{k,w}}\left[\ln \mathsf{Bin}(s_{n,w} \mid t_{n,w},\theta_{k,w})\right] .
\end{aligned}
$$

Now the inner expectation rewrites as

$$
\mathbb{E}_{\theta_{k,w}}\left[\ln \mathsf{Bin}(s_{n,w} \mid t_{n,w},\theta_{k,w})\right] = \mathbb{E}_{\theta_{k,w}}\left[s_{n,w} \ln \theta_{k,w} + (t_{n,w} - s_{n,w}) \ln(1-\theta_{k,w})\right] .
$$

which are the terms derived for $\ln \lambda_{nk}$ in the univariate case. Here the normalisation constant depends on the dimension $\ln C_{n,w} = \ln t_{n,w}! - \ln s_{n,w}! - \ln(t_{n,w} - s_{n,w})!$ and we have

$$
\mathbb{E}_{\mathbf{Z},\boldsymbol{\theta}}\left[\ln p(\mathbf{X} \mid \mathbf{Z},\boldsymbol{\theta})\right] = \sum_{n,k} r_{nk}\Big\{\sum_{w} s_{n,w}\Big[\Psi(a_{k,w}) - \Psi(b_{k,w})\Big] + t_{n,w}\Big[\Psi(b_{k,w}) - \Psi(a_{k,w}+b_{k,w})\Big] - \ln C_{n,w}\Big\} .
$$

**Term** $\sum_k \omega[q(\boldsymbol{\theta}_k) \parallel p(\boldsymbol{\theta}_k)]$. If we rewrite

$$
\sum_{k} \omega[q(\boldsymbol{\theta}_k) \parallel p(\boldsymbol{\theta}_k)] = \sum_{k}\left\{\sum_{w} \omega[q(\theta_{k,w}) \parallel p(\theta_{k,w})]\right\}
$$

we retrieve a negative KL-divergence among Beta random variables (as in the univariate case),

$$
\omega[q(\boldsymbol{\theta}_k) \parallel p(\boldsymbol{\theta}_k)] = -\sum_{w} \mathsf{KL}[q(\theta_{k,w}) \parallel p(\theta_{k,w})] = \sum_{w}\left[\ln \frac{B(a_{k,w},b_{k,w})}{B(a_{0,w},b_{0,w})} + \phi_w\right] .
$$

where

$$
\phi_w = (a_{0,w} - a_{k,w})\Psi(a_{k,w}) + (b_{0,w} - b_{k,w})\Psi(b_{k,w}) + (a_{k,w} - a_{0,w} + b_{k,w} - b_{0,w})\Psi(a_{k,w}+b_{k,w})
$$

**The ELBO** $\mathcal{L}(q)$. Define the following quantities

$$
\begin{aligned}
\hat{\Psi}^{w}_{a_k,b_k} &= \Psi(a_{k,w}) - \Psi(b_{k,w}) & \hat{\Psi}^{w}_{b_k,a_k} &= \Psi(b_{k,w}) - \Psi(a_{k,w}+b_{k,w}) \\
\ln \rho_k &= \Psi(\alpha_k) - \Psi(\boldsymbol{\alpha}_*) & \omega[q(\boldsymbol{\theta}_k) \parallel p(\boldsymbol{\theta}_k)] &= \eta_k
\end{aligned}
$$

to see that the ELBO has analytical form

$$
\begin{aligned}
\mathcal{L}(q) = &\sum_{n,k} r_{nk}\Big\{\sum_{w} s_{n,w}\hat{\Psi}^{w}_{a_k,b_k} + t_{n,w}\hat{\Psi}^{w}_{b_k,a_k} - \ln C_{n,w} + \ln \rho_k - \ln r_{nk}\Big\} \\
&+ \ln \frac{C(\boldsymbol{\alpha}_0)}{C(\boldsymbol{\alpha})} + \sum_{k}\Big[(\alpha_0 - \alpha_k)\ln \rho_k + \eta_k\Big] .
\end{aligned}
$$

### E. Deriving an ELBO.

**Relation to the Kullback–Leibler divergence.** When we derive the ELBO among $p$ and $q$ (distributions) we have

$$
w(X,Y) = \mathbb{E}_{X\sim q}\left[\ln Y \sim p\right] - \mathbb{E}_{X\sim q}\left[\ln X \sim q\right]
$$

where $X$ is the random variable associated to the posterior approximation, and $Y$ to the prior. Here we show that it is possible to derive a compact form for $w(X,Y)$ without computing explicitly the two expectations. First, both expectations are under $q$, while the expectation term comes from either $p$ or $q$. Thus, the general ELBO term is a negative cross-entropy

$$
\mathbb{E}_{X\sim q}\left[\ln Y \sim f_Y\right] = \int q(x) \ln f_Y(x)\, dx = -\Big\{\mathcal{H}[X \sim q] + \mathsf{KL}(q \parallel f_Y)\Big\} = -\underbrace{\mathcal{H}[X \sim q, Y \sim f_Y]}_{\text{cross-entropy}},
$$

with $f_Y = \{p, q\}$. This is an actual cross-entropy when $X \neq Y$ ($f_Y = p$), and reduces to a negative entropy when $X = Y$ ($f_Y = q$) because the KL-divergence from self is 0

$$\mathbb{E}_{X\sim q}[\ln X \sim q] = \int q(x) \ln q(x)\, dx = -\mathcal{H}[X \sim q] \, .$$

This means that we can find a form for the ELBO which does not require the expectations with respect to $q$ and $p$. In practice

$$\begin{aligned} w(X,Y) &= \mathbb{E}_{X\sim q}[\ln Y \sim f_Y] - \mathbb{E}_{X\sim q}[\ln X \sim q] \\ &= -\mathcal{H}[X \sim q, Y \sim p] - (-\mathcal{H}[X \sim q]) \\ &= \mathcal{H}[X \sim q] - \mathcal{H}[X \sim q, Y \sim p] \\ &= \mathcal{H}[X \sim q] - (\mathcal{H}[X \sim q] + \mathsf{KL}(q \parallel p)) = -\mathsf{KL}(q \parallel p) \end{aligned}$$

This alternative formulation becomes handy when the KL-divergence among $q$ and $p$ is known.

**Terms independent of the mixture likelihoods (see (7)).**

**Proportions of the mixture $\pi$.** The KL-divergence for Dirichlet distributions with parameters $\boldsymbol{\alpha}$ and $\boldsymbol{\beta}$ is

$$\begin{aligned} \mathsf{KL}[q \sim \mathsf{Dir}(\boldsymbol{\alpha}) \parallel p \sim \mathsf{Dir}(\boldsymbol{\beta})] &= \underbrace{\ln \frac{\Gamma(\boldsymbol{\alpha}_*)}{\prod_k \Gamma(\alpha_k)}}_{C(\boldsymbol{\alpha})} + \underbrace{\ln \frac{\prod_k \Gamma(\beta_k)}{\Gamma(\boldsymbol{\beta}_*)}}_{1/C(\boldsymbol{\beta})} + \sum_k (\alpha_k - \beta_k) \underbrace{\Big[\Psi(\alpha_k) - \Psi(\boldsymbol{\alpha}_*)\Big]}_{\ln \rho_k} \\ &= \ln \frac{C(\boldsymbol{\alpha})}{C(\boldsymbol{\beta})} + \sum_k (\alpha_k - \beta_k) \ln \rho_k \end{aligned}$$

where $\boldsymbol{\alpha}_* = \sum_k \alpha_k$. In our case we have a scalar hyperparameter $\alpha_0$, which leads to $p \sim \mathsf{Dir}(\boldsymbol{\alpha}_0 = [\cdots \alpha_0 \cdots])$

$$\begin{aligned} w(q(\boldsymbol{\pi}), p(\boldsymbol{\pi})) &= -\mathsf{KL}[q \sim \mathsf{Dir}(\boldsymbol{\alpha}) \parallel p \sim \mathsf{Dir}(\boldsymbol{\alpha}_0)] \\ &= -\left\{ \ln \frac{C(\boldsymbol{\alpha})}{C(\boldsymbol{\alpha}_0)} + \sum_k (\alpha_k - \alpha_0) \Big[\Psi(\alpha_k) - \Psi(\boldsymbol{\alpha}_*)\Big] \right\} \\ &= \ln \frac{C(\boldsymbol{\alpha}_0)}{C(\boldsymbol{\alpha})} + \sum_k (\alpha_0 - \alpha_k) \ln \rho_k \, . \end{aligned}$$

To avoid numerical issues in the implementation it is convenient to use the transform

$$\ln \frac{C(\boldsymbol{\alpha}_0)}{C(\boldsymbol{\alpha})} = \ln C(\boldsymbol{\alpha}_0) - \ln C(\boldsymbol{\alpha}) = \ln \Gamma(K\alpha_0) - K \ln \Gamma(\alpha_0) - \ln \Gamma\left(\sum_k \alpha_i\right) + \sum_k \ln \Gamma(\alpha_i) \, .$$

**Latent variables.** For this term we have

$$\begin{aligned} w(q(\mathbf{Z}), p(\mathbf{Z} \mid \boldsymbol{\pi})) &= \mathbb{E}_{\mathbf{Z},\boldsymbol{\pi}}[\ln p(\mathbf{Z} \mid \boldsymbol{\pi})] - \mathbb{E}_{\mathbf{Z}}[\ln q(\mathbf{Z})] \\ &= \sum_{n,k} r_{nk} \ln \rho_k - \sum_{n,k} r_{nk} \ln r_{nk} \\ &= \sum_{n,k} r_{nk} (\ln \rho_k - \ln r_{nk}) \, . \end{aligned}$$

with as above $\ln \rho_k = \Psi(\alpha_k) - \Psi(\boldsymbol{\alpha}_*)$. It is easy to verify why this is correct: if one changes sign this becomes an entropy minus a cross-entropy, which is a KL-divergence.

### 3. Note 3: Software and availability

*The implementation of* MOBSTER *is preliminary available to the reviewers, as an attachment to the submission.*

MOBSTER is an R package freely available at

https://github.com/caravagn/MOBSTER

The implementation is parallelised, and provides S3 objects with methods to compute summary statistics from the fit, and summary plots from both the data and the fit. The Variational Dirichlet Mixture Models with Binomial data described in Note 2 is also implemented in the tool.
Other packages used are:

- DPpackage version 1.1-7.4 (available at CRAN).
- pyClone version 0.13 (available at Conda).
- DPclust version 2.2.5 (available at GitHub).
- SciClone version 1.10 (available at GitHub).

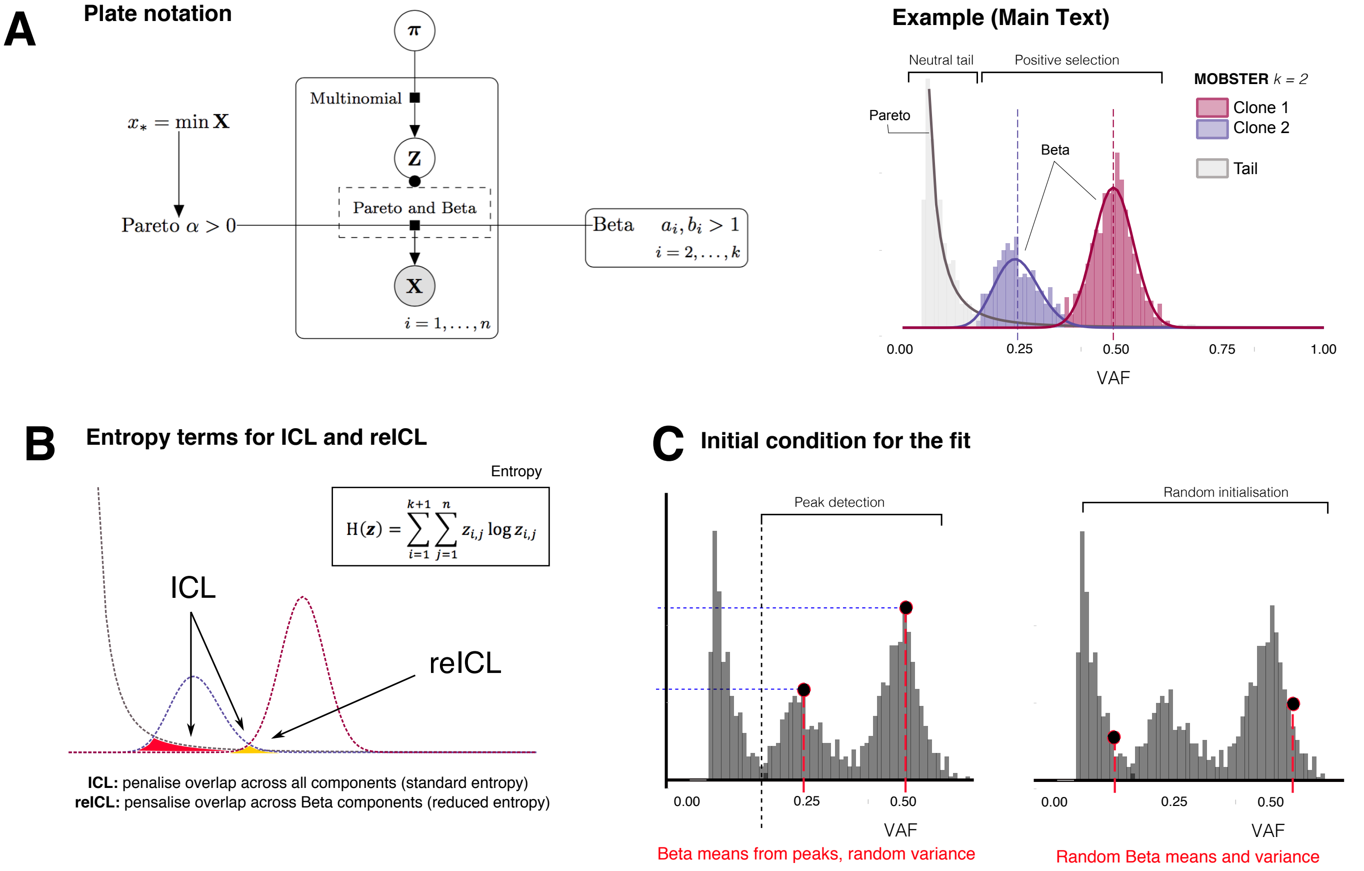

**Supplementary Figure S1. A.** MOBSTER combines $k$ Beta for the clones, and one Pareto for a power-law tail. For the tail, we can fix the scale $x_*$ from data, and learn $\alpha > 0$. For $k$ Beta components we learn the full distributions. We compute point estimates of the parameters via MLE (numerical EM), or with an iterative method of moments (analytical). In the right plot we show the example model discussed in the Main Text when introducing the method. **B.** In clustering, we use entropy to reduce the overlap between the components of the mixture. In this case, the ICL uses a standard entropy which penalises the overlap all mixture components (Beta and tail). This can lead to excessive penalisations when there is a true subclone in the data, which by definition overlaps with the tail (red and yellow areas). This is a peculiar characteristic of this clustering problem, where we do not have total separation between the components of the mixture. reICL is a variation to the ICL which uses a reduced entropy (only yellow area). This is computed by first removing from the set of latent variables the ones that are assigned to the tail (hard clustering assignments), renormalising the latent variable posterior estimates to account only for the Beta components, and using there a standard entropy. **C.** Initialisation of the fits is computed either via peak detection, or randomly. Peaks are detected from a minimum VAF value, looking at the empirical density of the data, while variance is randomly sampled. Random initialisations are totally stochastic.

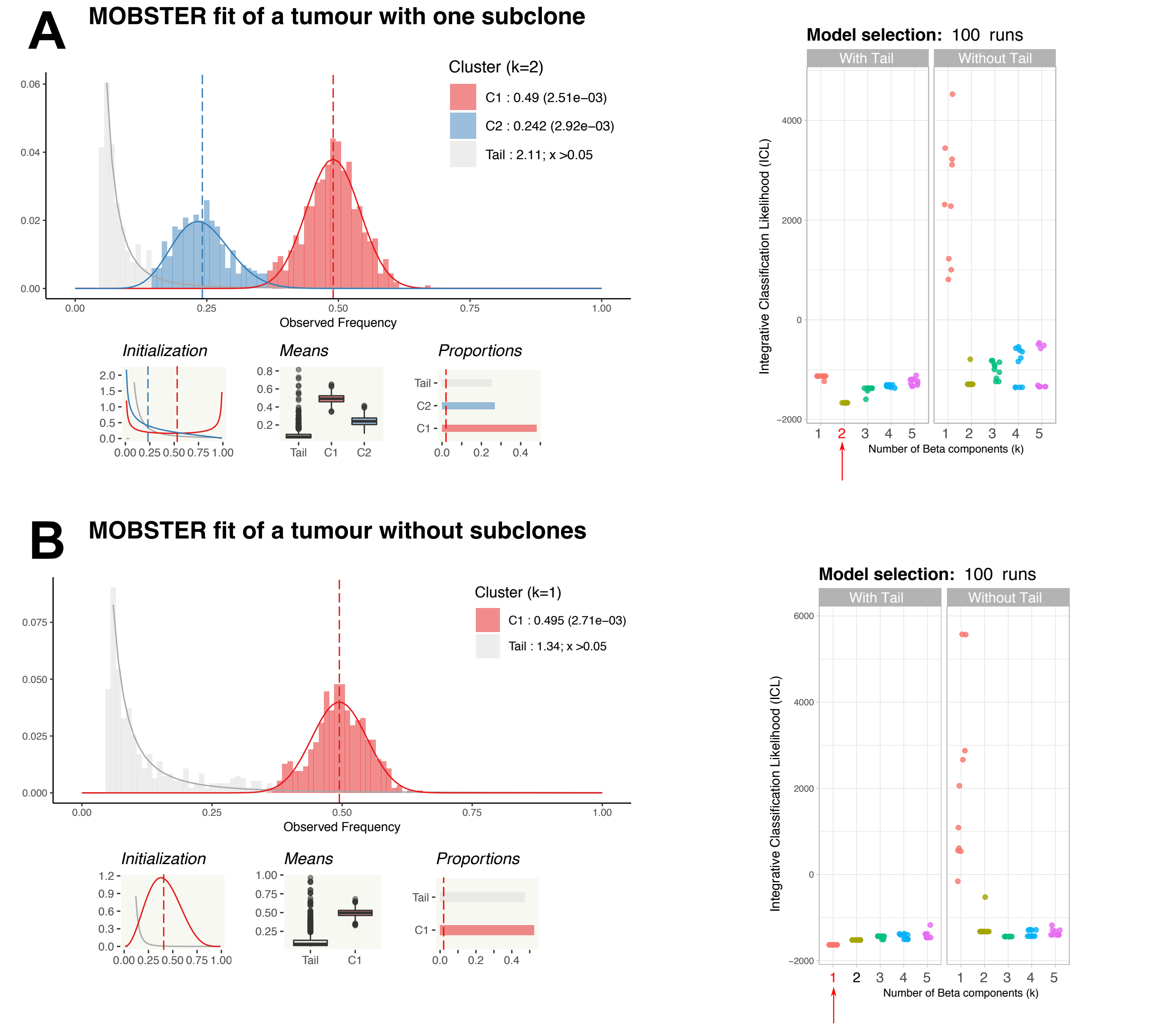

**Supplementary Figure S2. A.** Perfect fit for a simulated tumour with 1 subclone (coverage mean $120\times$, purity $1$, all mutations diploid). The top left histogram is the VAF data coloured according to hard clustering assignments; the tail is in grey and the components' parameters are reported in the caption. The smaller panels below the histogram show the initial condition of the fit computed by MOBSTER's peak detection routine, mean values for the distributions bootstrapped from posterior estimates of the fit and mixing proportions. Right boxplot are ICL scores for all the tested parameters, with and without tails during fit. Model-selection selects $k = 2$ for this data, and matches the ground truth. **B.** Successful fit of a neutral tumour (i.e., a tumour without subclones); plots are arranged as in panel A.

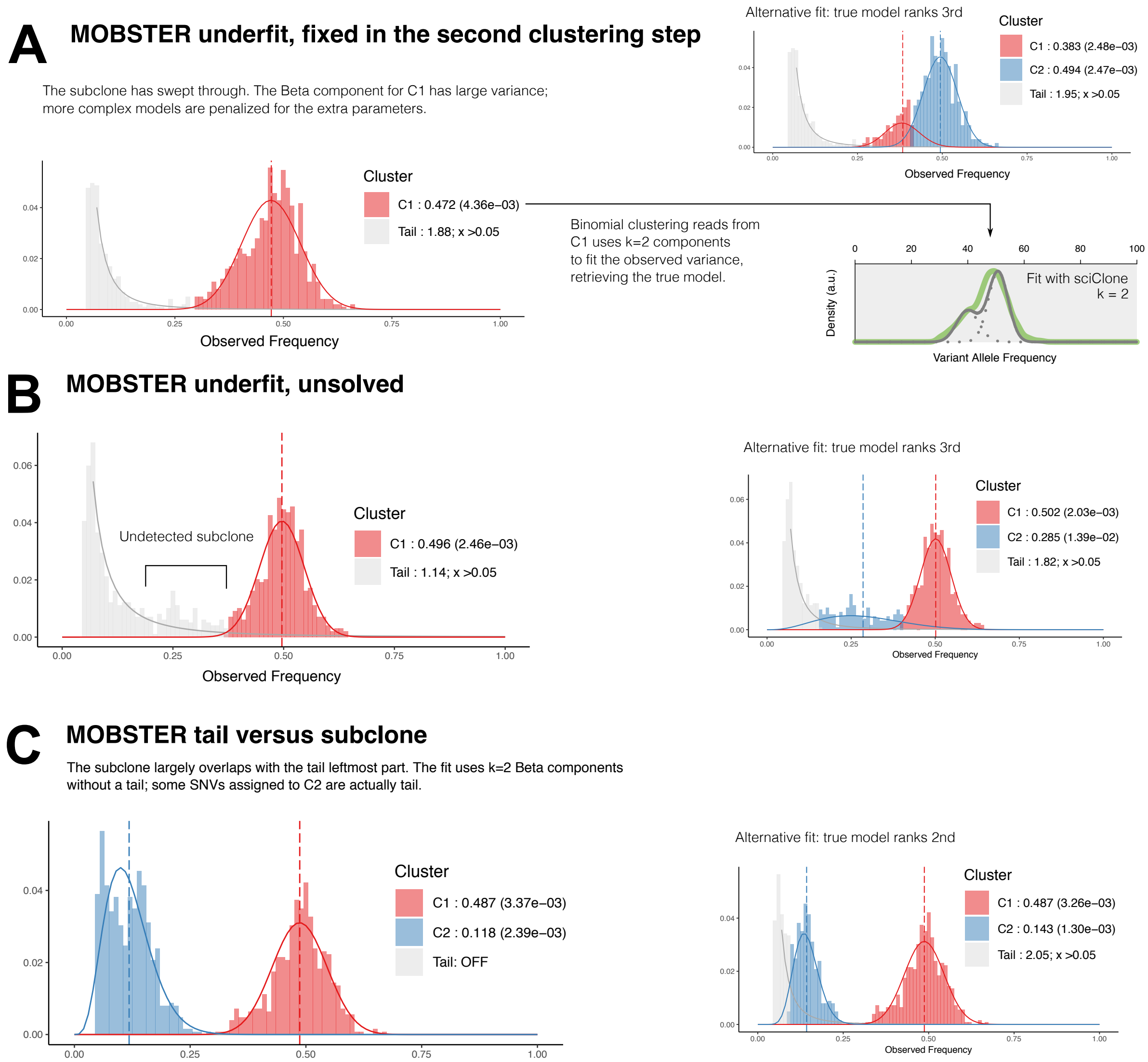

**Supplementary Figure S3.** **A.** Because Beta distributions can have large variance, the fit with MOBSTER might return $k = 1$ Beta components for a tumour where the subclone has almost totally swept through the ancestral population, misleading in this first step the true number of clusters. The true model is only ranked 3rd, in this case. The second step of clustering, however, uses Binomial distributions, which have a stricter model of the variance which depends only from the coverage. In this case, a fit with sciClone retrieves $k = 2$ Binomial components from read counts of the mutations in C1, which fixes the problem. **B.** Mistaken fit for a tumour with a small subclone with peak at $0.25$. For this VAF distribution MOBSTER prefers to use a tail; the true model is only ranked 3rd in terms of ICL score. **C.** Fit for a tumour with a very low-frequency large subclone. MOBSTER prefers a fit without a tail, and correctly places a Beta distribution on the subclone. This comes at the expenses of neglecting the leftmost part of a tail, whose SNVs get wrongly assigned to the subclone; the peak of the Beta component slightly deviates to the left, as due to the extra SNVs assigned. Other model selection scores like reICL can mitigate these situations and call both a tail and a subclone (see Supplementary Figure S7).

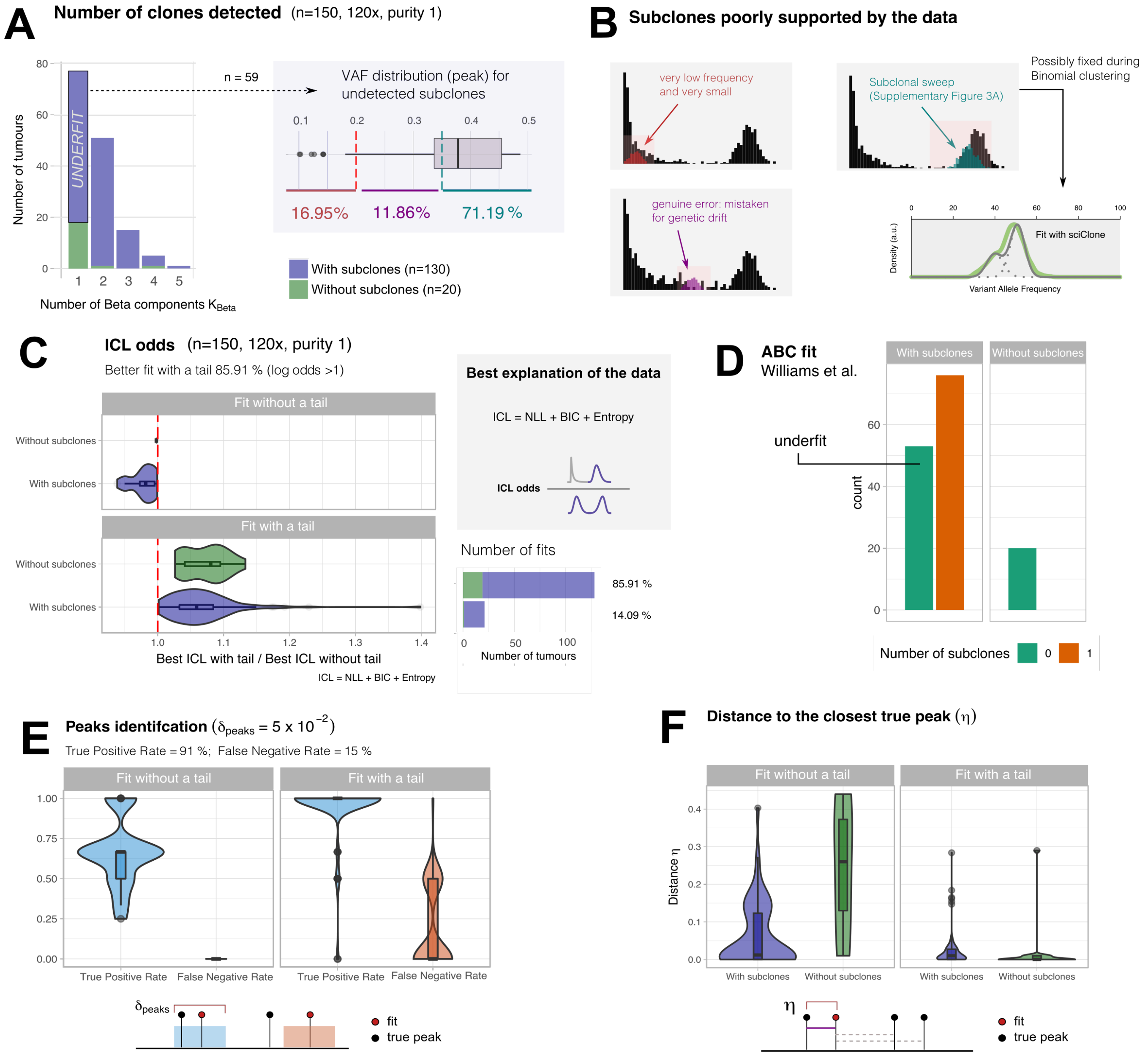

**Supplementary Figure S4. A,B.** Identification of the true number of clones in the data, per tumour type. In $59$ cases we do not detect the subclone in the data. We investigate this and find that in those cases input data harbours a weak signal of selection. Of $59$ cases, in $\sim 17\%$ $(10)$ a very low frequency subclone "hides" under a tail – this could be due to lo a small selective advantage compared to its ancestor. In $\sim 70\%$ $(40)$ cases a subclonal sweep is observed, MOBSTER uses one Beta component to model overlapping clusters – these cases might be resolved in a subsequent analysis (Supplementary Figure S3A). In the remaining cases $(9)$ MOBSTER genuinely misleads the subclone for genetic drift. **C.** We measured the confidence of the method towards its prediction. This is given by computing the ICL odds between the top model with, and without tail. Odds above 1 support tails: if MOBSTER fits a tail to data, the confidence is much higher than when i does not. This is a trend regardless the type of tumour which suggests that tails increases substantially the ability to describe the data. **D.** We also run the analysis with the ABC method described by (8), which uses a branching process simulator to fit the data. This method by construction will always fit a tail to the VAF distribution; in this tests we find underfit in 60 cases of tumours with a subclone where ABC fits a model without subclonal clusters. **E, F.** Given the outputs, we measured the precision of the fit as the rate of calling or missing a true peak (true positive, false negative). Peaks are matched with tolerance $\delta_{\text{peaks}} = 0.05$, and MOBSTER is very precise. We measure also the distance between the predicted fits and their closest true peak. Results show that tails reduce the error of the fit, consistently.

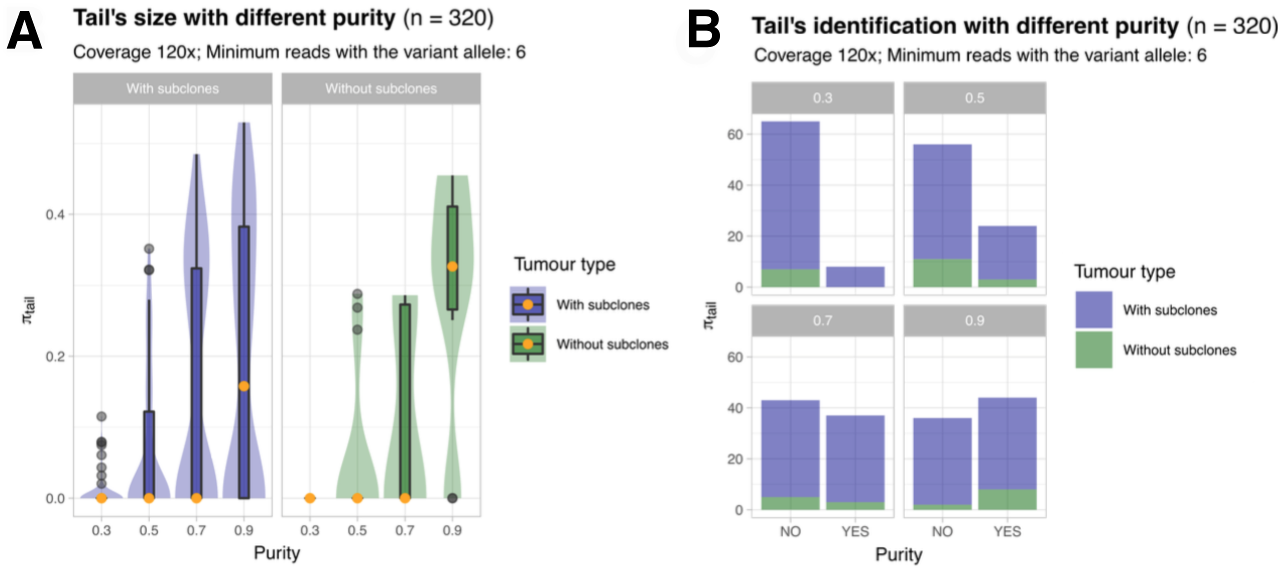

**C** **Summary**

| Parameter | mu | median | q = 0.025 | q = 0.975 |
|---|---|---|---|---|
| Mutation rate (mu) | 100 | 91.3890814 | 77.2711223 | 115.7983974 |
| Selection coefficient (s) | 100 | 0.6374105 | 0.2294687 | 1.8287272 |
| Subclone CCF | 100 | 0.5288401 | 0.4717509 | 0.5930701 |
| Age of the subclone (t) | 100 | 7.2281390 | 3.3465723 | 12.0527810 |
| Mutation rate (mu) | 20 | 18.5488965 | 14.6812041 | 21.8202356 |
| Selection coefficient (s) | 20 | 0.7599839 | 0.2522840 | 4.3417555 |
| Subclone CCF | 20 | 0.5334059 | 0.4574397 | 0.5891115 |
| Age of the subclone (t) | 20 | 8.0119639 | 3.8988010 | 15.8390315 |

High mutation rate

Low mutation rate

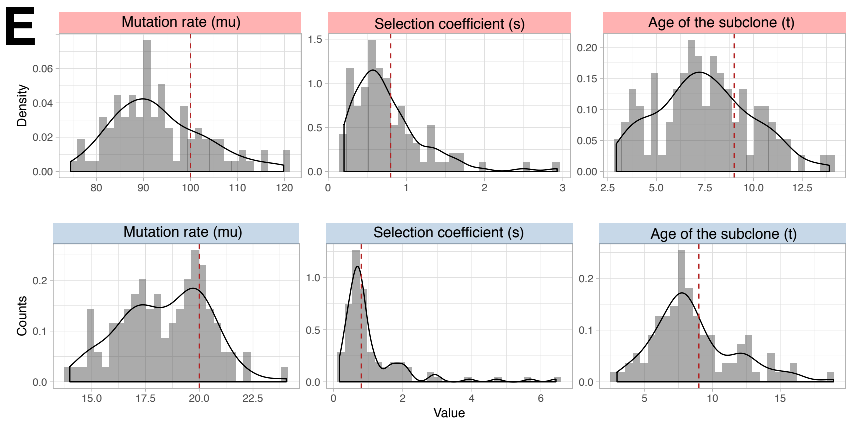

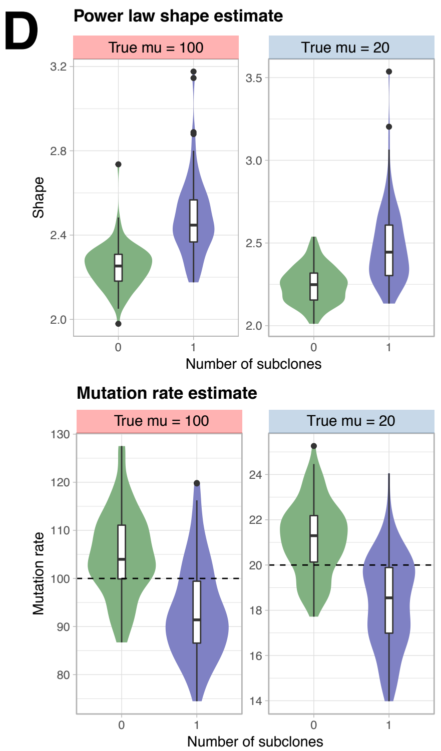

**Supplementary Figure S5. A.** We tested on $480$ tumours ($80$ per configuration) the ability of detecting tails as a function of purity spanning $0.3$, $0.5$, $0.7$ and $0.9$, assuming a coverage of $120\times$. The average tail size is reported (number of SNVs assigned to the tail in the fit). **B.** Barplot counts for the configurations of data and fit. **D,E,F.** We have simulated another batch of tumours ($n = 100$, with 0 or 1 subclone) fixing low or high mutation rate ($\mu = 20, 100$), run the fit and measured the evolutionary parameters that we can extract from MOBSTER's fits. Panel A reports summary statistics. Panel B shows boxplots of the power law shape estimate and the mutation rate (dashed line, ground truth), in tumour cell doublings. Panel C shows the histogram of the estimates of mutation rate, selection coefficient ($s$) and age of the subclone estimated; the simulated value (dashed line) is the line.

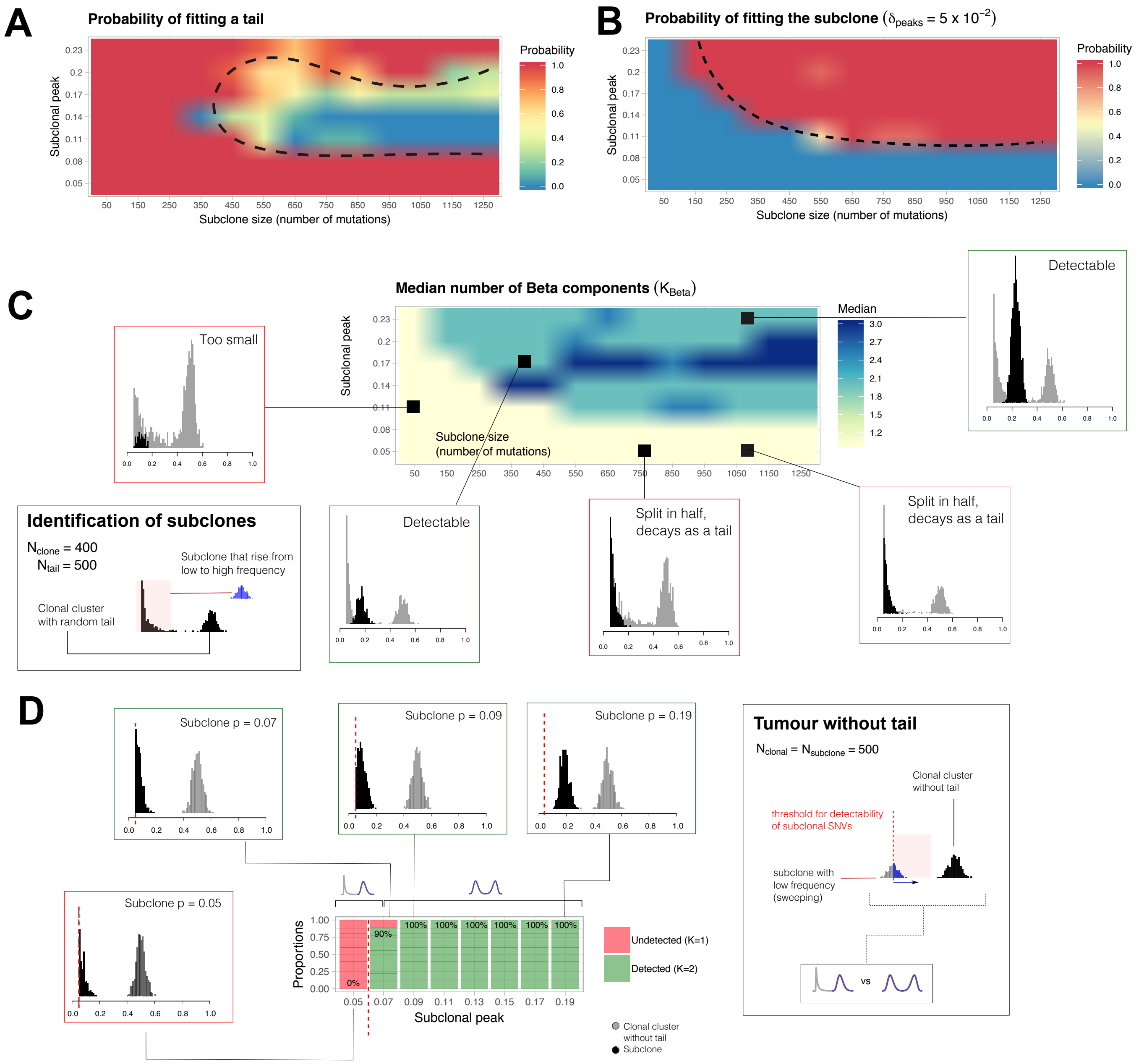

**Supplementary Figure S6. A,B.** Through a direct simulation of the frequency spectrum we assessed the ability of the method to call subclones, as a function of their size (number of SNVs), and peak. We simulated a clonal cluster with 400 SNVs (Beta $a = b = 100$, so $\mu = 0.5$) , and a tail with 500 SNVs (Pareto scale $x_* = 0.05$, shape randomly sampled in $[0.1, 3]$). We simulated a subclone with peak spanning from 0.05 to 0.23 (increases by $0.03$), and with a number of SNVs that span from $50$ to $1250$ (increases by $100$). We measure the probability of fitting a tail, and compare it to the probability of fitting the subclone with 10 independent runs per parameters configuration. Results show when subclonal detection is difficult because the subclone is either small, or at very low frequency. **C.** We measure the median number of Beta components fit by MOBSTER. In the boxes we annotate examples from this test. **D.** We used the frequency spectrum to assess the bias of the method towards calling tails. We simulated a low-frequency subclone and a clonal cluster without a tail, both with $500$ SNVs each. We considered a detectability limit on allelic frequency of $5\%$ $(0.05)$ and simulated the subclonal peak spanning from $0.05$ to $0.2$, with increases of a factor $0.02$. This mimics a subclone growing out of the frequency spectrum of the tail. In the panel we report the probability of detecting the subclone with a Beta (true model) against the probability of mistaking the subclone for a tail. Results show that the method is not biased, meaning that as soon as the subclonal distribution is evident (i.e., peak greater than $7\%$) the methods fits the subclone with a Beta, and the true model is correctly retrieved. The subclone is mistaken when its distribution is peaked exactly at the detectability limit of allelic frequency. See also the examples plot from this test.

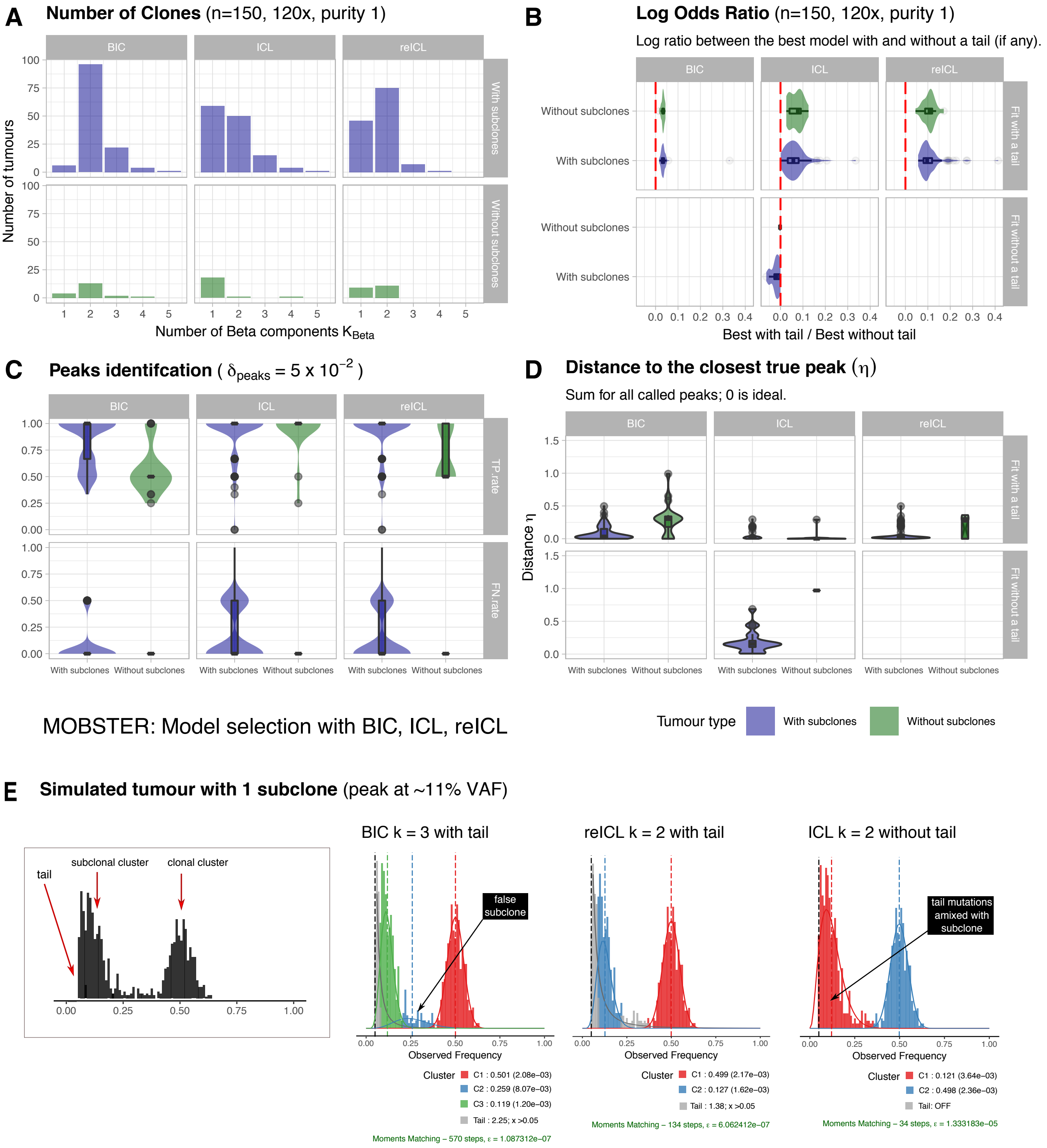

**Supplementary Figure S7. A.** For the same synthetic tests described in the Main Text, we show the number of clones detected by performing model selection with BIC, ICL and reICL. **B,C,D.** Similarly, we plot the same measures of precision and fit confidence for the best models obtained with those different criteria of model selection. **E.** Example different fits for a tumour with 1 subclone. We can see how BIC calls multiple clones, plus a tail. reICL retrieves the true model and ICL drops a tail in favour of a subclonal component. This is due to the different entropy terms used in ICL, against reICL; BIC does not use entropy at all.

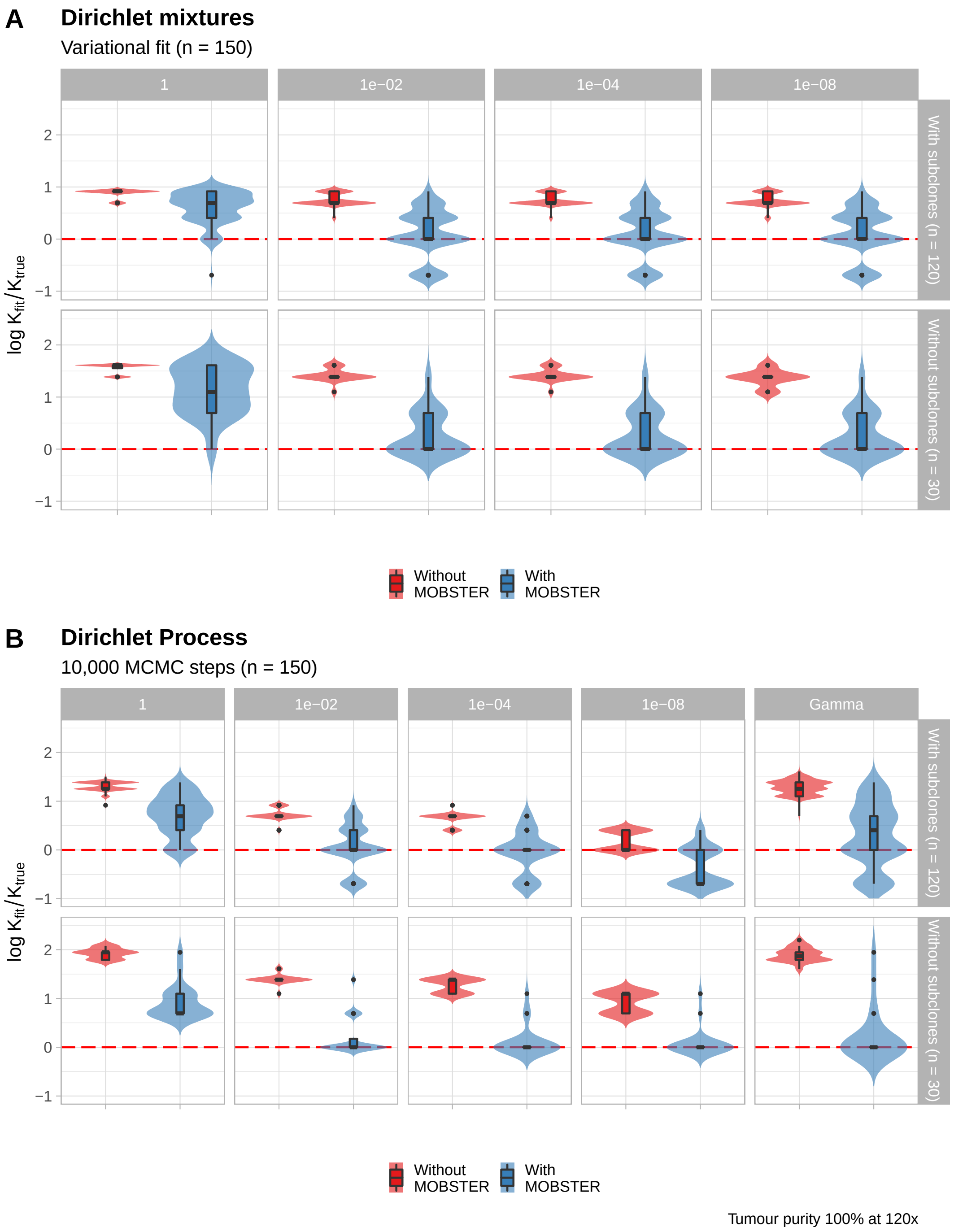

**Supplementary Figure S8. A.** Extended results presented in the Main Text with values of the concentration parameter $\alpha$ that span over $1, 10^{-2}, 10^{-4}$ and $10^{-8}$, for the variational inference fit of a Dirichlet mixture. **B.** The analogous results for a Dirichlet Process implementation, where we also test the Gamma prior on $\alpha$.

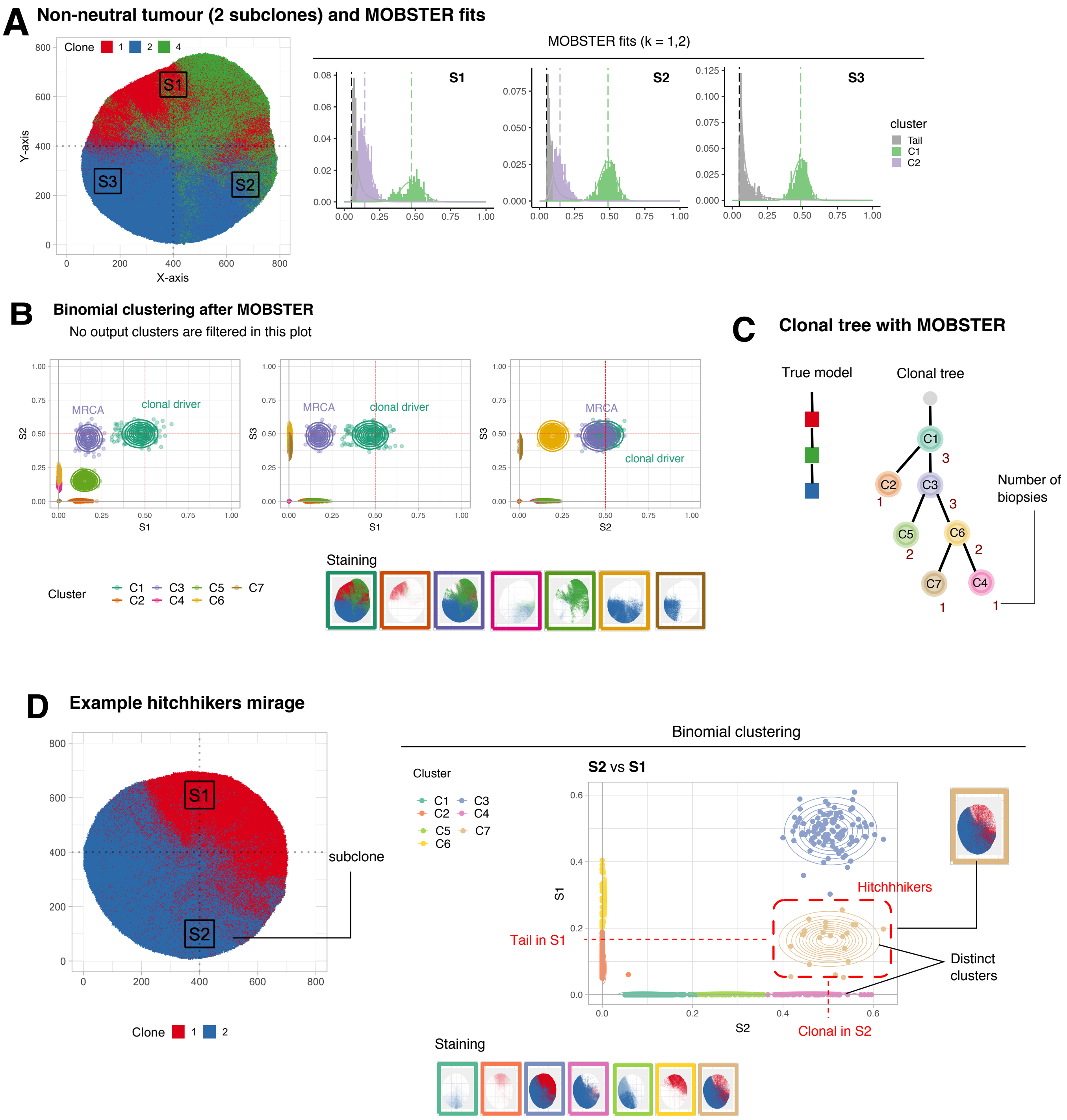

**Supplementary Figure S9.** **A.** As in the Main Text, we show a simulated 2D tumour with $2$ subclones, for which we sample 3 biopsies. This sampling is tricky, as 2 biopsies (S1 and S2) are boundary and thus contain an admixed signal. The fit for all biopsies with MOBSTER is perfect, and the clones are properly detected with 2, 2 and 1 Beta components. **B.** We run the variational fit for multivariate Binomial mixtures available in MOBSTER on the read counts of non-tail mutations. In bottom we show the virtual staining of the simulated tumour, which shows the presence of clear most recent common ancestors among the clusters. **C.** We manually curate a clonal tree from the virtual staining in panel B, and annotate the number of times each cluster occur in the 3 biopsy samples. **D.** Example hitchhiker mirage for the type of tumour discussed in the Main Text. Cluster C7 contains mutations that in sample S1 are tail, while in S4 they are clonal because they hitchhiked to the subclonal driver which originated the blue subclone.

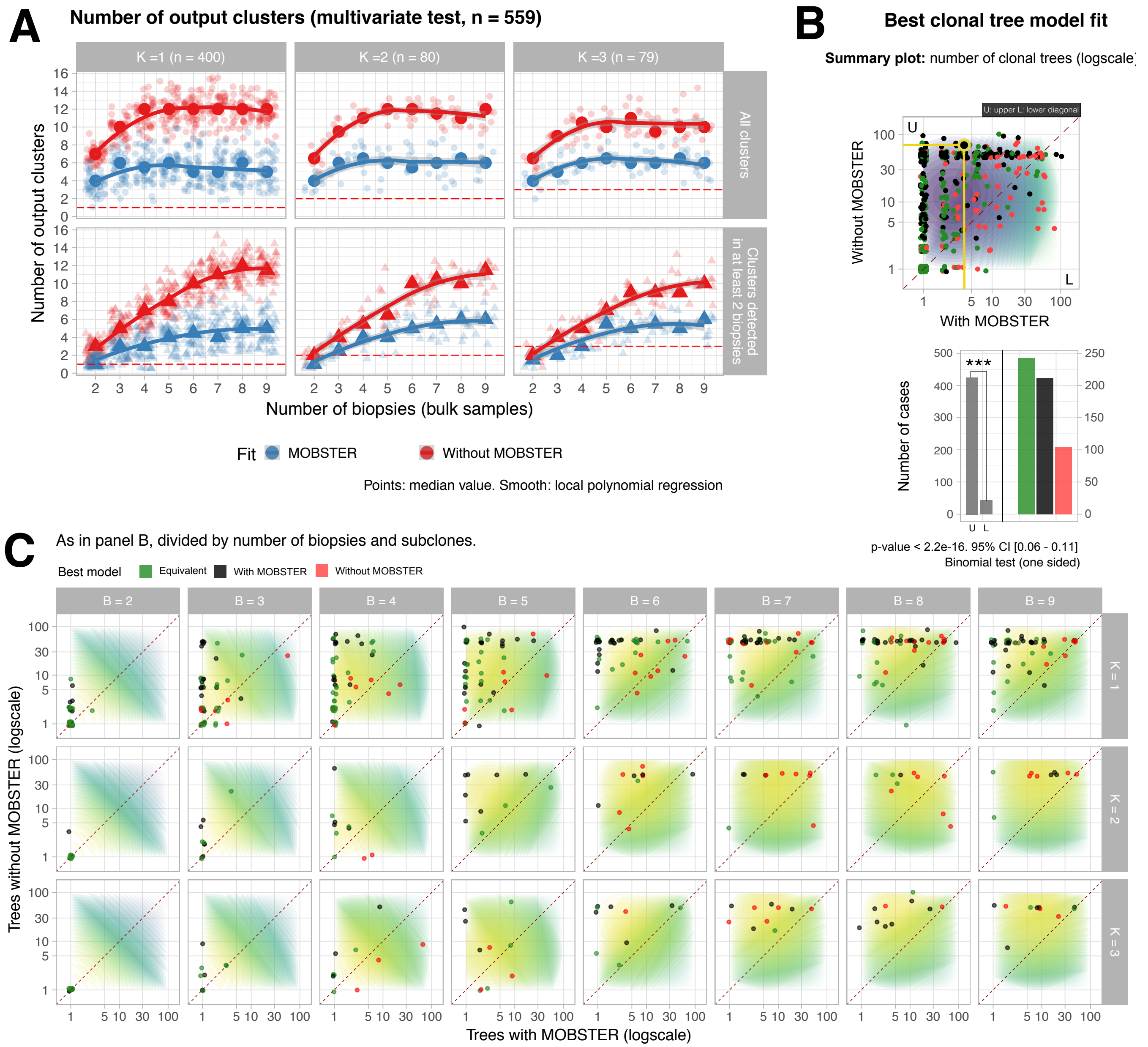

**Supplementary Figure S10.** We expand the results shown in the Main Text, Figure 5. **A.** For variable number of overall clones $k$ ($k-1$ subclones) we report the number of clones called with and without MOBSTER, estimated via variational Binomial mixtures as in Supplementary Figure S9. The red dashed line shows the actual number of true clones under positive selection in the tumour; we compare the overall number of clusters, and the reduced count where we remove those that occur only in a single biopsy. **B.** The sample plot of Figure 5 Main Text, with each point coloured to show if the best model is obtained with MOBSTER (black), without (red) or equivalently (green). The bottom barplots shows the number nuimber of points in the upper and lower diagonals of the plot. **C.** We plot the same counts for the number of detected clonal trees with and without MOBSTER, divided by number of subclones in the simulation, and number of collected biopsies (2 to 9). Results show that the detection of more points in the upper diagonal, which means fewer trees available with MOBSTER, is systematic across all simulated configurations of tumour and sampling.

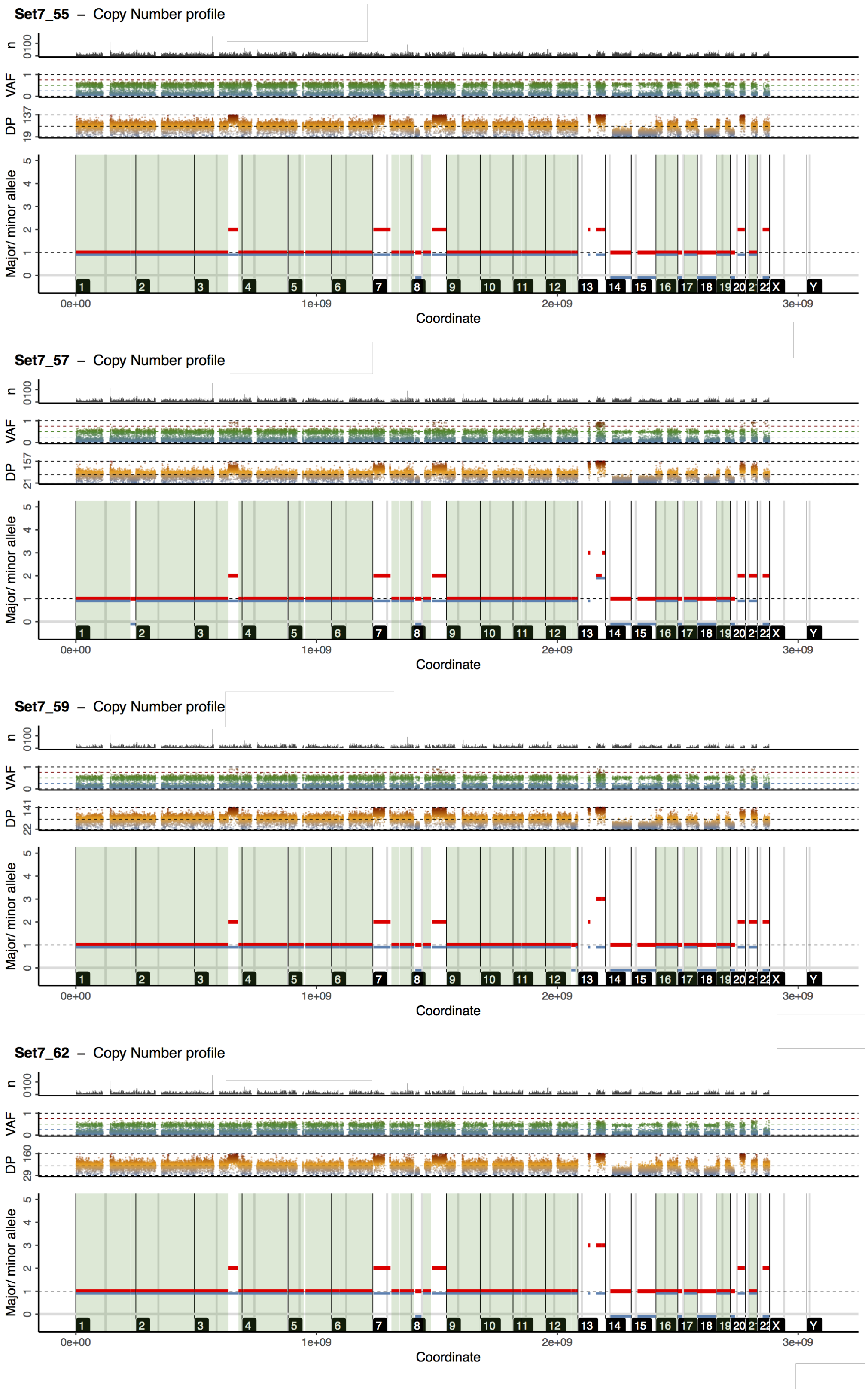

**Supplementary Figure S11.** CloneHD Copy Number calls for the `Set07` colorectal carcinoma. For each sample, we show in the bottom panel the major and minor copy number estimate across the whole genome. We show on top of that a track with mutation counts (binned every megabase, filtered 10 megabases around the centromere), the VAF value adjusted for copy number state and purity of the sample (in $[0, 1]$), and the observed depth (quantiles $[0.01, 0.99]$ of the distribution). We shadow diploid regions across all four samples (minor = Major = 1) in green.

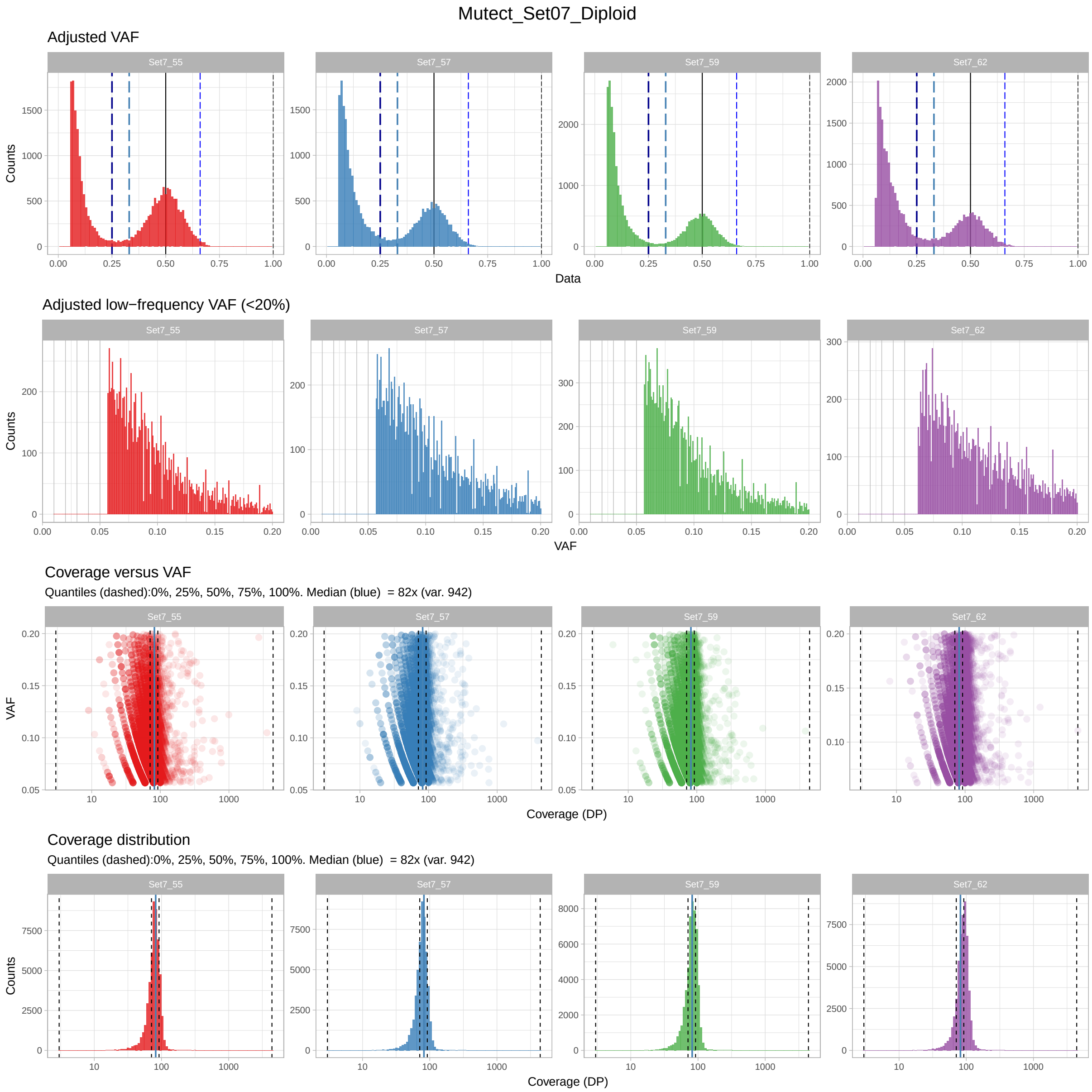

**Supplementary Figure S12.** SNVs called with Mutect for the `Set07` colorectal carcinoma, and their VAF distributions adjusted for copy number state and purity of each sample (0.88, 0.88, 0.88 and 0.80). The top row shows the full VAF distribution (binsize $0.01$), below we zoom in the interval of VAF below $0.20$ (binsize $0.001$). The third row shows the VAF against the depth value (DP), with logscale on the x-axis, and the last row shows the distribution of sequencing depth with quantiles and median annotated.

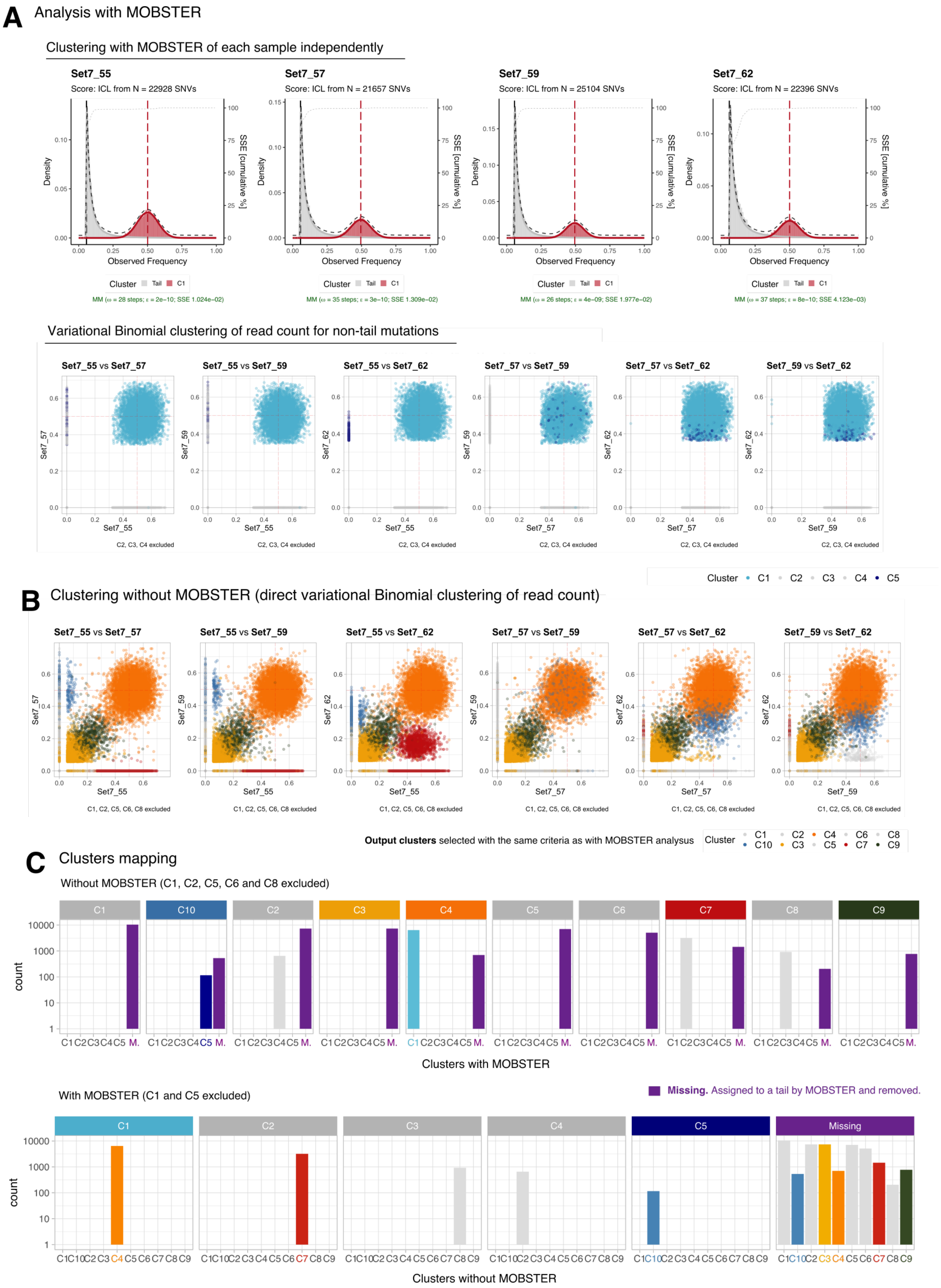

**Supplementary Figure S13. A.** Clusters obtained with MOBSTER and the analysis proposed show that, with the current data of this tumour, there is no evidence of subclonal selection. The top row are the fits computed from running MOBSTER independently on each of the available tumour samples, the bottom row is the result of the variational multivariate Binomial method available in MOBSTER, applied to non-tail mutations. Output clusters in gray are found in only one tumour biopsy (private mutations), and removed. There are two clusters of mutations shared across patients: the clonal cluster and cluster C5 (darkblue), which however accounts only for <1% of the overall number of mutations of this tumour and, for this reason, we reject also this cluster and conclude that this tumour is of monoclonal origin. **B** Variational multivariate Binomial clustering of all tumour mutations – i.e., without MOBSTER – shows a completely different evolution, with $k = 5$ clusters when we use the same criteria used in panel A to select output clusters. **C.** The two clustering analysis predict very different clonal compositions of this tumour. We show the the reciprocal mapping among clusters computed with and without MOBSTER (top and bottom plots). This visualisation shows that all the mutations assigned to cluster C3 by a standard analysis are clustered into the tails that we can retrieve with MOBSTER, and therefore removed from downstream read counts clustering. We also observe that the signal which dominates several subclonal clusters without MOBSTER is entirely driven by tail mutations (purple bars). For this reason, we can conclude that the tumour composition retrieved with these analyses are totally different.

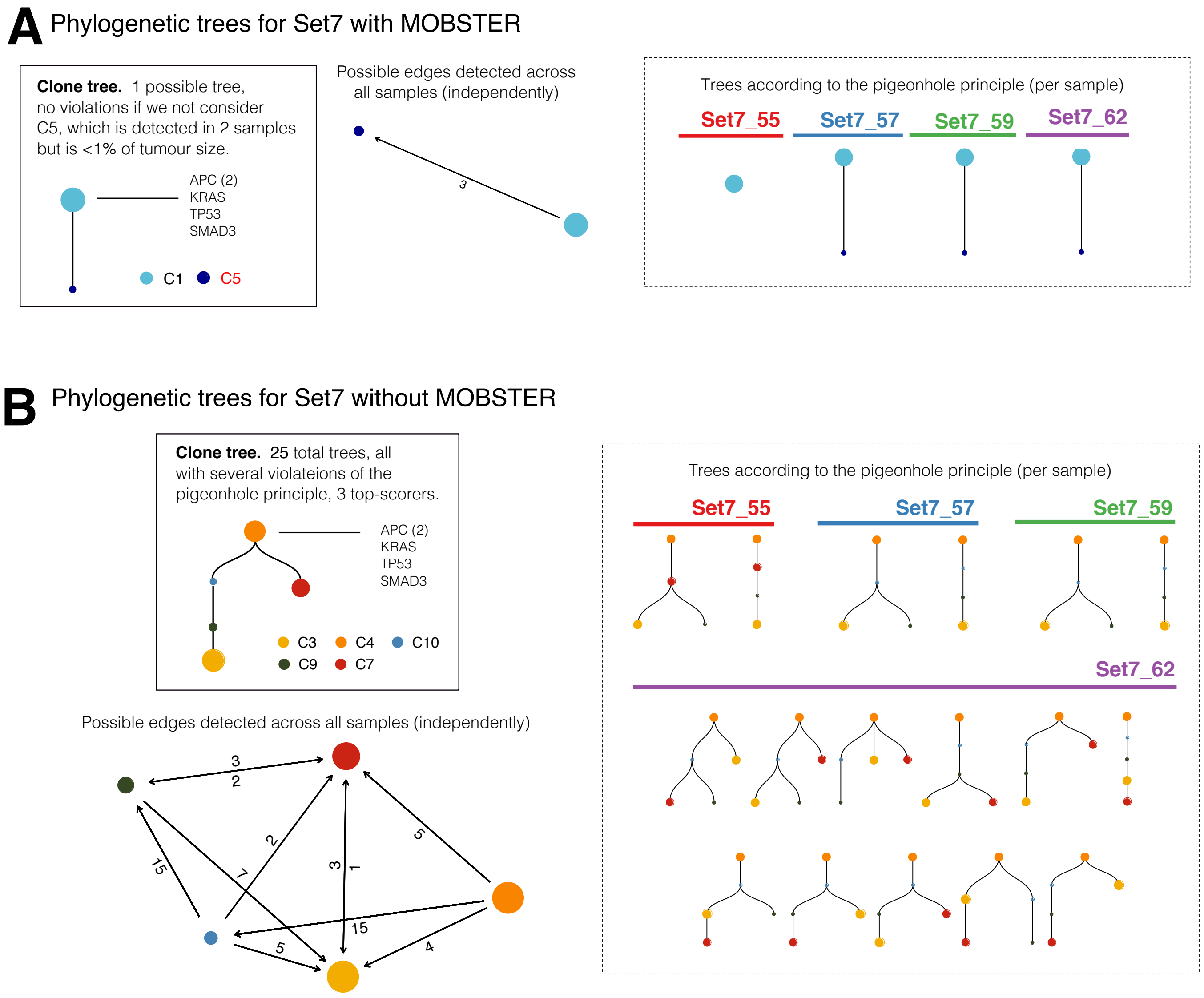

**Supplementary Figure S14. A.** Clone trees that we can compute from the output clusters with MOBSTER. We show the result of computing all possible model fits per sequenced sample, and compute the union graph where each edge reports the number of times a certain edge among tree nodes is detected. From that, we use as a generator of all possible trees that can fit all samples at once, and a previously described method (9) to compute the number of violations of the pigeonhole principle for each tree. We find only 1 tree that can be fit to all the regions of this tumour. **B.** We do the same procedure with the output clusters that we can compute with a standard analysis without MOBSTER. This shows a much more complex clone tree for this tumour, with consequent higher uncertainty propagated downstream this analysis.

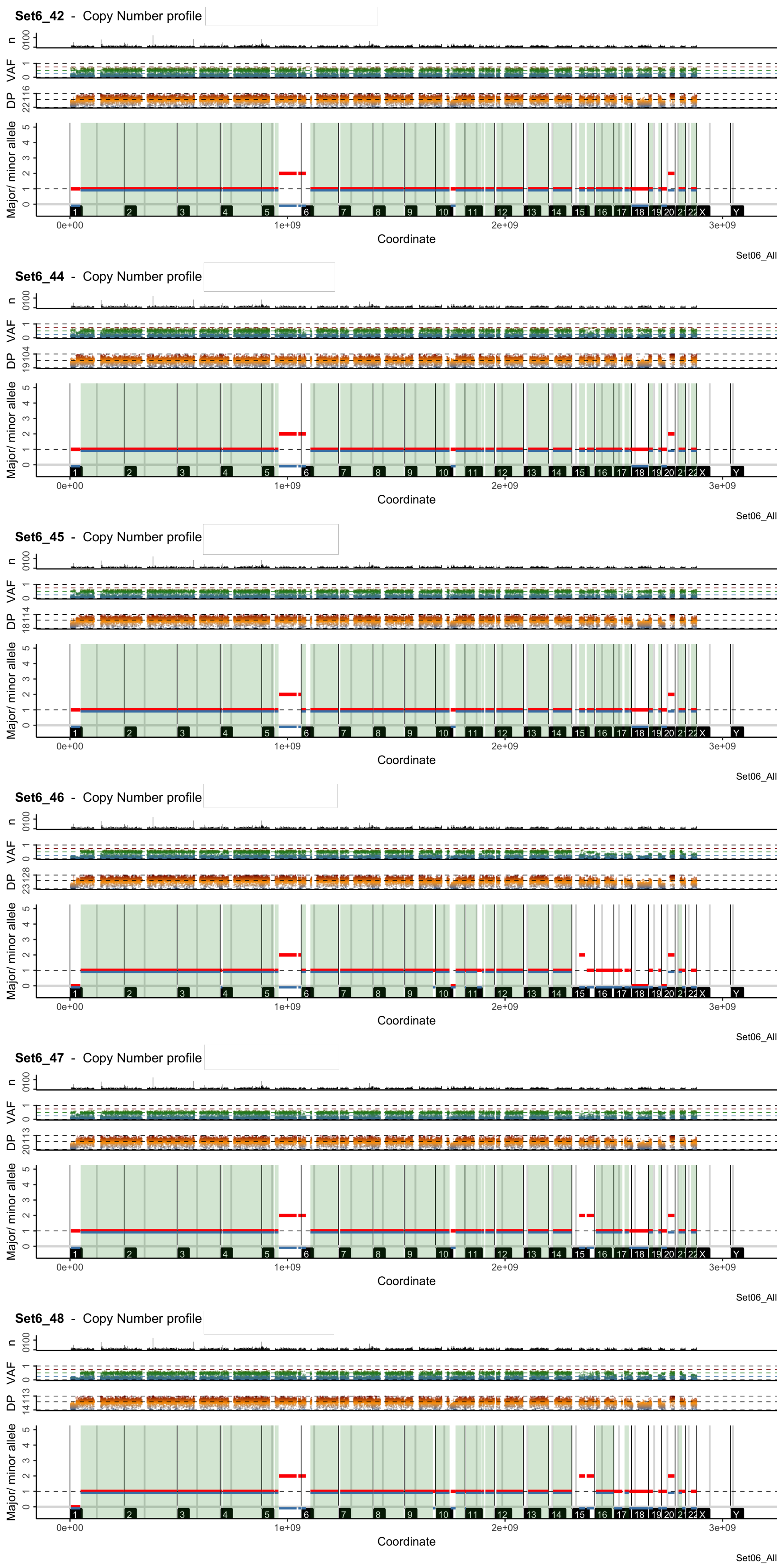

**Supplementary Figure S15.** CloneHD Copy Number calls for the `Set06` colorectal carcinoma, with the same notation of Supplementary Figure S11.

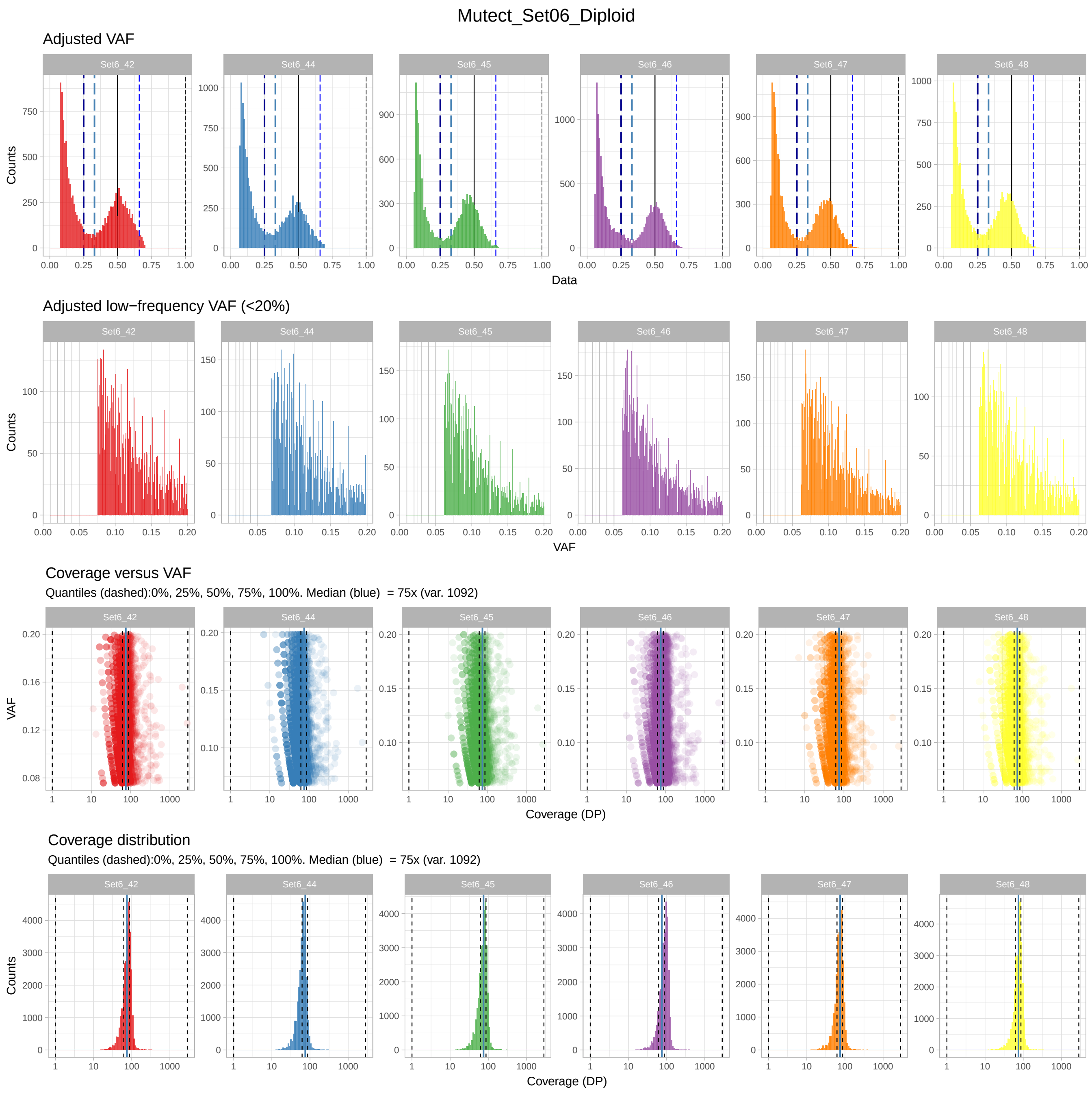

**Supplementary Figure S16.** SNVs called with Mutect for the `Set06` colorectal carcinoma, with the same notation of Supplementary Figure S12. The estimated tumour purity of the samples are 0.66, 0.72, 0.80, 0.80, 0.80, and 0.80.

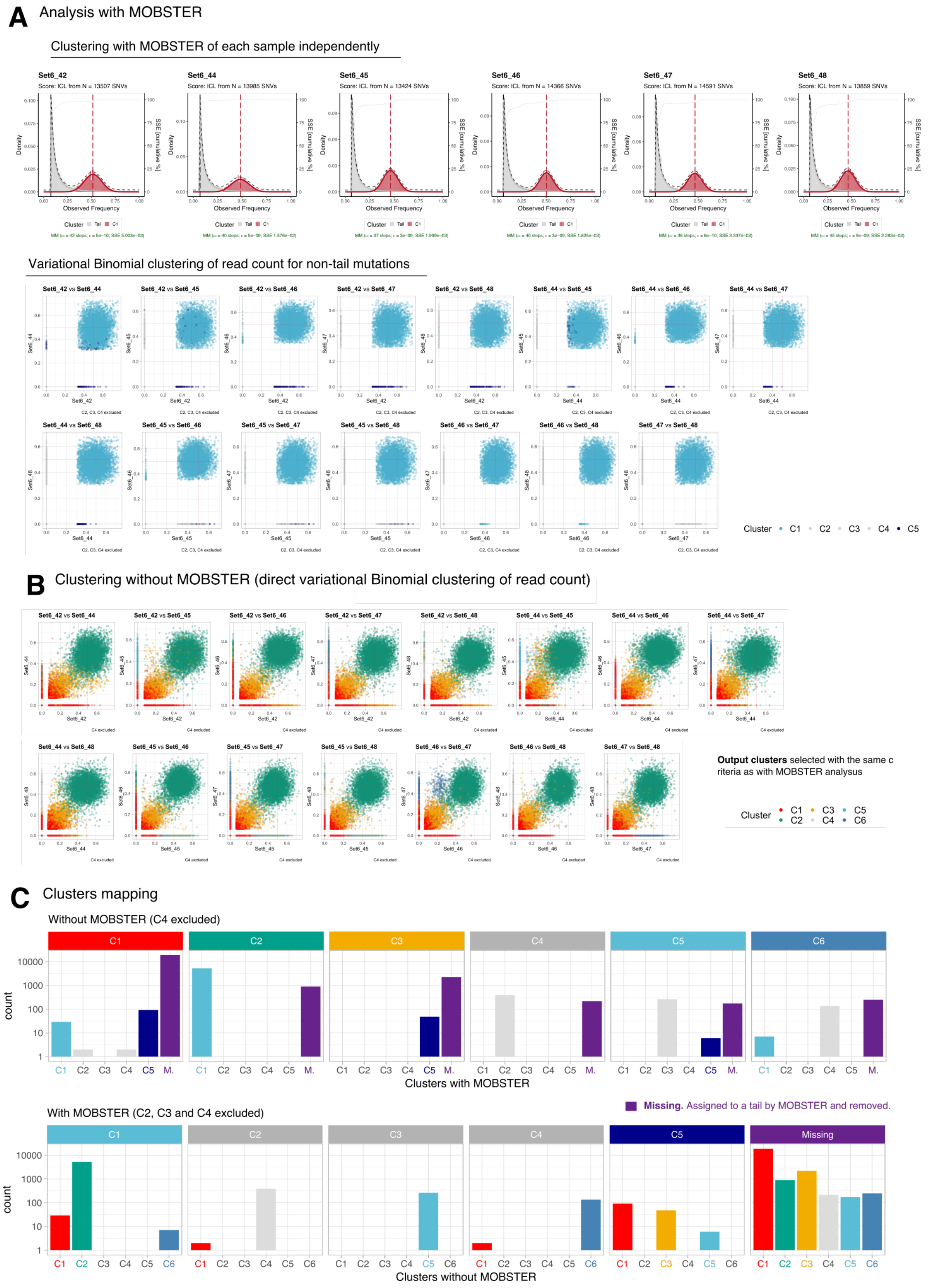

**Supplementary Figure S17.** With the same notation and procedure described in Supplementary Figure S13 we show the analysis of this tumour with MOBSTER (A), with a standard analysis (B) and we compare the clustering assignments (C). As for `Set06` the results of these analyses show a very different clonal architecture when we use MOBSTER to control for tails.

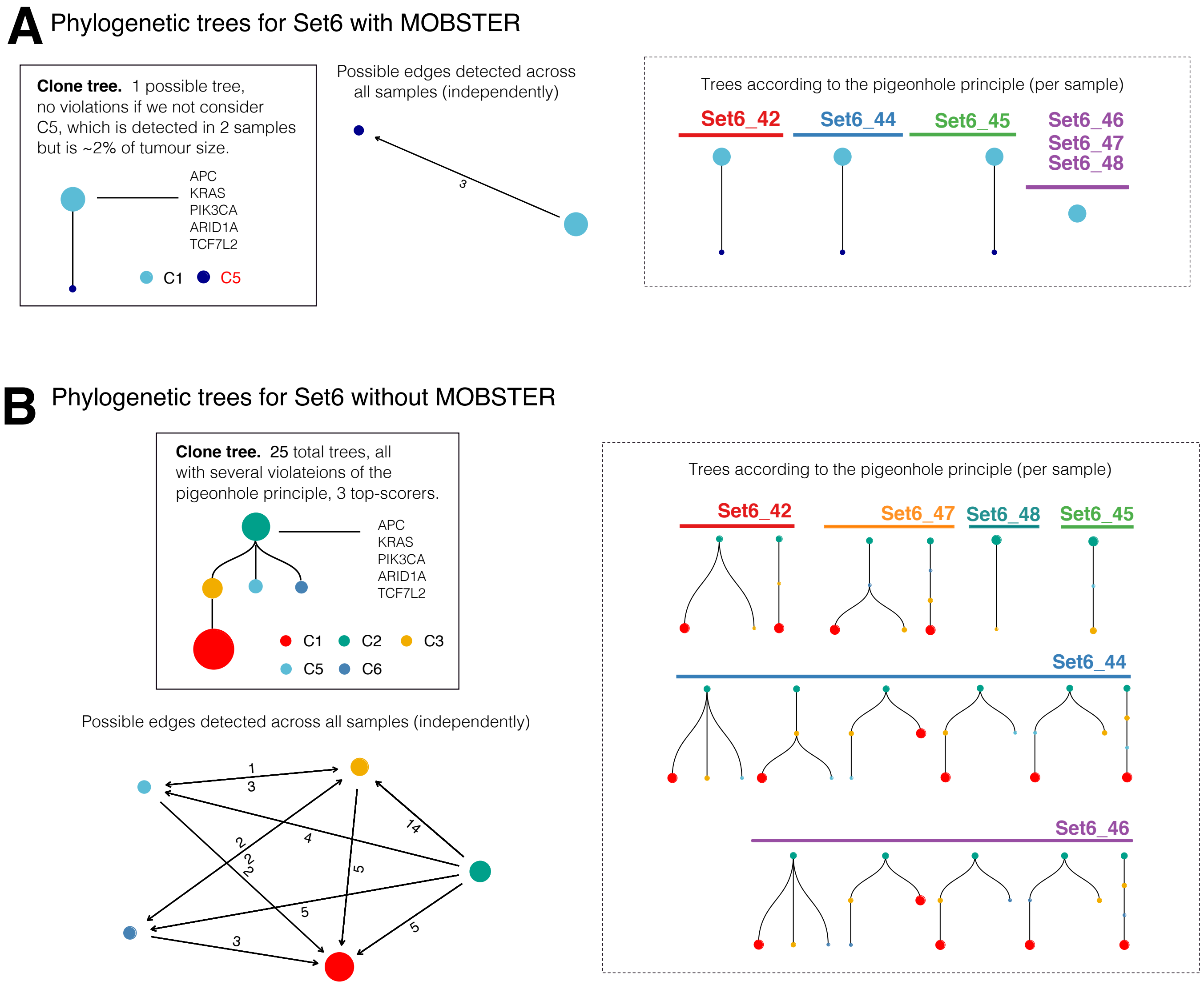

**Supplementary Figure S18.** We used the same procedure described in Supplementary Figure S14 to compute the clone trees for `Set06`. The results are very similar to the one computed for the other tumour, and highlight the difference in the prediction of a monoclonal tumour, with MOBSTER, compared to an estimated excess of subclonal selection which is heavily confounded by tail mutations in a standard analysis.

**Additional data table S1 (Supplementary_Table_S1_MSeq_calls)**

Whole Genome Sequencing data for MSeq patients Set06 and Set07.
